# Increased risk of many early-life diseases after surgical removal of adenoids and tonsils in childhood

**DOI:** 10.1101/158691

**Authors:** Sean G. Byars, Stephen C. Stearns, Jacobus J. Boomsma

## Abstract

**BACKGROUND:** Surgical removal of the adenoids and tonsils are common pediatric procedures, with conventional wisdom suggesting their absence has little impact on health or disease. However, little is known about long-term health consequences beyond the perioperative risks. Such ignorance is significant, for these lymphatic organs play important roles in both the development and the function of the immune system.

**METHODS:** We tested the long-term consequences of surgery in the population of Denmark by examining risk for 28 diseases with ̴1 million individuals followed from birth up to 30 years of age depending on whether any of three common surgeries (adenoidectomy, tonsillectomy, adenotonsillectomy) occurred in the first 9 years of life. To weigh costs and benefits, we also compared the absolute risks for these diseases to the risks for the conditions that these surgeries aimed to treat. We obtained robust results by using stratified Cox regressions with statistically well-powered samples of cases (with surgery) and controls (without surgery) whose general health was no different prior to surgery. We adjusted our estimates of risk for diseases occurring before surgery, stratified for sex (and other effects) and for 18 covariates, including parental disease history and birth metrics.

**RESULTS:** We found significantly elevated relative risks for many diseases, with effects on respiratory, allergic and infectious disorders after removal of adenoids and tonsils being most pronounced. For some of these diseases, absolute risk increases were considerable. In comparison, many risks for conditions that surgeries aimed to treat were either not significantly different or significantly higher following surgery up to 30 years of age. This suggests that any immediate benefits of these surgeries may not continue longer-term, while resulting in slightly compromised early adult health due to significantly increased risk of many non-target diseases.

**CONCLUSIONS:** Our results indicate that surgical removal of tonsils and adenoids early in life are associated with longer-term health risks. They underline the importance of these organs and tissues for normal immune functioning and early immune development, and suggest that these longer-term disease risks may outweigh the short-term benefits of these surgeries.

## Introduction

Removal of adenoids and tonsils are common surgical procedures in childhood [1–4] with conventional wisdom suggesting their absence has little negative longterm impact on health [3], yet not much support for this assertion is available beyond assessments of the short-term perioperative risks. Establishing the longer-term impact of these surgeries is critical because the adenoids and tonsils are parts of the larger immune system [3, 5], have known roles in early pathogen detection and defense [3, 5], and are most commonly removed during ages that are most sensitive for early immune system development. Some recent single-disease studies have shown subtle shorter-term changes in risk after surgery [6-9], but no longer-term assessments of risk across a broad range of diseases is yet available. Here we analyzed the long-term health outcomes post-surgery by examining risk for 28 diseases in ̴1 million individuals who were followed from birth to up to 30 years of age depending on whether adenoidectomy, tonsillectomy or adenotonsillectomy occurred during the first nine years of life.

Historically, the functions of tonsils and adenoids have intrigued scientists and medical professionals alike [10, 11]. Current research suggests that these tissues play specialized roles in immune development and function. For example, the tonsils help protect directly against pathogens [3, 5] and indirectly by stimulating further immune responses via communication with the rest of the lymphatic system [3, 5, 12]. The three main types of tonsils (pharyngeal, palatine, lingual) surround the apex of the respiratory and digestive tract forming ‘Waldeyer’s ring’, which provides early warnings for ingested or inhaled pathogens [3, 5, 12]. Evidence from other fields is beginning to suggest that altered early-life immune pathways (including dysbiosis, [13]) can have lasting effects on adult health, warranting concern that the surgical removal of adenoids and tonsils in childhood may affect later health in ways that have been underappreciated.

Surgeons often remove adenoids and tonsils to treat common childhood illnesses (e.g. recurrent tonsillitis, middle ear infections). Research on the consequences of these surgeries primarily relates to perioperative risks [3, 14] and the ensuing short-term consequences of surgical procedures for the symptoms to be treated. That tonsils (particularly the adenoids) tend to atrophy with age, being largest in childhood and absent in adulthood [1], has suggested that their absence would not affect adult health [3]. However, this does not exclude the possibility that these lymphoid tissues are most immunologically active in early life because they have a critical role in normal immune system development [3, 5]. In that perspective the potential long-term risks of these surgeries deserves to be assessed in light of an extensive general literature about perturbations to early growth and development that may alter risk of many adult diseases (e.g. [15, 16]).

Recent work provides support for such analyses. Except for rhinosinusitis, ear and throat infections [17, 18], and sleep apnea [19] there has been little work on the health consequences of removing the adenoids or tonsils in childhood. Evidence for consequences of adenoidectomy for the risk of asthma is mixed [9].

Other studies indicated that tonsillectomy does not necessarily reduce the risk of respiratory diseases in adults, but there have been hints that it may increase the risk of chronic disease, including inflammatory bowel disease [8], and improvements in sleep apnea of children appear to be less than hoped [19]. Some work suggests that surgery changes the risk of non-respiratory diseases, with tonsillectomy being associated with significantly increased risk of breast cancer [6] and premature acute myocardial infarctions [7]. The benefits of tonsillectomy for kidney disease are only realized for a very small sample [20]. These single-disease studies make it clear that more comprehensive assessments of long-term health risks of childhood surgeries could be illuminating.

In our present large-scale study, we estimated disease risk depending on whether adenoids or tonsils were removed in the first nine years of life. In contrast to previous single-disease, single-surgery studies that typically focus on short-term risks, we:

1. examined effects of all such surgeries at the younger ages that comprise the most sensitive period for early-life immune development, which coincides with the time-window in which these surgeries are most commonly performed (i.e. generally [1, 21] and in Denmark, see Fig. 1);
2. calculated long-term risks up to 30 years of age for 28 different diseases belonging to 14 generally acknowledged disease groups;
3. compared relative and absolute risks and number (of patients) needed to treat (to obtain a first case of harm) to adjust risk metrics for naturally occurring background rates of disease and to translate risk statistics into numbers that are clinically applicable;
4. compared long-term post-surgical absolute risks and benefits for diseases and conditions that these surgeries aim to treat; and
5. tested for general health differences between cases and controls prior to surgery to establish that individuals who underwent surgery were not sicklier on average than the control group.

**Figure 1.**
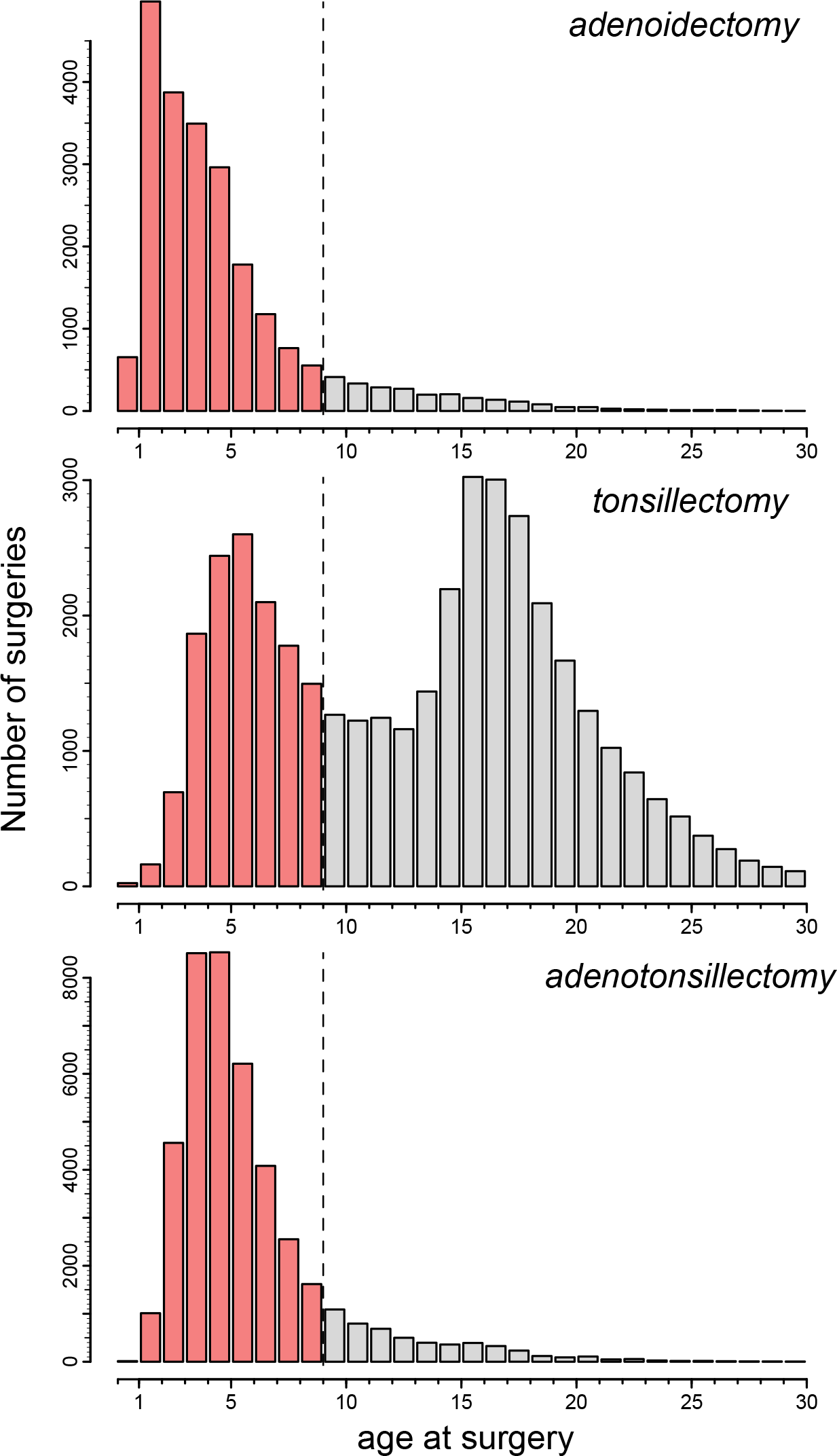
Age at adenoidectomy, tonsillectomy and adenotonsillectomy for 1,753,100 Danes born between 1978-2009 and the selected surgery observation window of nine years. This cut-off for inclusion as surgery cases (red bars] was deemed optimal because the first decade of life is critical for normal immune system development, it represents most of the period in which these surgeries are usually performed, and maximized the number of years available for disease follow-up, post-surgery. For tonsillectomy this meant that we ignored a second peak at ̴ 16-17 years, for inclusion of these surgeries would have implied insufficient time for follow-up (to 30 years of age]. Our study thus explores the impact of the three types of surgery when performed during childhood rather than adolescence. Individuals with these surgeries beyond the 9-year observation end-point (dashed vertical line] were not included as either cases or controls. Individuals were also excluded if they had multiple surgeries at different ages, i.e. some individuals underwent adenoidectomy followed by tonsillectomy years later or vice versa. Such cases were rare in the sample (<0.2%].

## Materials and Methods

### Study sample obtained from the Danish health registries

We utilized health data on ̴1 million individuals born (as singleton live births) between January 1978 and January 2009 from the Danish Birth Registry. This included healthy cases and controls who were not diagnosed with focal diseases during the surgery observation period (i.e. from birth to 9 years of age) and who had no surgery performed during the follow-up period from 9 to maximally 30 years of age; see Table s1 for sample sizes). This sample design ensured sufficient statistical power to detect differences in disease risk between cases and controls while avoiding obvious confounding effects that could suggest reverse causality. Figure 1 provides details on how we optimized observation and follow-up ages. We did not include individuals with missing covariate data or values outside typical ranges for birth weight (i.e. 1850-5400 grams), gestation length (30-42 weeks), paternal age (15-60 yrs), and maternal age (15-46 yrs) at birth, those who were living during (but not born between) 1978 and 2009 because their health records were incomplete (see Supplementary Methods - study sample, for further details).

Data on covariates were obtained from the Danish Birth Registry (i.e. pregnancy, birth and pedigree information) and other registries including: Danish Patient Registry holding nation-wide hospital admission and ICD diagnosis data; Danish Psychiatric Registry with psychiatric diagnoses for inpatient admissions; Danish Civil Registration and Cause of Death Registries containing date of death, migration, socioeconomic and other demographic information. We combined individual-level information from different registries using unique personal identification numbers (de-identified from central-person register CPR numbers). Because the health system in Denmark is largely free for all residents and public health data are continuously and rigorously collected for every resident in Denmark thanks to the CPR number assigned from birth, we are confident that we obtained the complete health and socioeconomic histories of the ̴1 million individuals in our data set. This allowed us to employ rigorous controls and validations (see below).

### Defining surgery groups

There are three main types of tonsils: pharyngeal tonsils (hereafter referred to as the adenoids), palatine tonsils (hereafter referred to as the tonsils) and lingual tonsils. Adenoids are located towards the posterior wall of the nasopharynx, tonsils are located along the anterolateral walls of the oropharynx, and lingual tonsils are located at the base of the tongue. We focused on surgery removing the first two (adenoidectomy, tonsillectomy), for the lingual tonsils are not commonly removed. We included adenotonsillectomy where adenoids and tonsils are removed in the same surgery.

Surgery codes are based on early ICD operation classification codes from Danmarks Statistics (up to 1996) and the later Nordic Medico-Statistical Committee (NOMESCO) Classification of Surgical Procedures (NCSP) from 1996 onwards including: adenoidectomy, 2618, EMB30; tonsillectomy, 2614, EMB10; adenotonsillectomy, EMB20. There was no surgical procedure code for adenotonsillectomy prior to 1996, so this type of surgery was recorded when both surgical codes (2618, 2614) were entered on the same date.

### Selecting disease groups

We selected diseases based on whether they were likely to be affected by changes in the immune system (i.e. infections, allergies), and we added other disorders because they have been implicated in previous studies examining shorter-term health impacts for some of the focal surgeries of our study (i.e. respiratory infections). We also included broader disease groups (all circulatory, nervous system, endocrine, and autoimmune diseases) because immune dysfunction or dysbiosis could affect a wide range of organismal processes (see Table s2 for a full list of disease groups and their ICD codes). In Denmark, the International Classification of Diseases, 8th revision (ICD-8) was used from 1969 to 1993, and the 10th revision (ICD-10) from 1994 onwards. To reduce the possibility of encountering false negative results, we carried out power analyses in R and excluded some diseases that did not meet sample size thresholds needed to adequately test the null hypothesis of no association between predictor and incident disease (see Supplementary Methods - power analyses, for further details).

### Covariates adjusted for

To account for specific health and socioeconomic effects on the prevalence of focal diseases, we included the following covariates: Binary variables for maternal pre-existing conditions (see Table s2 for ICD codes) including hypertension (primary or secondary hypertension, hypertensive heart or renal disease), diabetes (i.e. type-I or type-II, malnutrition-related, other or unspecified), previous spontaneous or induced abortions; maternal pregnancy-related variables including gestation length (in weeks) and binary variables for the presence of maternal bleeding (i.e. haemorrhage, placenta praevia), fetal oxygen deprivation (i.e. hypoxia, asphyxia), and pregnancy oedema. Parental variables included a binary marker for whether either parent had ever been given a disease diagnosis within the same disorder group as the child to account for familial transmission, total number of years of education summed across both parents, and average income (in Dkr) across the study period summed across both parents. Birth-related variables included birth weight (in grams), birth season (calendar month, 1 to 12), birth cohort (3-year cohorts between 1978 - 2009 to account for any changes in diagnostic criteria over time), and the APGAR5 (Appearance, Pulse, Grimace, Activity, Respiration) score of 1-10 (maximally 2 points for each category) given to babies shortly after birth ranging from poor to excellent health. Other child-related variables included sex (0 = male, 1 = female), nationality (0 = Danish national, 1 = immigrant), parity (i.e. whether child was first, second, third, fourth (or higher)-born), and the region within Denmark (Hovedstaden = Copenhagen Area, Sjælland, Syddanmark, Midtjylland and Nordjylland) in which a child had resided for the longest period to account for possible regional differences in diagnoses.

### Statistical design and analysis

We used Cox regressions to model the relative risk for each of the 28 diseases up to age 30 (with age as timescale), depending on whether surgery occurred within the first 9 years of life. We validated the assumptions of the Cox regression model and ensured proportional hazards by stratifying for sex, birth cohort, birth season and demographic parity before estimating proportional hazards. A further 18 additional effects (described above and in Fig. 3) on disease prevalence were accounted for as regression covariates. To reduce the possibility of false positives, P values from Cox regressions (presented in Fig. 2–3) were Bonferroni corrected for 78 tests (α=0.05/78= 0.000641). To provide clinically useful statistics, absolute risks and number of patients needed to treat (NNT) before causing benefit or harm to one of them were also calculated from relative risks and disease prevalence within the first 30 years of life (see Supplementary Methods - converting RR to AR and NNT, for further details).

**Figure 2.**
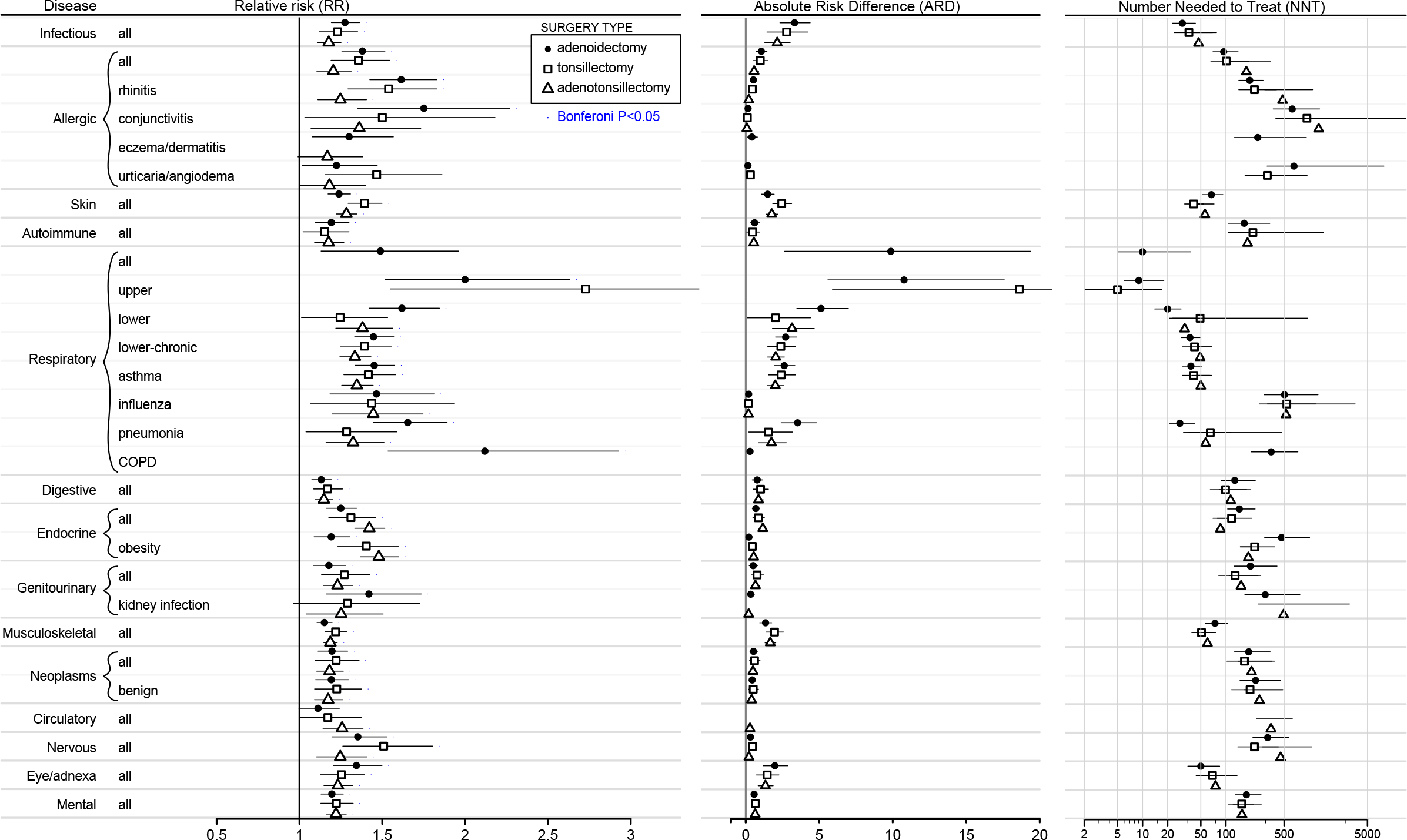
Risk of disease up to 30 years of age after removal of tonsils and adenoids in the first 9 years of life. Relative risks (RR] ± 95% CI are the exponents from Cox regressions that capture risk of each disease up to 30 years of age depending on whether surgery was performed. See key for surgery type. P-values significant after Bonferroni corrections for 78 tests are shown by a blue point above the upper confidence interval for each disease. RR values are presented only for analyses with sufficient power for hypothesis testing (see methods]. Absolute risk differences (ARD] ± 95% CI were estimated as ARD = 100 x CR x (1-RR), where CR (control risk] is the risk of the disease in the control sample and RR is the relative risk of disease in individuals post-surgery relative to the disease risk in the control sample that did not undergo surgery. Numbers needed to treat (NNT) ± 95% CI were estimated as NNT = 100 / ARD. Values above or below zero indicate ‘harm’ or ‘benefit’ of surgery, respectively, with values closer to zero indicating harm occurring more often to patients. For example, for risk of asthma after adenoidectomy (i.e. RR=1.45, 95% CI 1.33-1.57], the event rate in the control group (or control risk, CR] for asthma up to 30 years of age in our dataset was 5.8%, so ARD = 100 x 0.058 x (1-1.45] = 2.6 and NNT = 100/2.6=38. Relative risk of asthma was 1.45 and thus 45% higher after adenoidectomy compared to controls (no surgery], which translates to an absolute risk difference of 2.6% or 2.6 more cases of asthma per 100 treated patients. In other words, adenoidectomy needs to be performed in ̴38 children (100/2.6] to cause asthma to appear in one of them within the first 30 years of life.

**Figure 3.**
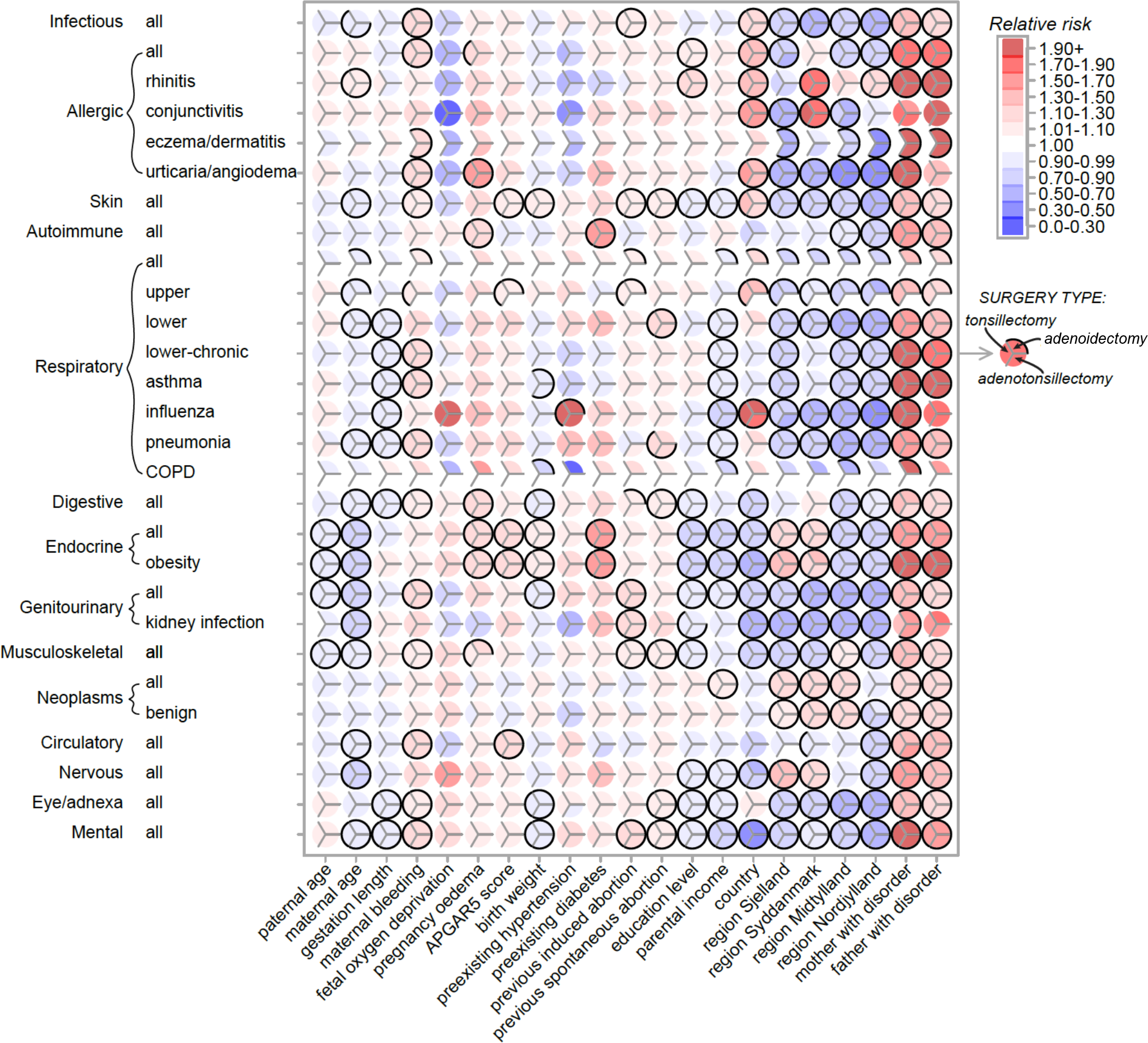
Disease risk patterns for covariates. Relative risk magnitude and direction correspond to red (increased relative risk] and blue (decreased relative risk] colors (see key, top right] derived from Cox regressions capturing the risk of diseases (vertical axis] within the first 30 years of life depending on 21 covariates (horizontal axis]. Within each circle there are three divisions corresponding to surgery type (see mid-right key], A black border indicates whether risk for that particular disease-covariate combination was significant after Bonferroni correction for 78 tests; a complete black border surrounding a circle indicates that risks were significant for all three surgeries. Disease risks for the covariate ‘region most lived in Denmark’ are relative to Hovedstaden (Copenhagen region].

### Estimating risks for non-immune diseases and conditions that surgeries aim to treat

To quantify whether disease risks due to the focal surgeries are likely to be justified given their proposed health benefits, we calculated relative risks, absolute risks and number needed to treat for conditions that these surgeries treat using the same samples and statistical setup as described above. Conditions that adenoidectomy, tonsillectomy and adenotonsillectomy treat include obstructive sleep apnea, sleep disorders, abnormal breathing, (chronic) sinusitis, labyrinthis, otitis media, and (chronic) tonsillitis (see Table s2 for ICD codes). To test whether these surgeries may affect risk for diseases known to be unrelated to the immune system, we estimated risk for osteoarthritis, cardiac arrhythmias, heart failure, acid-peptic disease and alcoholic hepatitis (see Table s2 for ICD codes) using the same sample and statistical setup described above. Results (see Table s3) showed that surgery did not modify risk up to 30 years of age for these non-immune diseases.

### Testing for biases in general health before undergoing surgery

With access to complete medical records from birth, we were able to test whether or not any such biases could influence the assessment of surgery effects on incident risk of disease. The null hypotheses tested were that there was no difference in general health between cases and controls for: (1) age at first disease diagnosis, or (2) age at any disease diagnosis for diseases recorded before surgery. Neither null hypothesis could be rejected, indicating that cases were no more sickly than controls prior to surgery for diseases that occurred in the first 9 years of life (see Supplementary Methods - testing for biases in early general health between cases and controls, for further details and findings).

## Results

We use relative risk, absolute risk, and number needed to treat to gain a balanced view of the actual general health effects that adenoidectomy, tonsillectomy or adenotonsillectomy were likely to have in the first 30 years of life in Denmark. We found that disease risks typically increased after any of the three surgeries were performed. For some, relative risks translated into substantial changes in absolute risk with the number needed to treat showing that only a few surgeries had to be performed to cause significant individual harm.

### The main impacts are on risk of respiratory disease

Tonsillectomy nearly tripled the relative risk of diseases of the upper respiratory tract (RR=2.72, 95% CI=1.54-4.80, Fig. 2, Table s4, S5(10)) and resulted in a substantial increase in absolute risk (ARD=18.61%, Table s4) and a small number needed to treat (NNT-harm=5, Table s4). Thus only about five tonsillectomies would need to be performed to cause diseases of the upper respiratory tract to appear in one of them. This suggests that the negative health consequences for this procedure in the overall population can be considerable.

Adenoidectomy more than doubled the relative risk of chronic obstructive pulmonary disorder (COPD, RR=2.11, 95% CI=1.53-2.92, Fig. 2, Table s4, S6(16)) and diseases of the upper respiratory tract (RR=1.99, 95% CI=1.51-2.63) and nearly doubled the relative risk of conjunctivitis (RR=1.75, 95% CI=1.35-2.26). This resulted in a substantial increase in absolute risk for upper respiratory tract disorders (ARD=10.7%, Table s4), but gave only very small increases for COPD (ARD=0.29%) and conjunctivitis (ARD=0.16%), consistent with the corresponding NNT values (NNT-harm: diseases of upper respiratory tract=9; COPD=349; conjunctivitis=624, Table s4). Although relative risk increases were reasonably similar for these diseases, the large differences in absolute risk were due to the overall prevalence of these disorders in the population. For example, diseases of the upper respiratory tract occur 40-50 times more frequently (i.e. in 10.7% of control individuals up to 30 years of age) than do COPD (0.25%) and conjunctivitis (0.21%).

### Other significant effects on long-term disease risks

For some diseases, even modest increases in relative risk (RR 1.17-1.65) resulted in relatively large increases in absolute risk (i.e. 2 to 9%) and relatively low number needed to treat values (i.e. NNT-harm < 50), which was largely due to the high prevalence of these diseases in the population (risk in controls ranged from 5 to 20%, Table s4). They mainly included respiratory diseases (groups including: all, lower, lower-chronic, asthma, pneumonia), infectious diseases (all), skin diseases (all), musculoskeletal (all) and eye/adnexa (all). For example, adenotonsillectomy significantly increased relative risk of infectious diseases by 17% (RR=1.17, 95% CI=1.10-1.25, Table s4, S7(1)), but because infectious diseases are relatively common (i.e. 12%, Table s4), the absolute risk increase was 2.14%, so that about 47 adenotonsillectomy surgeries would need to be performed for an extra infectious disease to occur as a result of one of these surgeries (Table s4).

When all 28 disease groups were considered, there were small but significant and consistent increases in relative risk for most of them: 94% of disease risks were significantly increased before Bonferroni correction, 78% after. Thus, the negative health consequences of these surgeries within the first 30 years of life are likely to be widespread, affecting a range of tissues and organ systems, underlining the importance of the uninterrupted presence of adenoids and tonsils for normal development. The removal of these organs early in life appears to slightly but significantly perturb many processes important for later-life health.

### Risk of conditions that surgeries directly aimed to treat were mixed

Risks for conditions that the focal surgeries of our study aimed to treat were mixed (Table s8). Surgery significantly reduced long-term relative risk for some conditions (i.e. 7 out of 23 or 30% of our disease-specific analyses), whereas there were no significant risk modifications for others (10 or 43%), and some significant increases in relative risk observed for a few conditions (6 or 26%).

For example, adenoidectomy resulted in significantly reduced relative risk for sleep disorders (RR=0.30, ARD=-0.083%), and all surgeries significantly reduced risk for tonsillitis and chronic tonsillitis (i.e. RR ranging between 0.09-0.54; ARD ranged between -0.29% to -2.10%). For abnormal breathing, there was no significant change in relative risk up to 30 years of age after adenoidectomy, tonsillectomy or adenotonsillectomy (Table s8), no change in relative risk for sinusitis after adenoidectomy or tonsillectomy, and no change in relative risk of labyrinthitis after adenotonsillectomy (Table s8). Conditions where relative risk significantly increased during the follow-up period included otitis media, which showed a 2 to 4-fold increase after surgery (i.e. RR ranged from 2.06-4.84; ARD ranged between +5.3% to 19.4%), and sinusitis, which increased significantly after adenotonsillectomy (RR-1.68; ARD +0.11%, Table s8).

This suggests that short-term health benefits these surgeries may provide for these conditions often do not continue longer-term, at least up to 30 years of age examined in this study. Indeed, apart from the consistently reduced risk for tonsillitis (after any of these surgeries) and sleep disorders (after adenoidectomy), longer-term risks for most other conditions (abnormal breathing, sinusitis, chronic sinusitis, labyrinthitis, otitis media) were either significantly higher post-surgery or not significantly different.

### Risk patterns for covariates

The many interesting patterns of disease risk based on covariates included in the analysis (summarized in Fig. 3, full details Table s5–s7) highlight the complexity of the factors influencing the focal disease risks of our study. For example, consider covariates that significantly modified risks of diseases of the upper respiratory tract (Fig. 3), and their largest increases in relative (RR=1.99-2.72) and absolute risk (ARD=10.77%-18.61%) after adenoidectomy and tonsillectomy (Fig. 2). The risks for these diseases slightly decreased for offspring born to older mothers (RR=0.96, 95% CI=0.95-0.98 for both surgeries), slightly increased (tonsillectomy analysis) when maternal bleeding occurred during pregnancy (RR=1.07, 95% CI=1.03-1.12), increased (both analyses) with APGAR5 score (RR=1.09, 95% CI=1.04-1.13, both surgeries), increased (both analyses) when mothers had a previous induced abortion (RR=1.09, 95% CI=1.06-1.12, both surgeries), increased in immigrants relative to Danish nationals (RR=1.40, 95% CI=1.33-1.47, both surgeries), decreased in those living anywhere in Denmark other than Copenhagen (RR=0.69 to 0.93), and increased when fathers or mothers had a history of the same disease (RR=1.29 to 1.38). All such effects were accounted for in our risk estimates and their significance (Fig. 2 and Fig. 3).

Some covariates showed very consistent effects on risk for many diseases. For example, the most consistent effect in Fig. 3 based on relative magnitude of disease risk (e.g. RR ranged from 1.10 to 3.71) and the direction of effect (i.e. risk was increased for every disease) occurred when either parent had been diagnosed previously with the same disease. Other covariates such as parental education, income and country of origin also showed many significant effects, but risk direction varied considerably depending on the disease considered. For example, mental disorders were less frequent in Danish nationals than in immigrants (RR 0.48 to 0.49), yet risk of influenza was much higher in Danes (RR 1.89 to 2.06). Several diseases were sensitive to many covariates (e.g. endocrine, mental) suggesting complex causation. We discuss other interesting and potentially important effects of these covariates in the Supplementary Results.

## Discussion

While most otorhinolaryngologists have focused on the short-term consequences of their procedures for the symptoms that they treat [14, 22–24], not all have been aware of the full range of long-term risks. These are difficult to evaluate due to many potential confounding effects that are unrelated to the original diagnosis and may obscure estimation of disease risk. Using the Danish public health data allowed us to unambiguously control for many medical, socioeconomic and statistical confounders, and we are confident that our estimates of increased disease risks later in life associated with removal of tonsils and adenoids are robust and quantitatively reliable. Our main findings are that tonsillectomy nearly triples the risk of diseases of the upper respiratory tract and that adenoidectomy doubles the risk of chronic obstructive pulmonary disorder (COPD) and diseases of the upper respiratory tract and nearly doubles the risk of conjunctivitis. There were corresponding large increases in absolute risk for diseases of the upper respiratory tract. Smaller but significantly elevated risks were found for a broad range of other diseases, which translated into measureable increases in absolute risk for diseases with high prevalence in the population (i.e. infectious, skin, musculoskeletal and eye/adnexa diseases). These findings add to previous research on single diseases that have shown increased risks of breast cancer [6] and premature acute myocardial infarctions [7] as a consequence of these surgeries. In contrast, the long-term benefits of surgery were generally minor and sometimes actually increased risk for the conditions they aimed to treat.

Our results again raise the important general issue of when the benefits of operating are sufficient to outweigh the combined short-and long-term morbidity costs. For much of the last century these operations were relatively common, but their frequencies have declined in more recent medical practice [25, 26], with the availability of alternative treatments for infections in ear, oral and nasal cavities coinciding with heightened appreciation of the possible shortterm risks of surgery [27]. The long-term costs that our study documents and quantifies add a novel perspective to these considerations. They suggest that renewed discussion may be timely, especially given that these surgical procedures remain amongst the most common medical interventions in childhood [3, 4]. It is important to appreciate that the cumulative long-term impact of surgery depends on the prevalence of specific conditions in the population, as we document for many disease categories, for these trends are not straightforward to extrapolate from relative risk ratios. Thus, the impacts of tonsillectomy and adenoidectomy on the absolute risk of diseases of the upper respiratory tract were substantial because these conditions are prevalent, whereas the impacts of adenoidectomy on the absolute risks of COPD and conjunctivitis were small because those diseases have low prevalence.

Our results suggest that there is a direct connection between surgical removal of immune organs in the upper respiratory tract during childhood and increased risk of respiratory and infectious diseases later in life. Given that tonsils and adenoids are part of the lymphatic system that itself is part of the circulatory system and plays a key role both in the normal development of the immune system and in pathogen screening during childhood and early life [3], it is perhaps not surprising that their removal would impair pathogen detection and increase risk of respiratory and infectious diseases emerging later in life. However, other correlations between these surgeries and diseases of the skin, eyes, and musculoskeletal system seem unlikely to involve direct causation, and will thus require further investigation. The growing body of research on developmental origins of disease [15, 28] has convincingly demonstrated that even small perturbations to fetal and childhood growth and development can have lifelong consequences for general health. Thus, causative pathways to disease are often complex and not always obvious given the interconnectedness of bodily systems with evolved tradeoffs and compensatory mechanisms. We note that our study sample could not address risks of diseases in those over 30 years of age, the limit of our sample, and even though records of the entire population of Denmark were available, we did not have large enough samples for the rarer diseases to obtain reliable estimates of risk.

## Conclusion

This is the first study to estimate long-term health consequences of tonsillectomies and adenoidectomies for a broad range of diseases. Impacts on long-term risks were significant for many diseases and large for some. The strength of our study is its very large coverage of a single relatively homogeneous Danish population, but this may mean that some of our results will not generalize to other populations. The effects of tonsillectomy and adenoidectomy that we uncovered in the Danish population warrant renewed evaluation of potential alternatives to surgery. Our study shows that both absolute risks and the number of patients needed to treat before health risks later in life become apparent were more consistent and widespread than the immediate population-wide benefits of childhood surgery for subsequent health within the first 30 years of life.

**Table 1.** **Characteristics of the study samples for initial surgery and control groups.** Numbers (n), means (μ), percentages (%, binary traits multiplied by 100, so they are percentages) and Standard Deviations (SD, in brackets) are provided.

|  | adenoidectomy | tonsillectomy | adenotonsillectomy | control | total |
| --- | --- | --- | --- | --- | --- |
| sample size (n) | 22637 | 39685 | 42384 | 1648394 | 1753100 |
| paternal age ($\mu$) | 30.1 (5.8) | 29.7 (5.6) | 30.1 (5.7) | 31.3 (5.7) | 31.2 (5.7) |
| maternal age ($\mu$) | 27.0 (4.9) | 26.8 (4.7) | 27.2 (4.8) | 28.5 (4.8) | 28.4 (4.9) |
| gestation length, weeks ($\mu$) | 39.4 (1.9) | 39.7 (1.6) | 39.6 (1.7) | 39.7 (1.6) | 40.0 (1.6) |
| maternal bleeding (%) | 9.6 (0.3) | 8.1 (0.2) | 9.0 (0.2) | 7.3 (0.2) | 7.4 (0.2) |
| fetal oxygen deprivation (%) | 0.13 (0.04) | 0.13 (0.04) | 0.09 (0.03) | 0.09 (0.03) | 0.09 (0.03) |
| pregnancy oedema (%) | 1.44 (0.12) | 1.42 (0.12) | 1.08 (0.10) | 0.79 (0.09) | 0.82 (0.09) |
| APGAR 5 score 1-10 ($\mu$) | 9.8 (0.7) | 9.8 (0.5) | 9.8 (0.6) | 9.8 (0.6) | 9.8 (0.6) |
| birth weight (grams) ($\mu$) | 3377 (597) | 3443 (537) | 3448 (574) | 3499 (541) | 3495 (543) |
| pre-existing hypertension (%) | 0.26 (0.05) | 0.15 (0.04) | 0.27 (0.05) | 0.33 (0.06) | 0.32 (0.06) |
| pre-existing diabetes (%) | 0.40 (0.06) | 0.35 (0.06) | 0.50 (0.07) | 0.39 (0.06) | 0.39 (0.06) |
| previous induced abortion (%) | 20.2 (0.4) | 19.5 (0.4) | 19.6 (0.4) | 18.2 (0.3) | 18.3 (0.3) |
| previous spontaneous abortion (%) | 16.3 (0.3) | 15.5 (0.3) | 15.6 (0.3) | 14.2 (0.3) | 14.3 (0.3) |
| education, combined total yrs ($\mu$) | 24.4 (4.9) | 24.5 (4.6) | 24.4 (4.6) | 25.6 (4.7) | 25.5 (4.8) |
| income, combined average Dkr. ($\mu$) | 381030 (188098) | 394585 (163759) | 361933 (145965) | 362241 (172797) | 363188 (172296) |
| nationality, 0=Danish, 1=other (%) | 5.58 (0.23) | 4.33 (0.2) | 7.69 (0.27) | 6.89 (0.25) | 6.83 (0.25) |
| region in Denmark most lived (%) |  |  |  |  |  |
| Hovedstaden | 29.1 (0.4) | 27.4 (0.4) | 23.6 (0.4) | 28.7 (0.4) | 28.5 (0.4) |
| Sjælland | 21.8 (0.4) | 18.6 (0.3) | 16.2 (0.3) | 14.5 (0.3) | 14.7 (0.3) |
| Syddanmark | 23.1 (0.4) | 19.7 (0.4) | 22.1 (0.4) | 22.4 (0.4) | 22.3 (0.4) |
| Midtjylland | 15.0 (0.3) | 23.3 (0.4) | 24.9 (0.4) | 23.7 (0.4) | 23.6 (0.4) |
| Nordjylland | 10.8 (0.3) | 10.7 (0.3) | 13.0 (0.3) | 10.5 (0.3) | 10.6 (0.3) |
| birth year ($\mu$) | 1988 (8) | 1987 (6) | 1991 (7) | 1994 (8) | 1993 (8) |
| birth season, months 1-12 ($\mu$) | 6.4 (3.3) | 6.3 (3.3) | 6.4 (3.3) | 6.4 (3.3) | 6.4 (3.3) |
| sex, 0=male, 1=female (%) | 40.7 (0.4) | 58.9 (0.4) | 44.5 (0.5) | 48.6 (0.5) | 48.7 (0.5) |
| demographic parity (%) |  |  |  |  |  |
| first born | 46.2 (0.5) | 45.5 (0.5) | 47.9 (0.5) | 44.1 (0.5) | 44.3 (0.5) |
| second born | 37.2 (0.4) | 38.2 (0.4) | 35.1 (0.4) | 37.3 (0.4) | 37.3 (0.4) |
| third born | 12.0 (0.3) | 12.3 (0.3) | 12.6 (0.3) | 13.8 (0.3) | 13.7 (0.3) |
| fourth (or higher born) | 4.33 (0.20) | 3.84 (0.19) | 4.33 (0.20) | 4.7 (0.21) | 4.67 (0.21) |

## Supplementary Contents

**Table S1.**
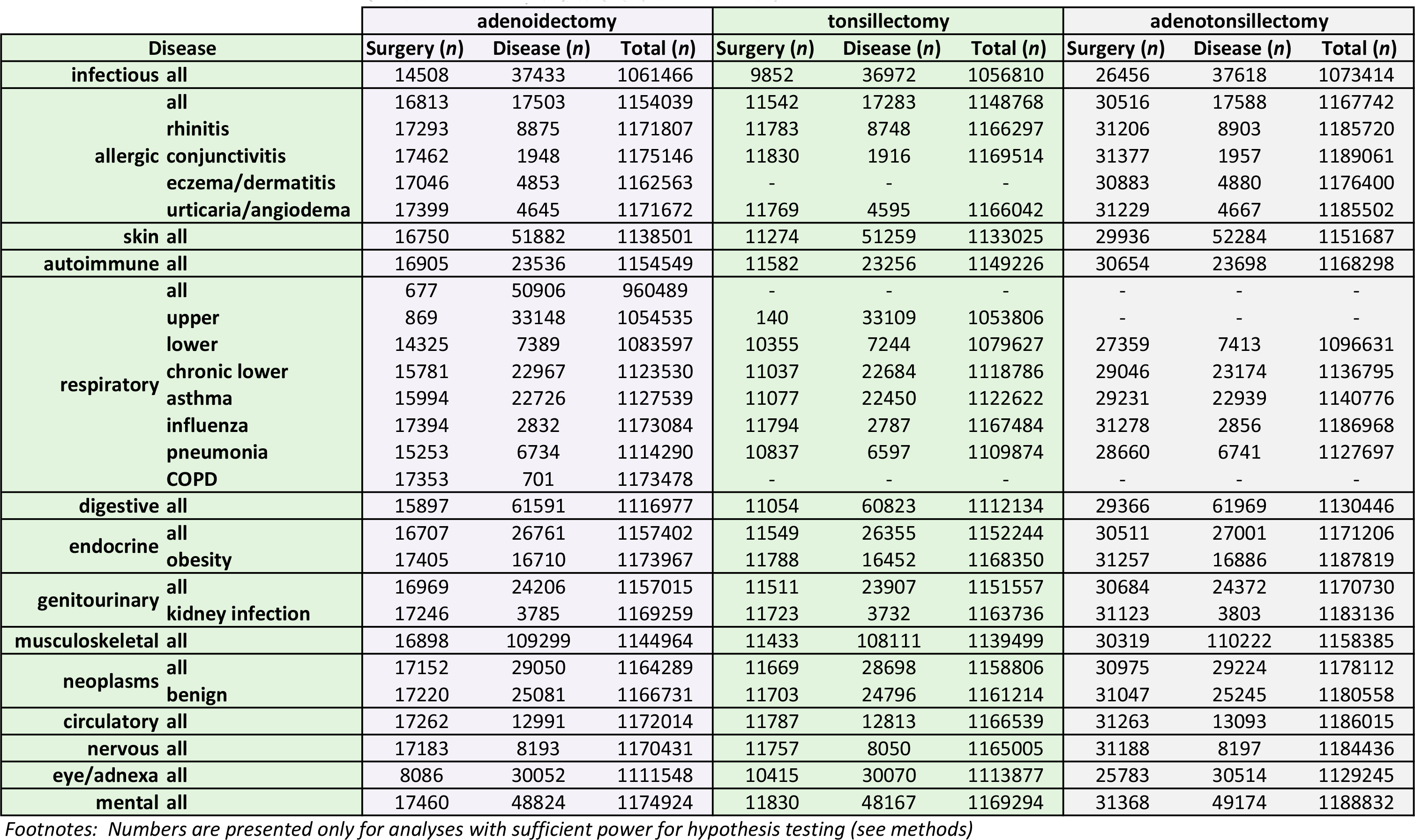
**Sample sizes used for each analysis including the number of individuals who underwent surgery between birth and 9 years, number diagnosed with a particular disease after 9 years and up to 30 years of age, and total sample size available**.

**Table S2.**
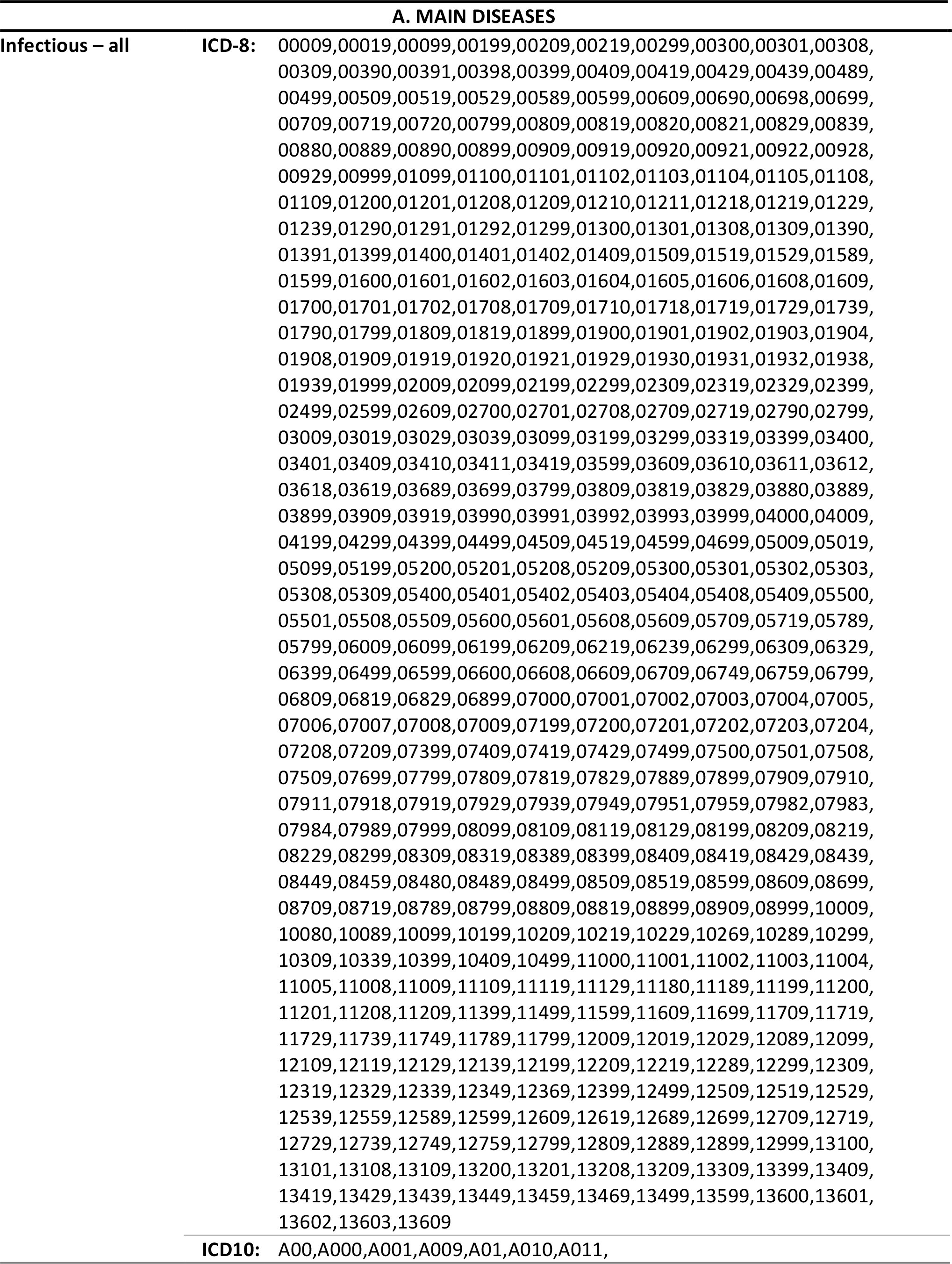
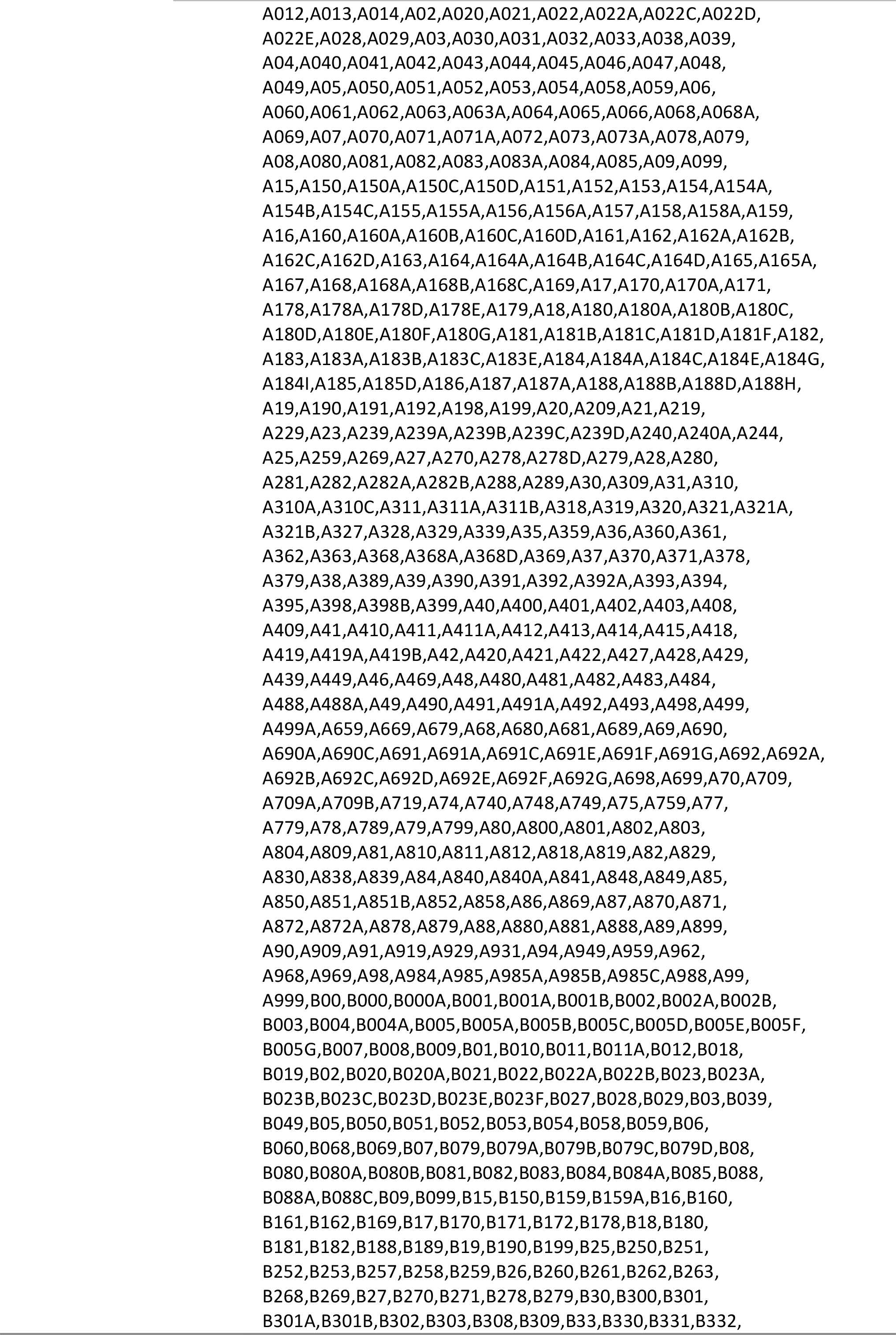
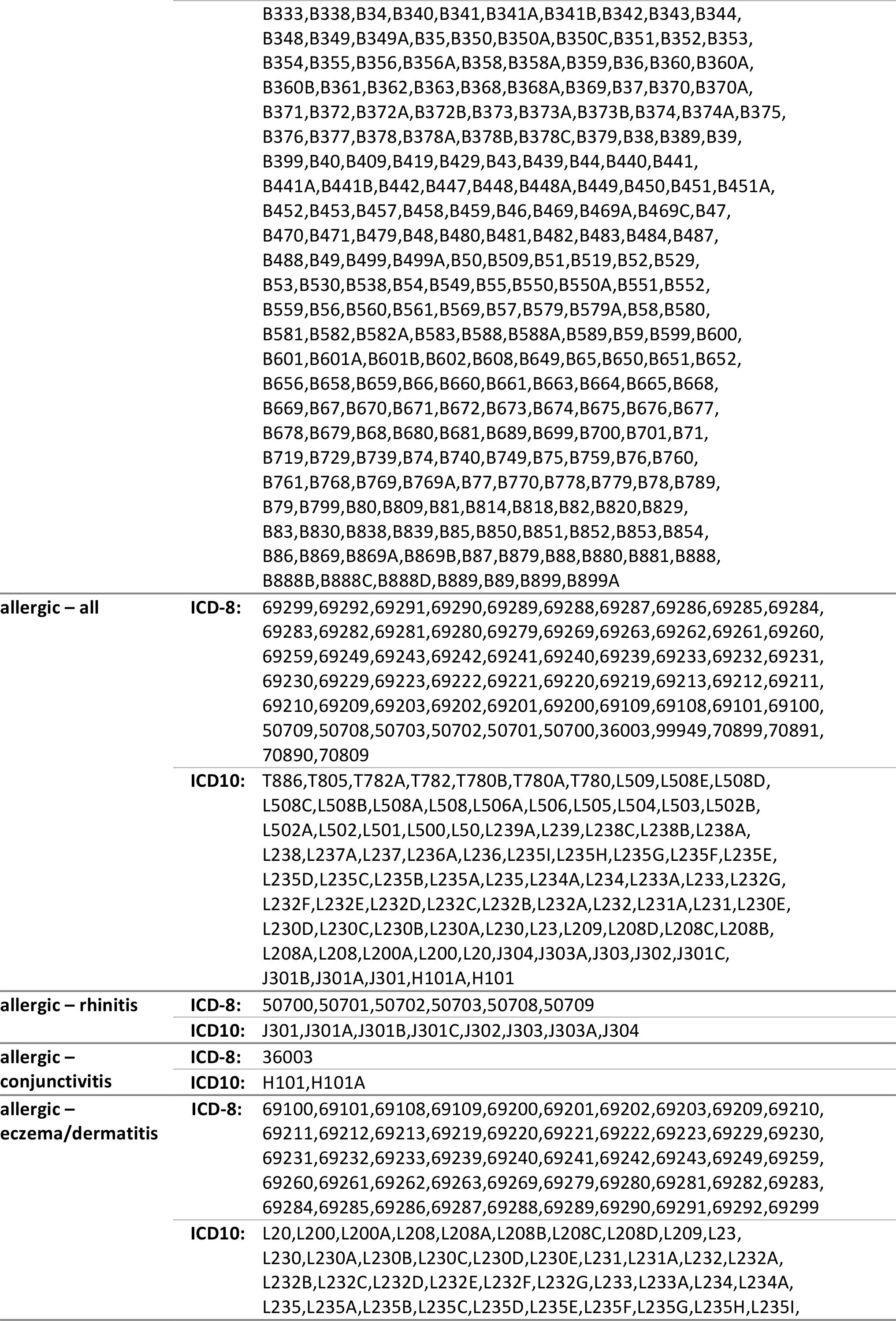
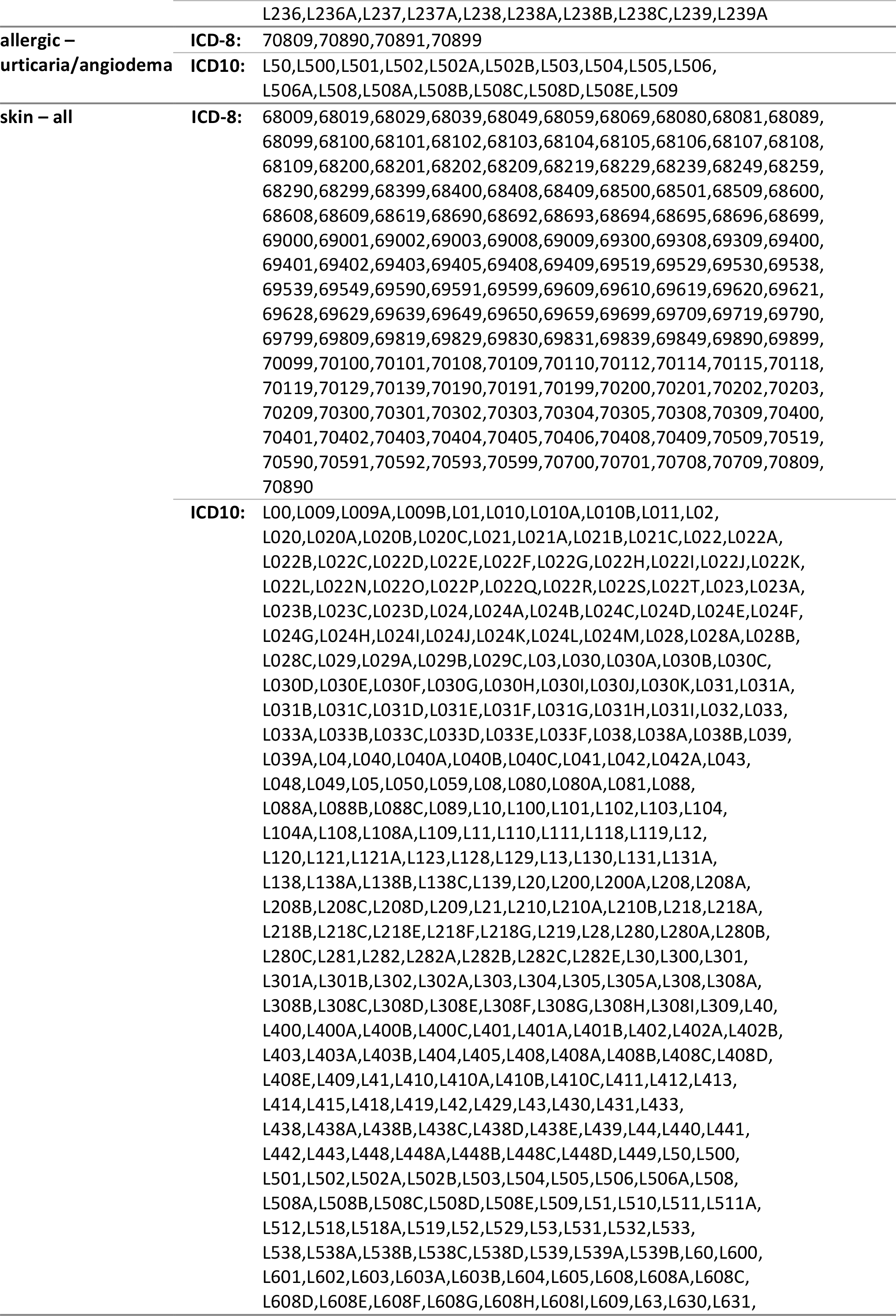
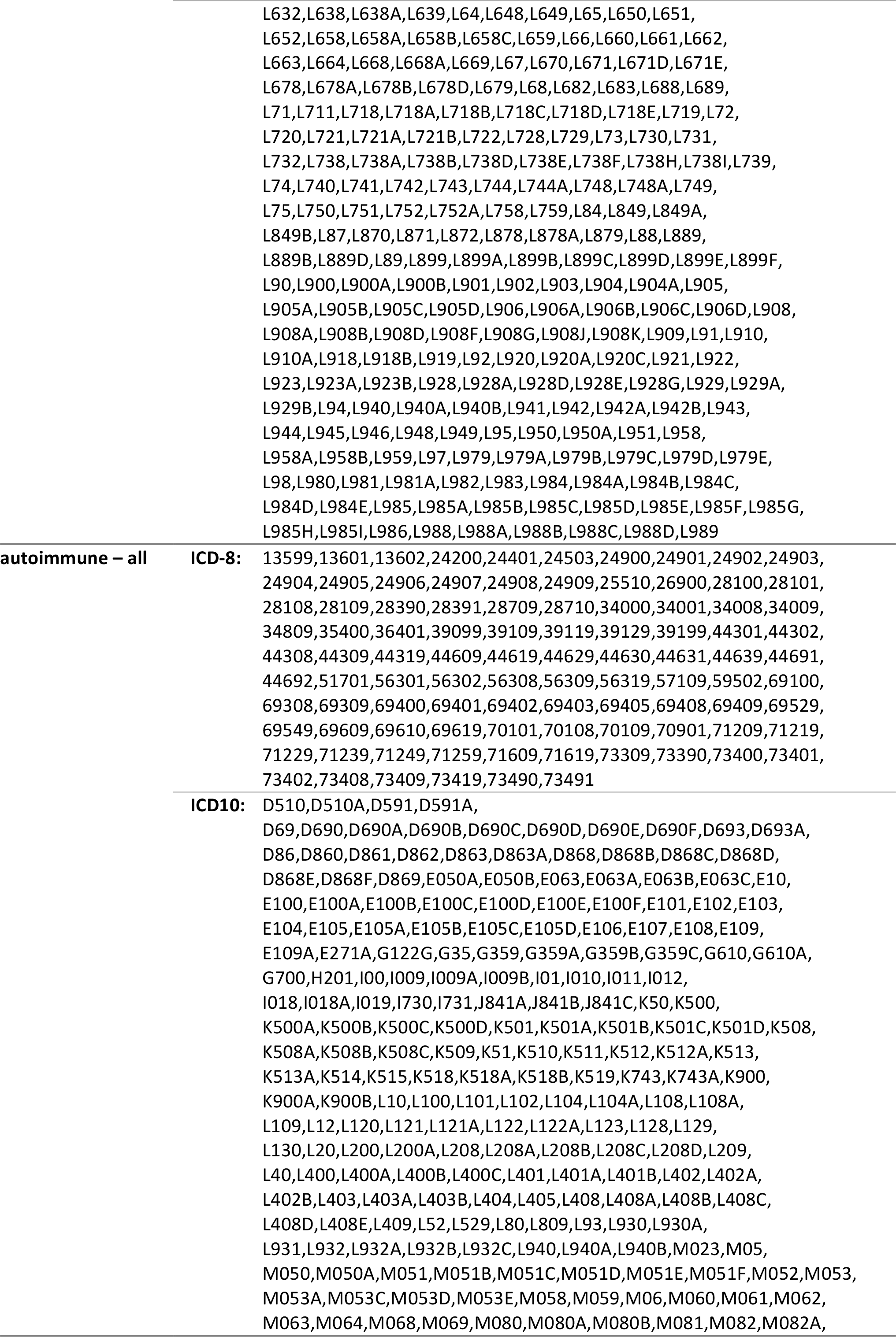
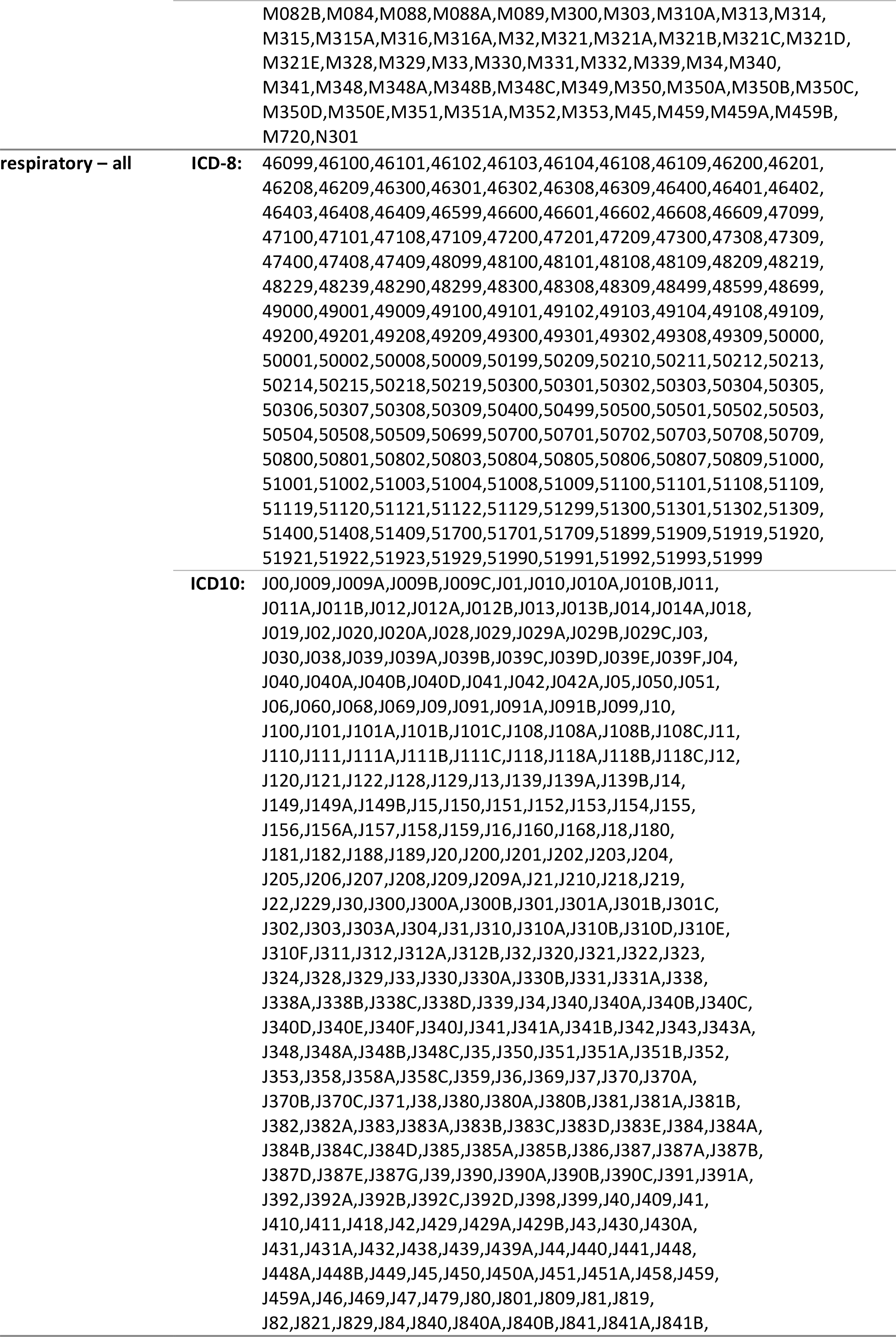
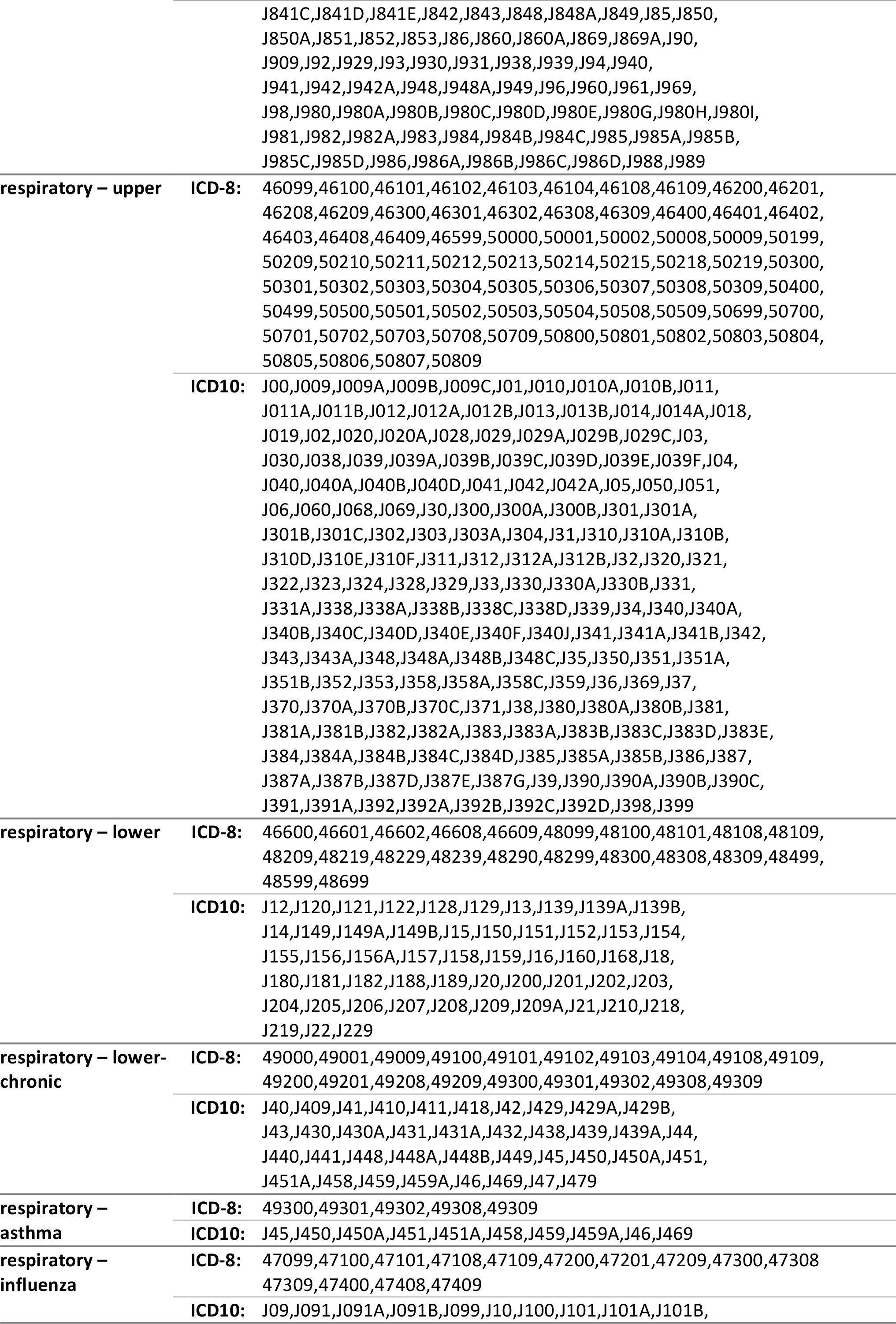
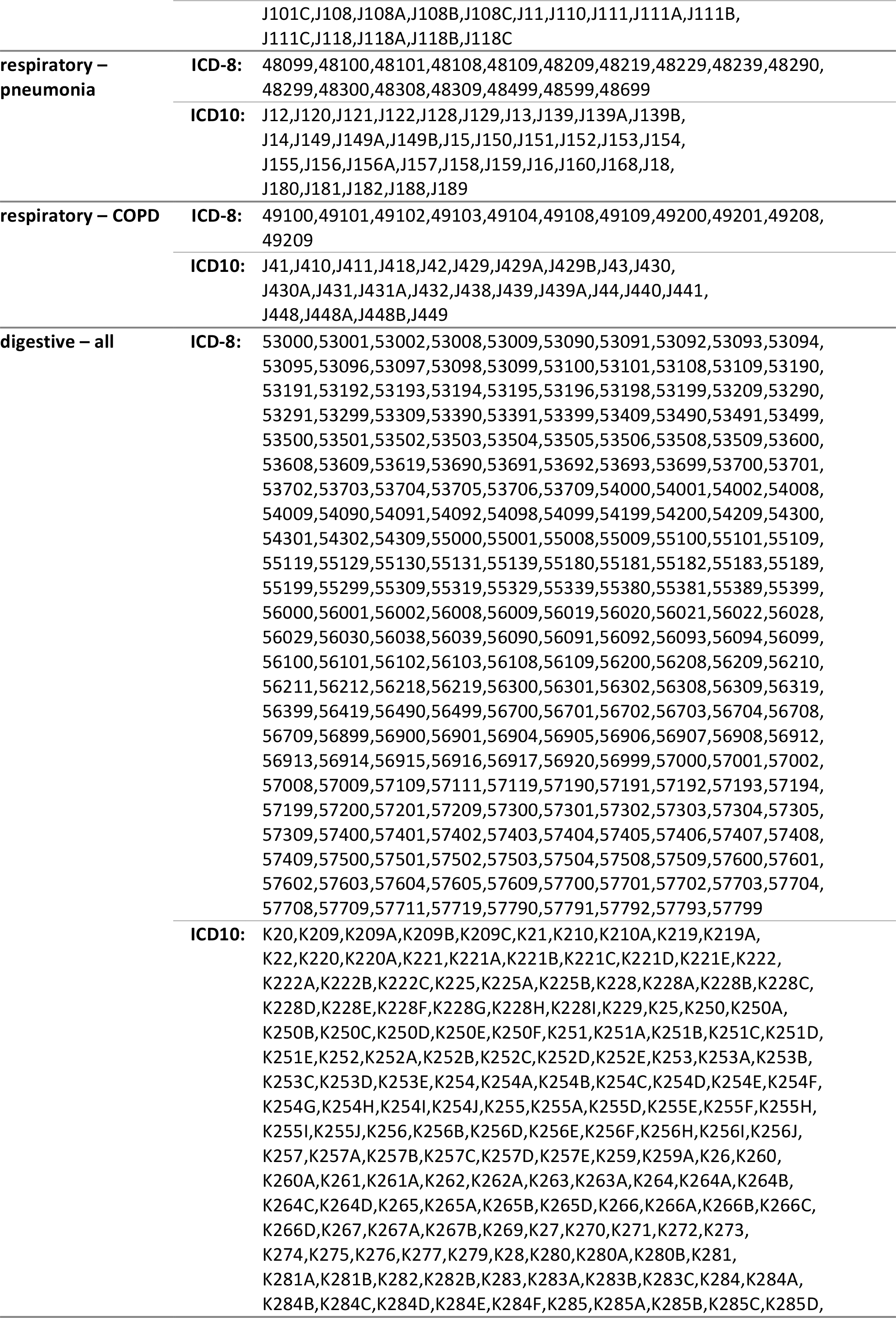
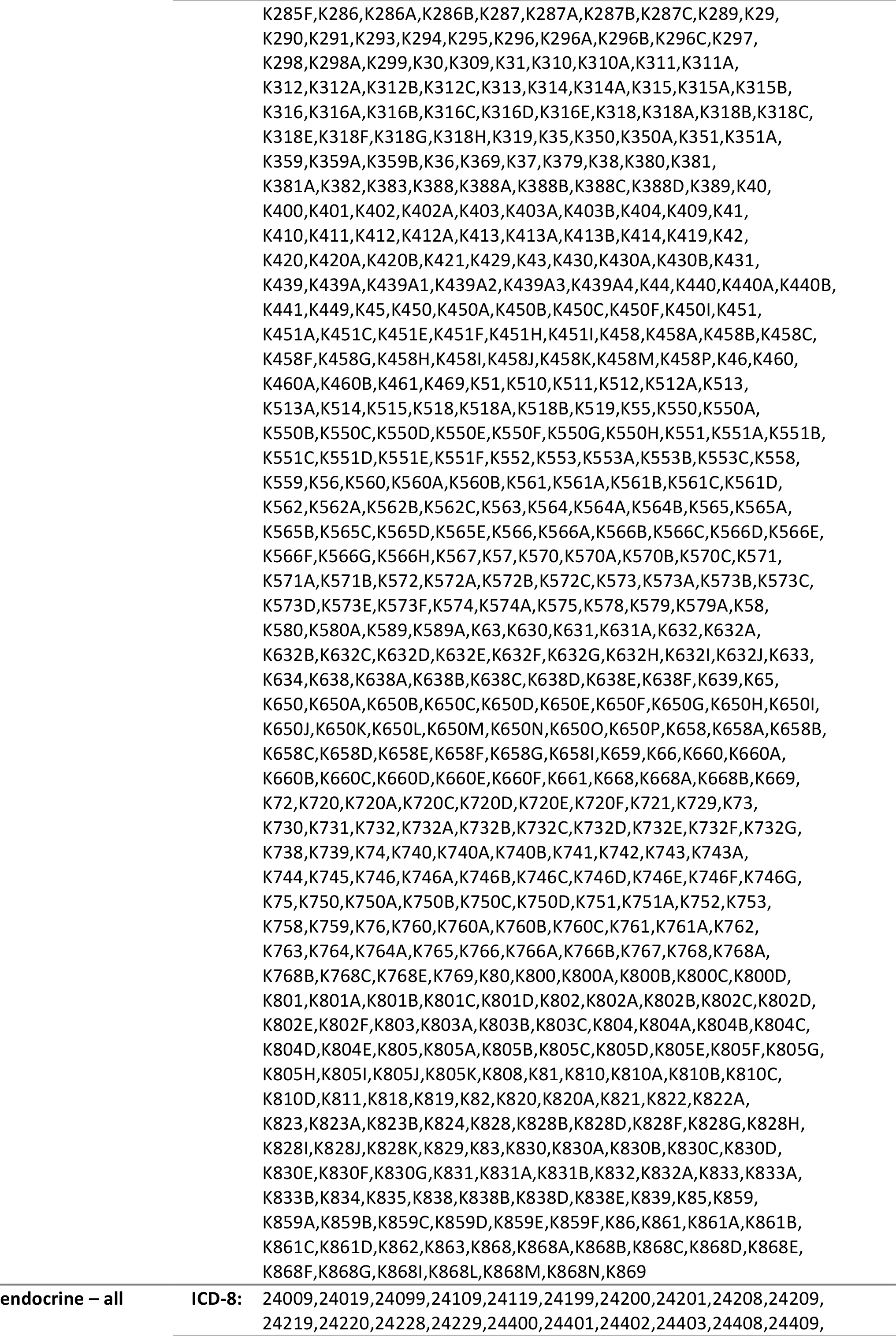
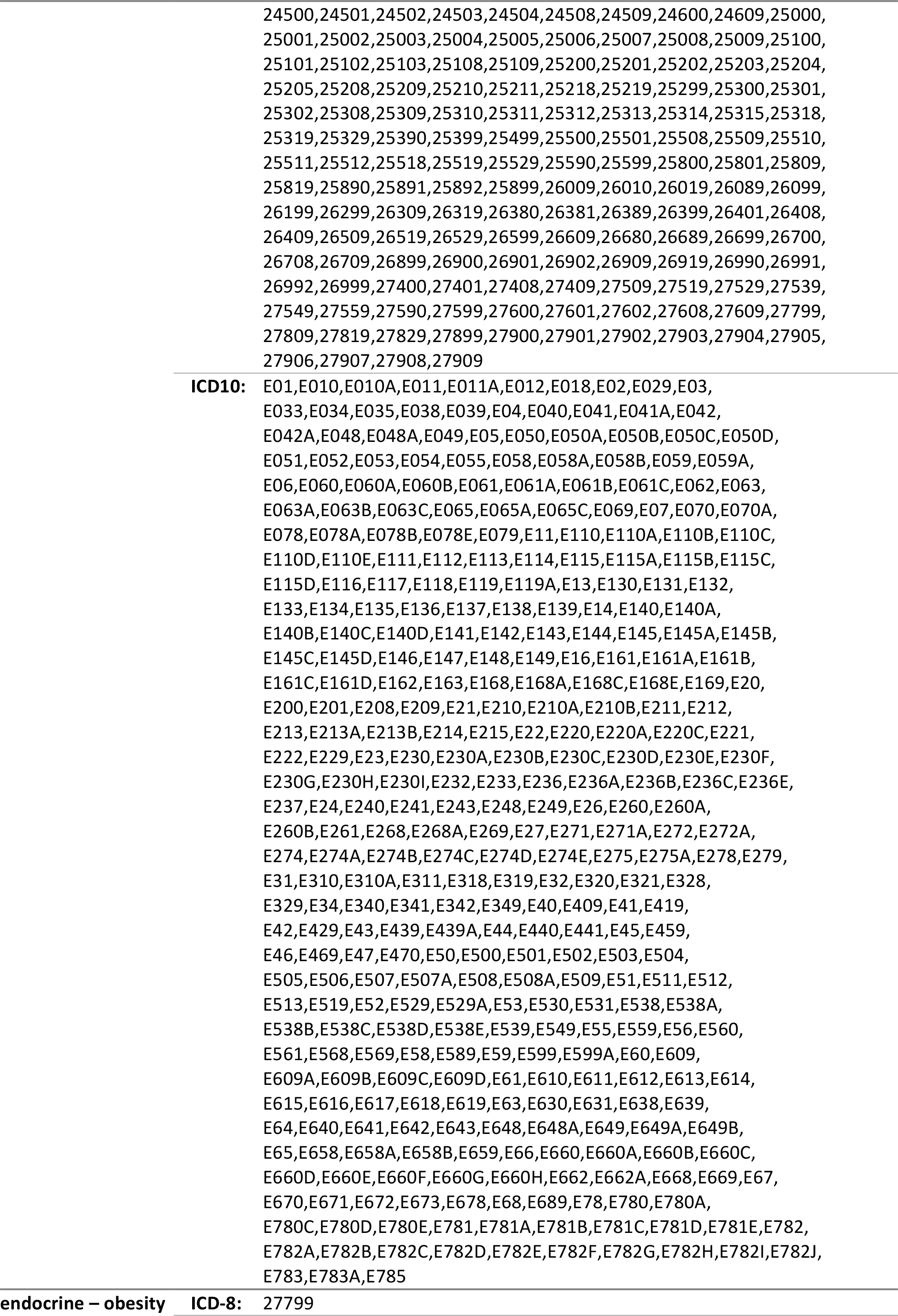
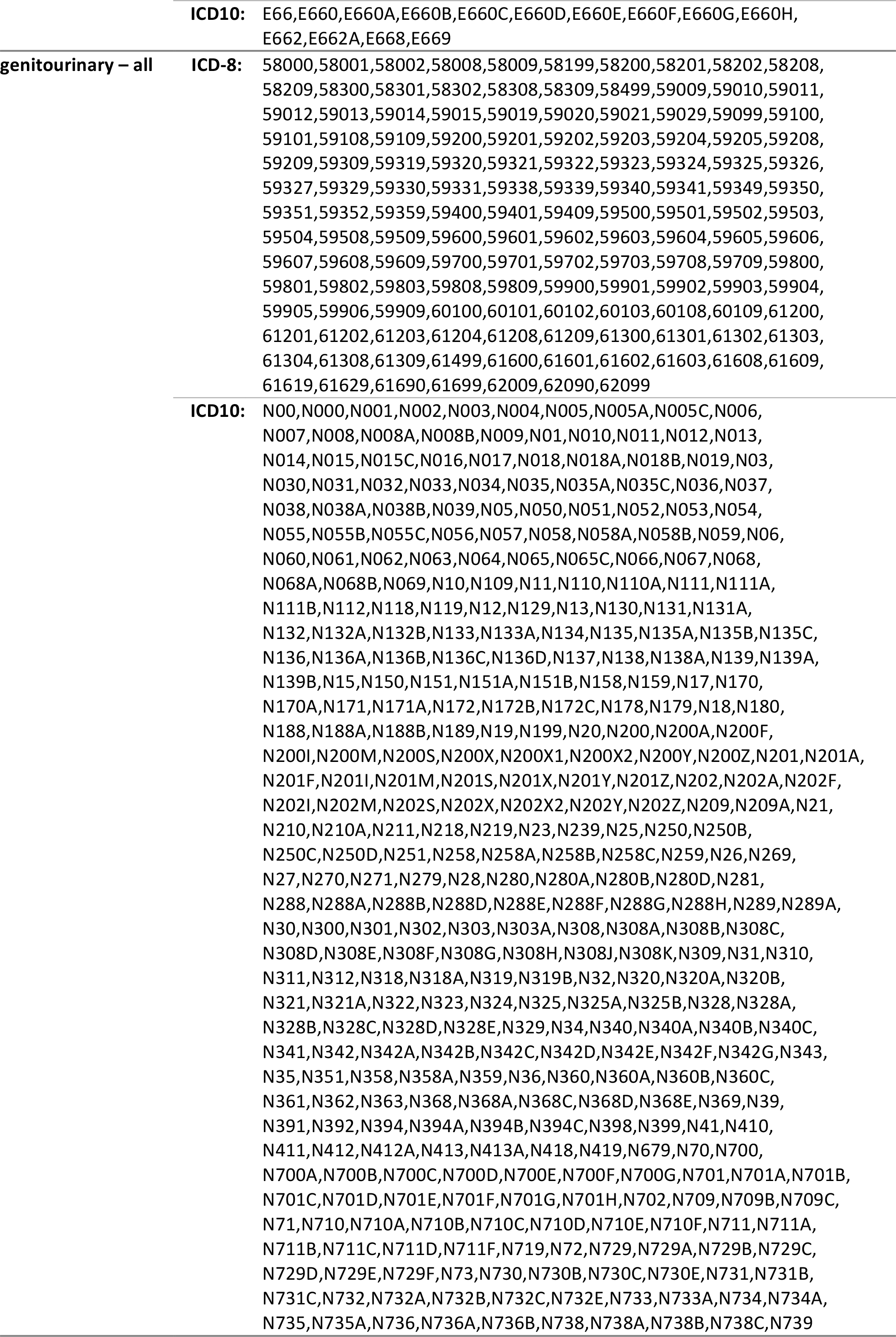
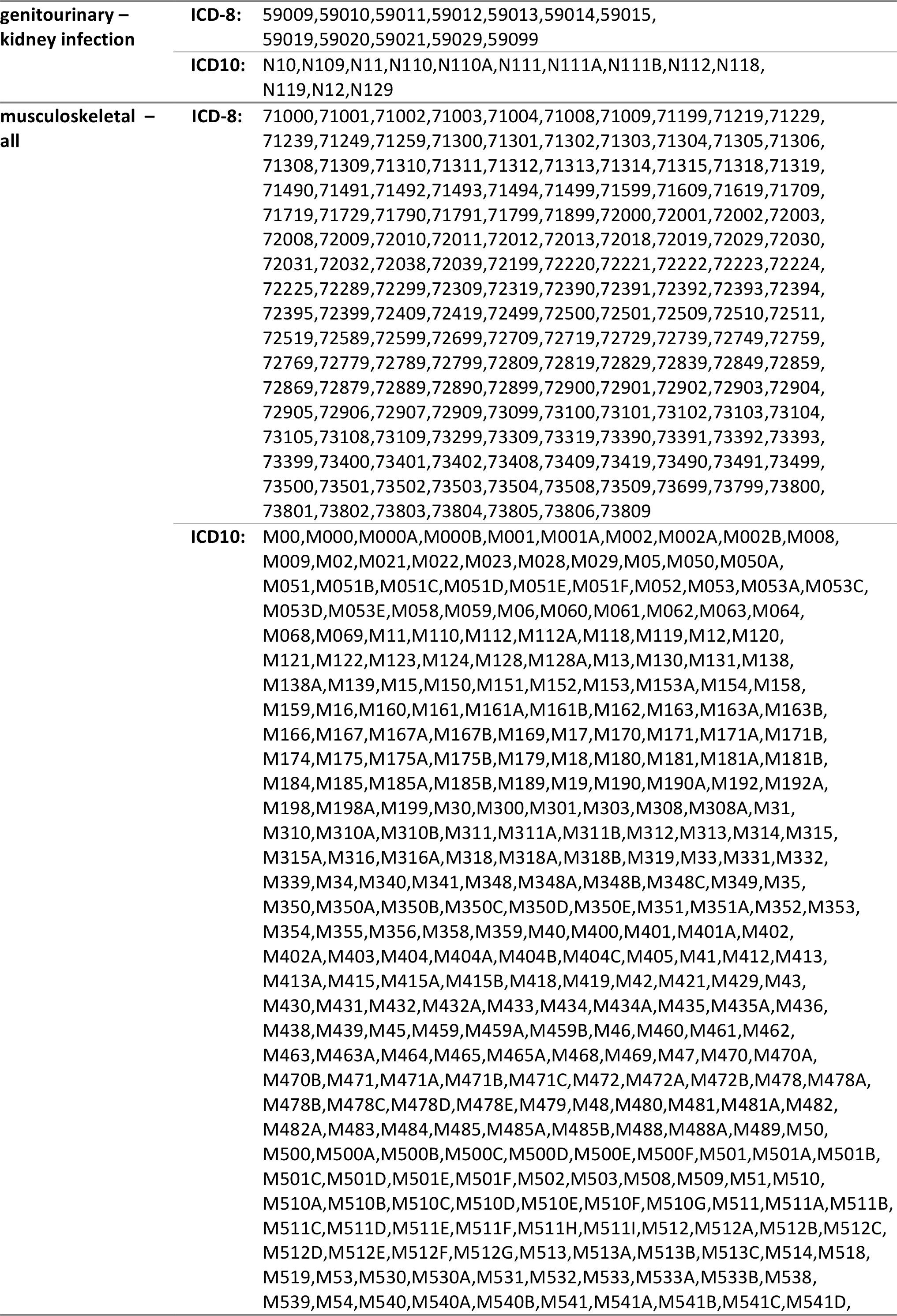
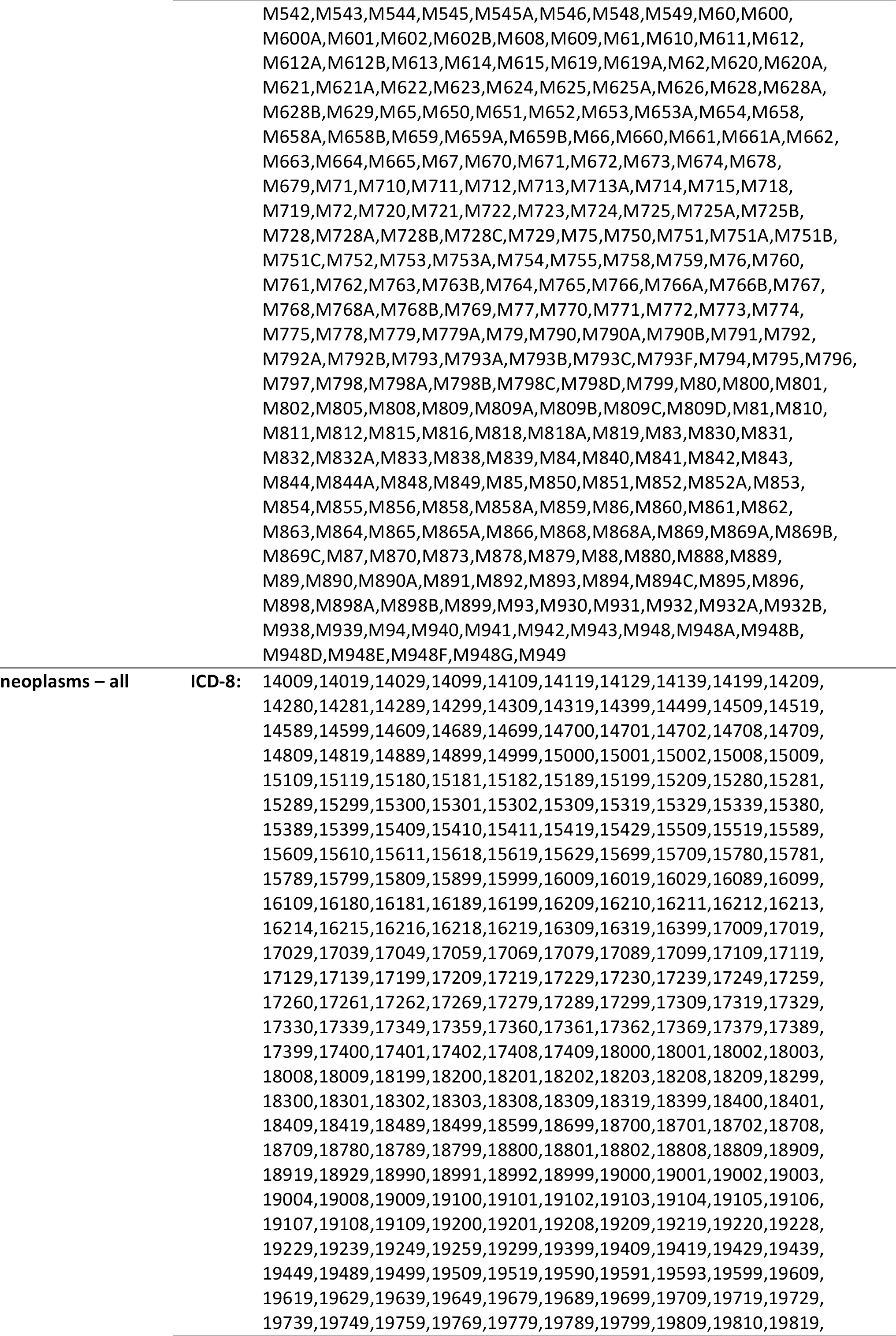
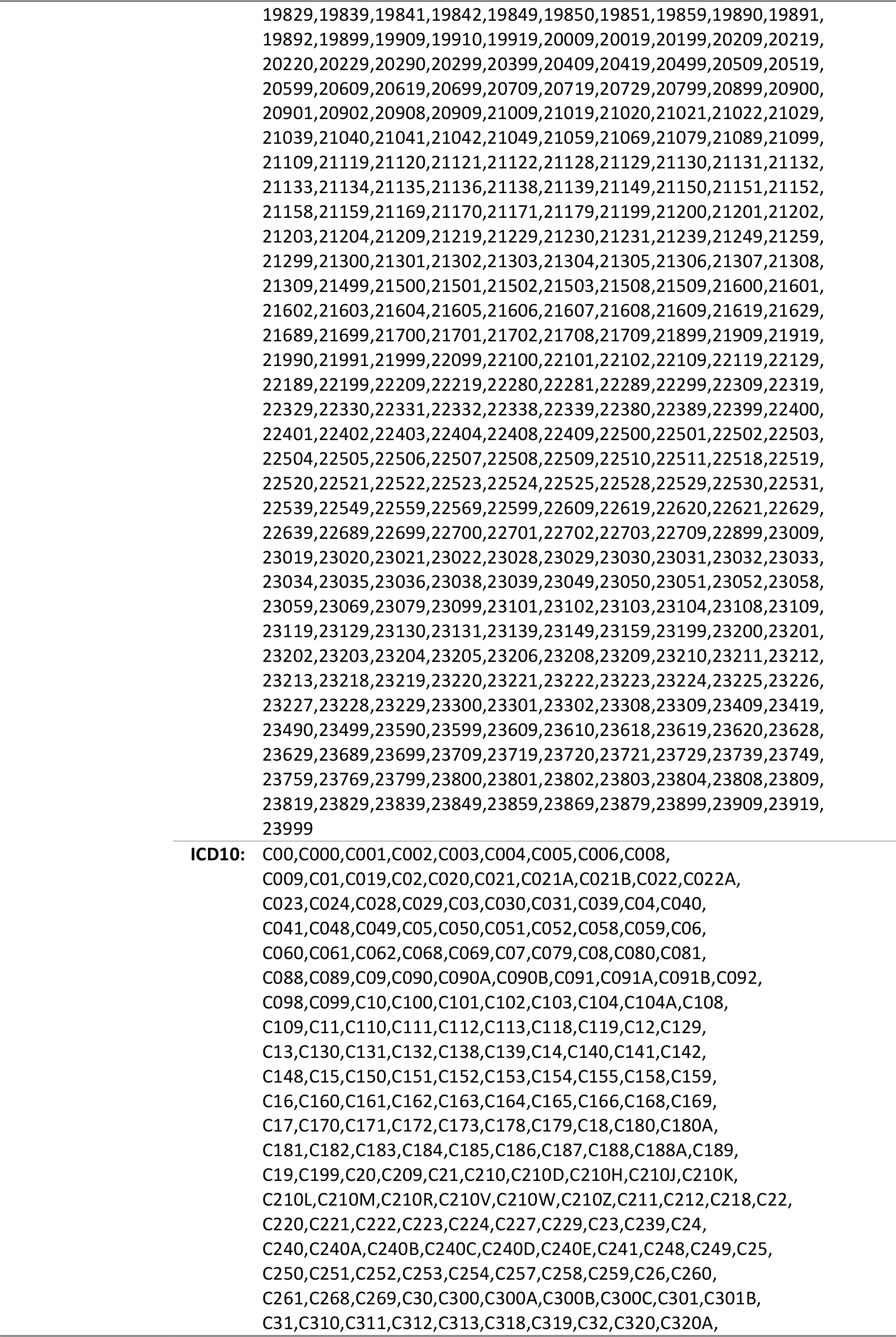
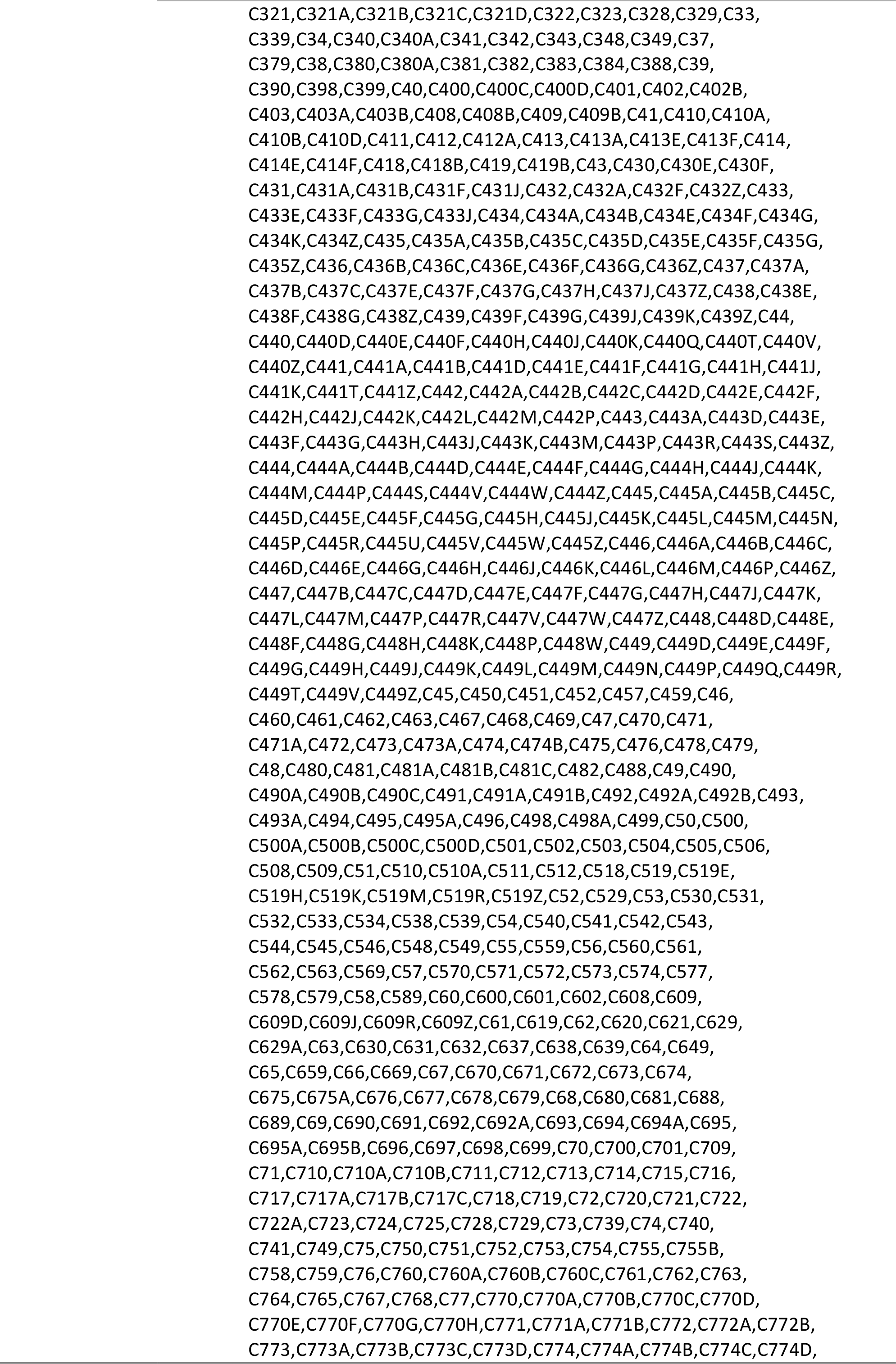
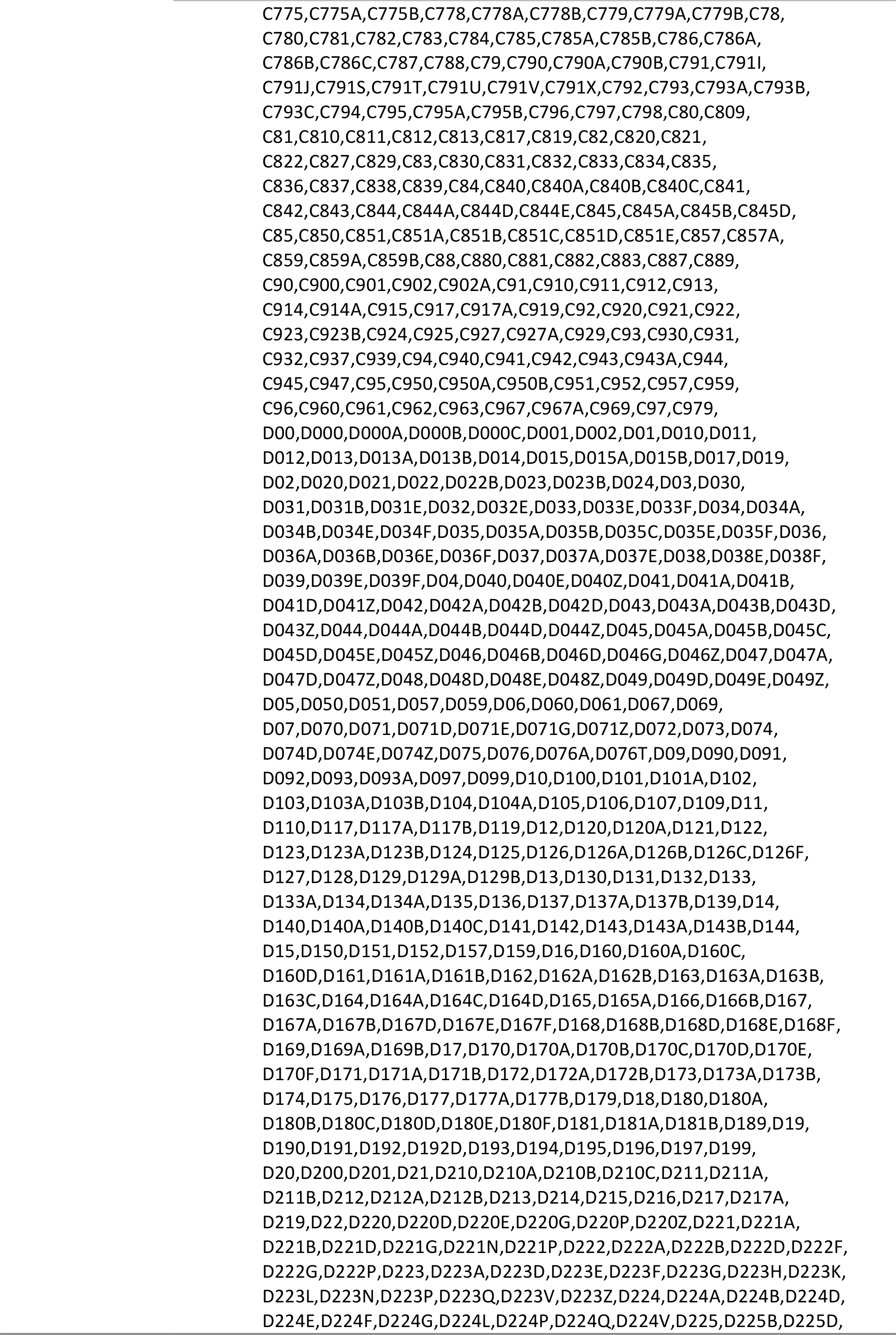
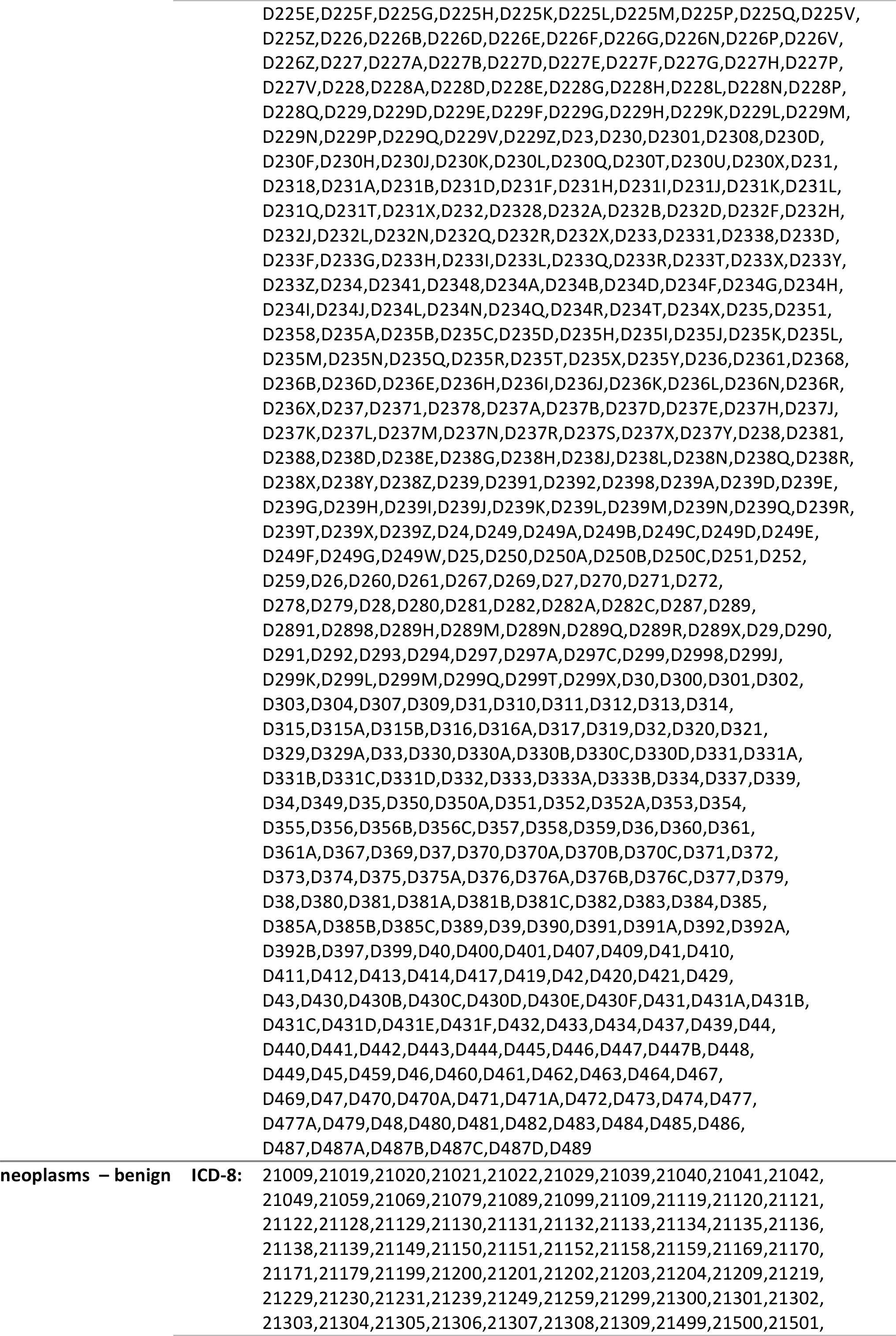
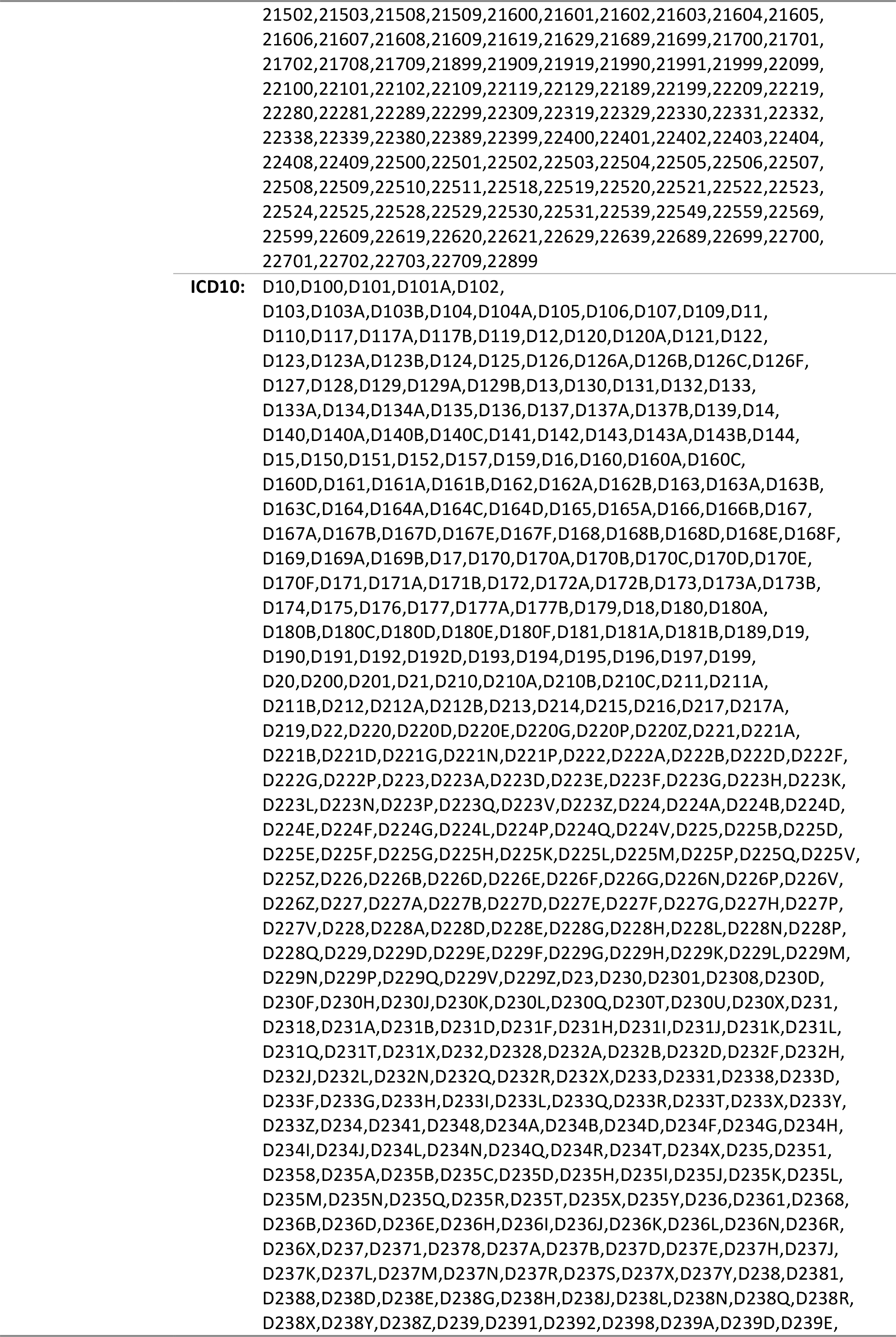
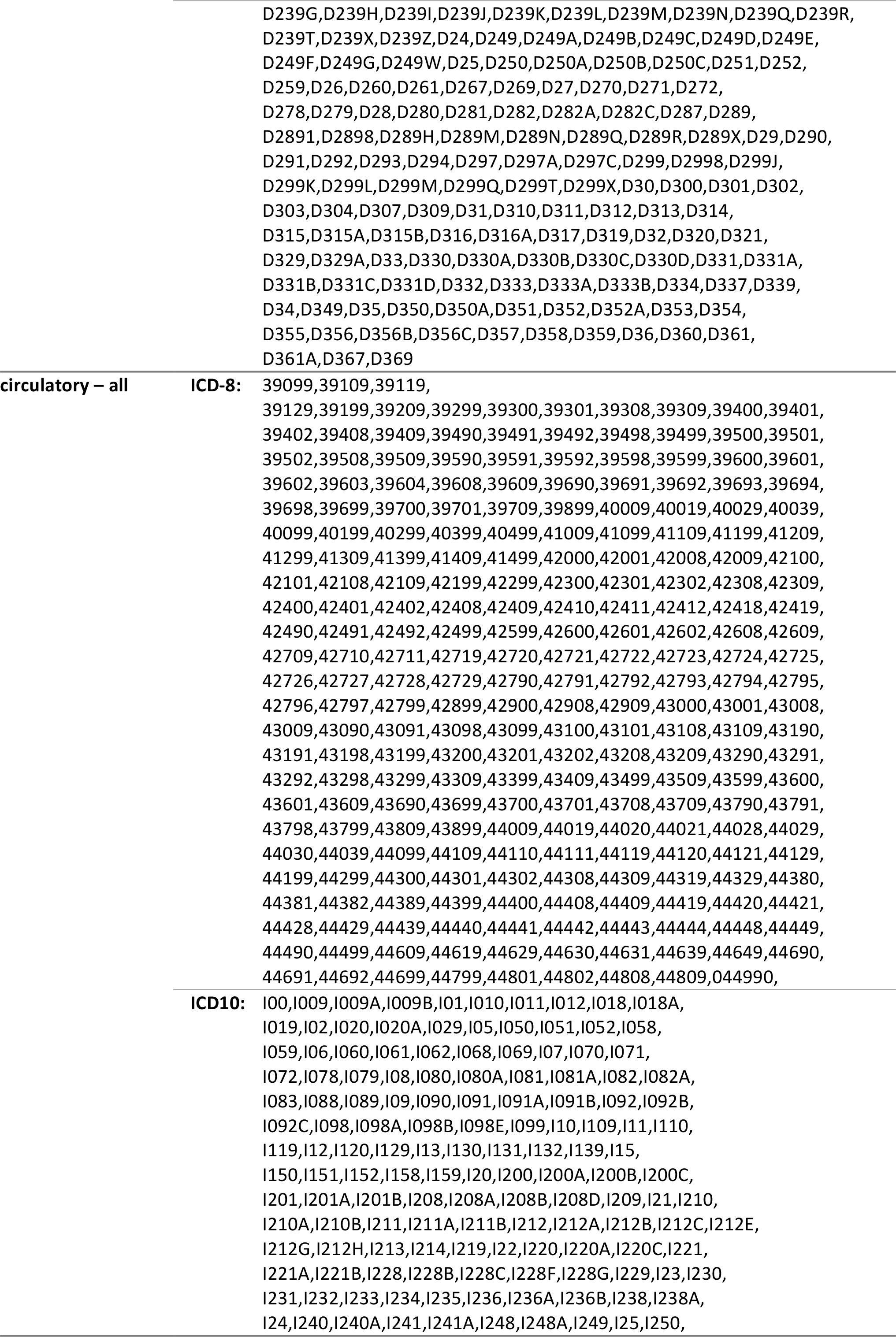
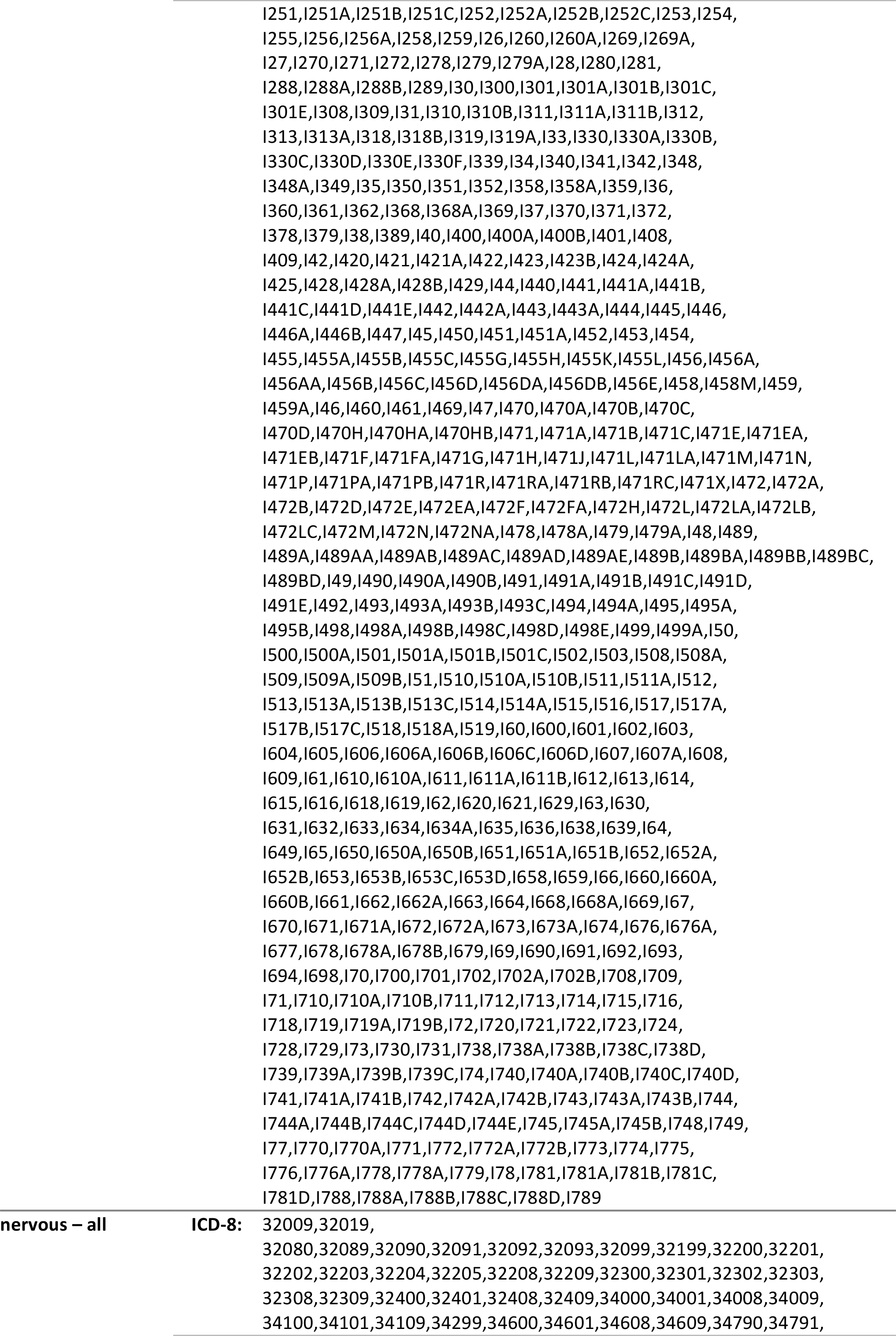
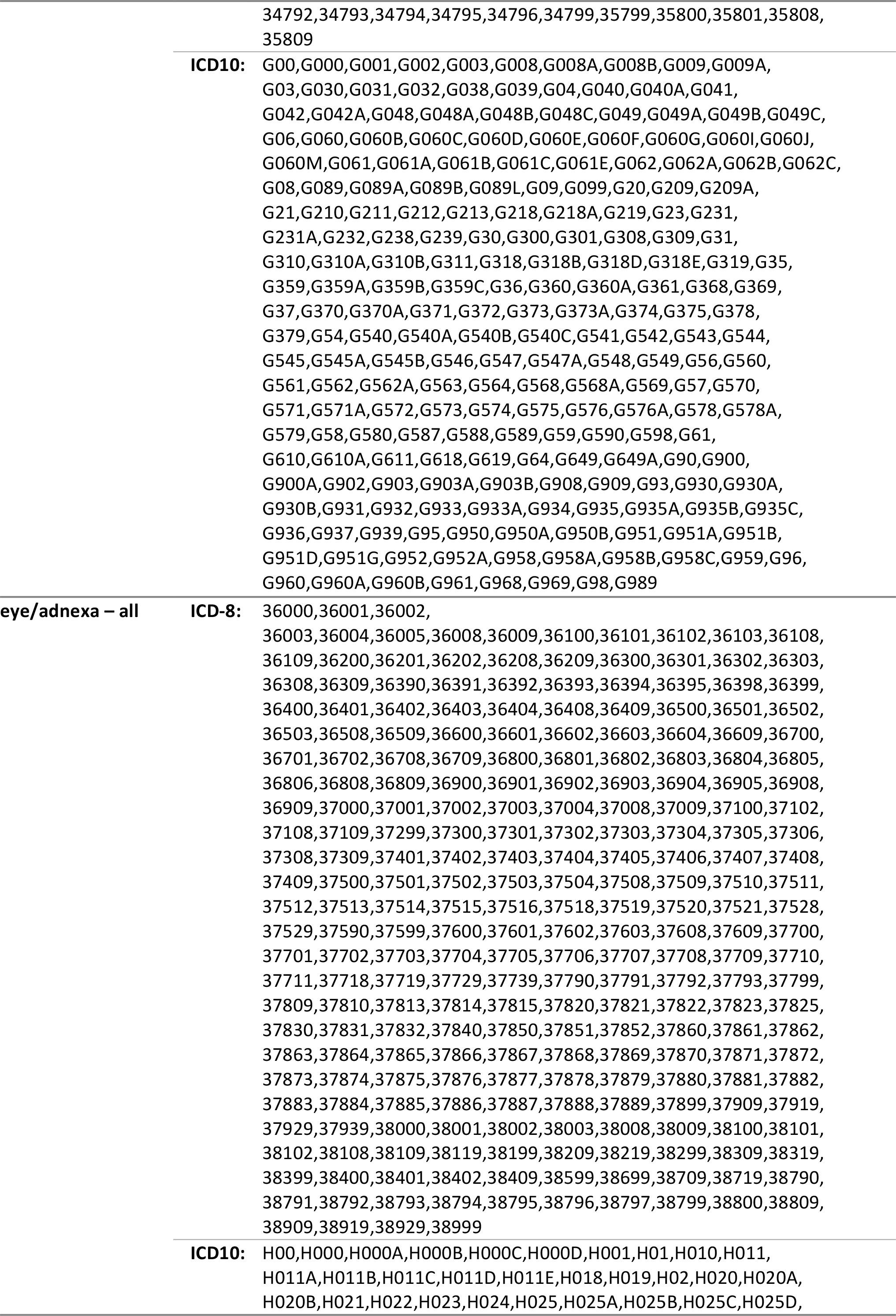
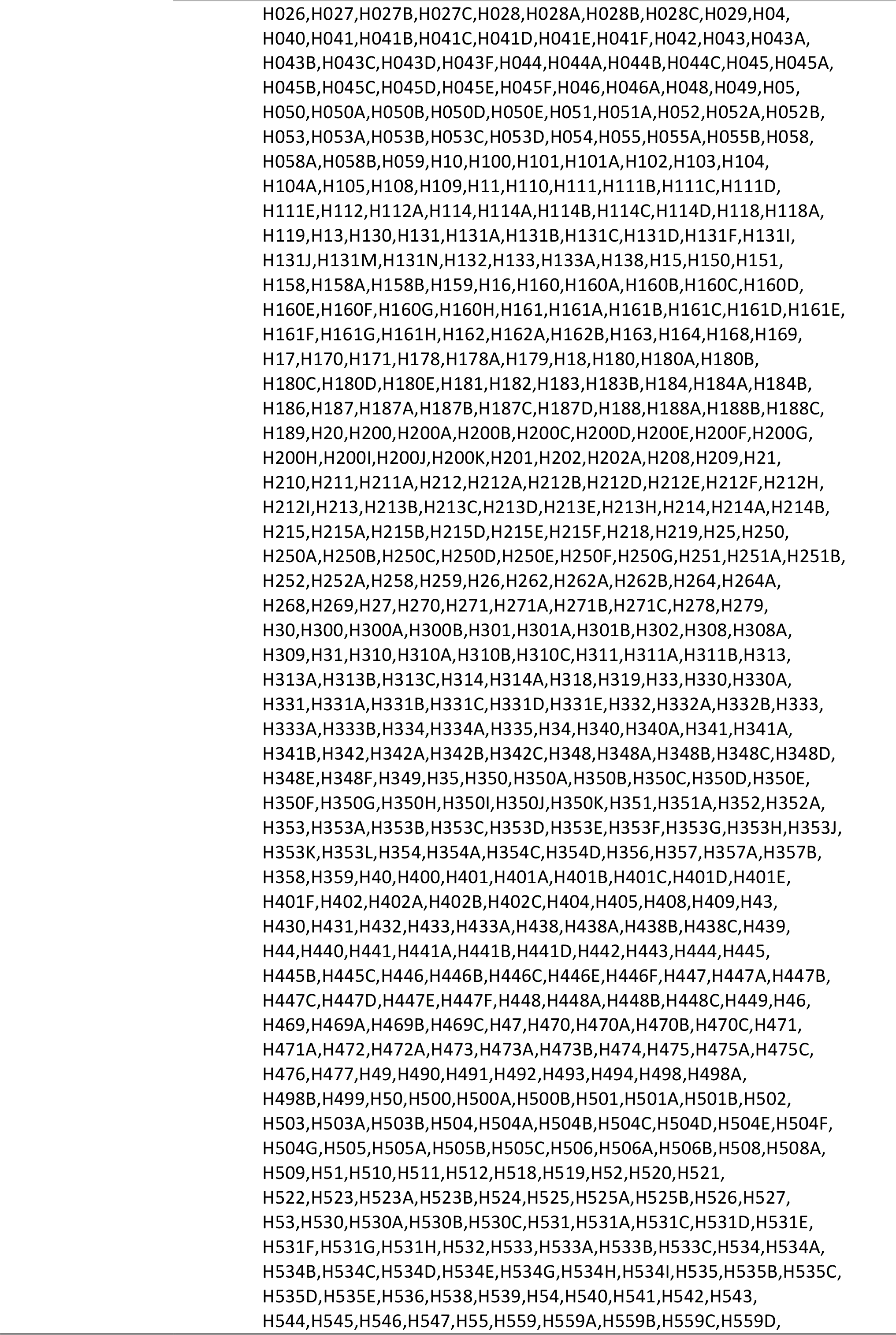
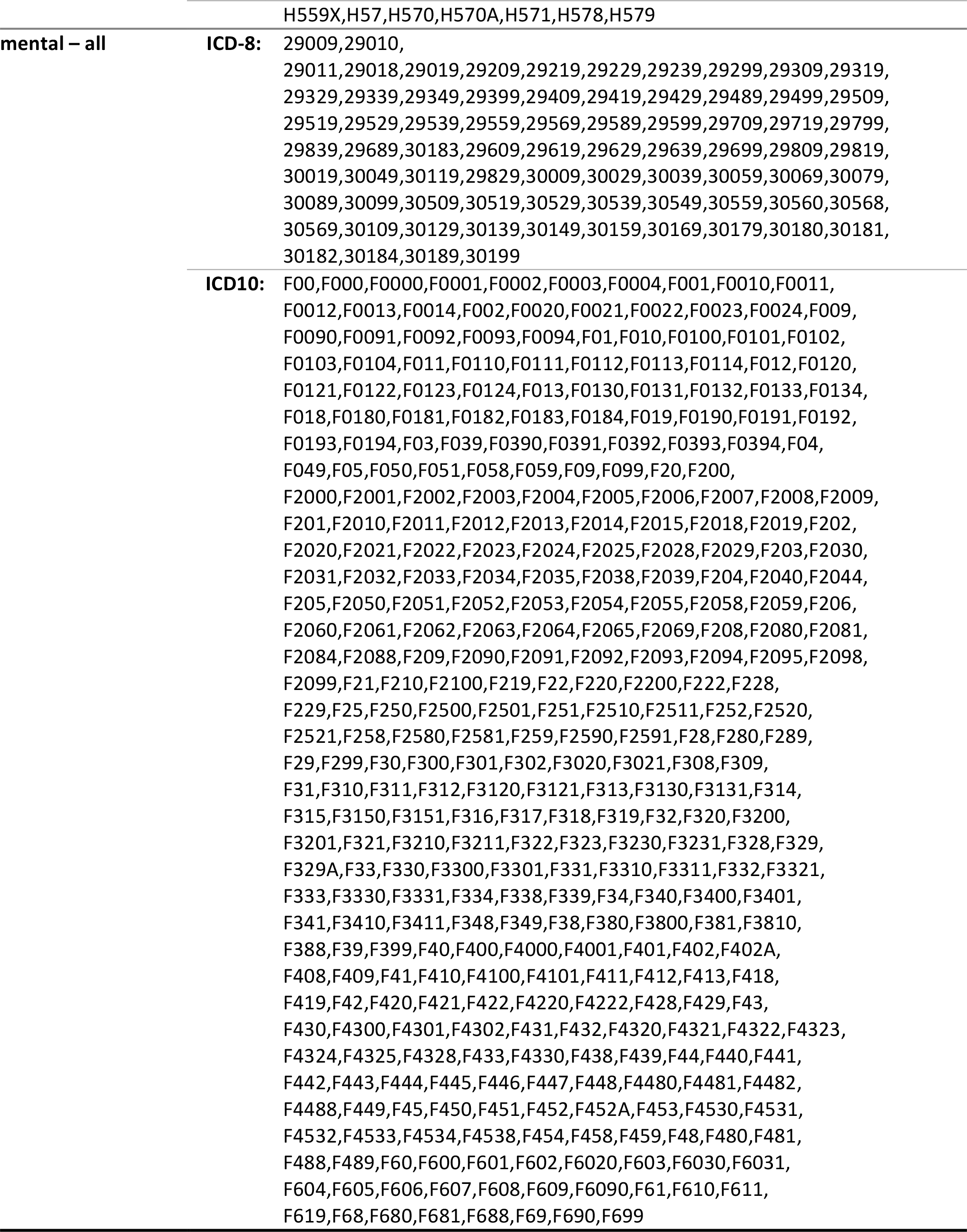
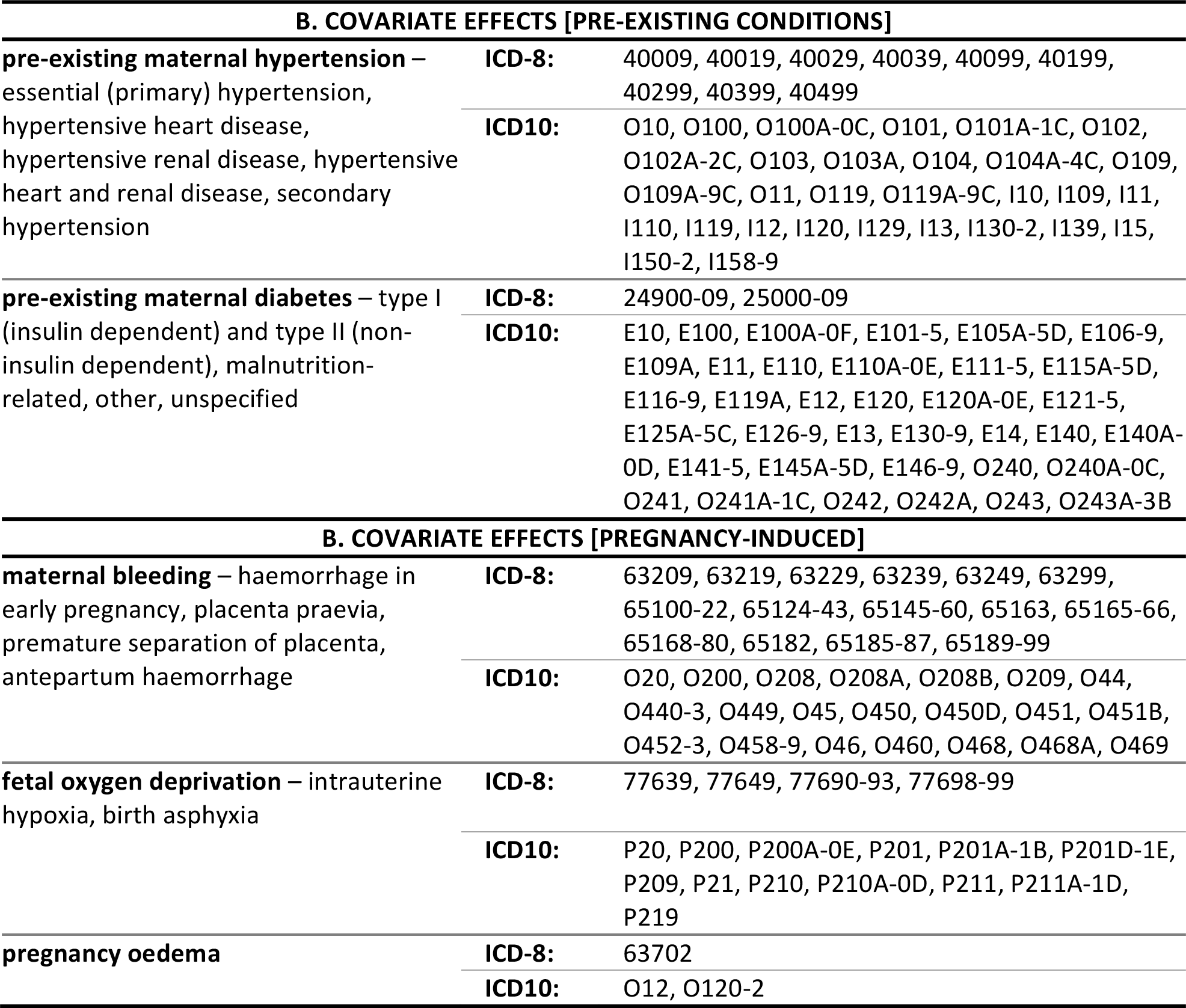
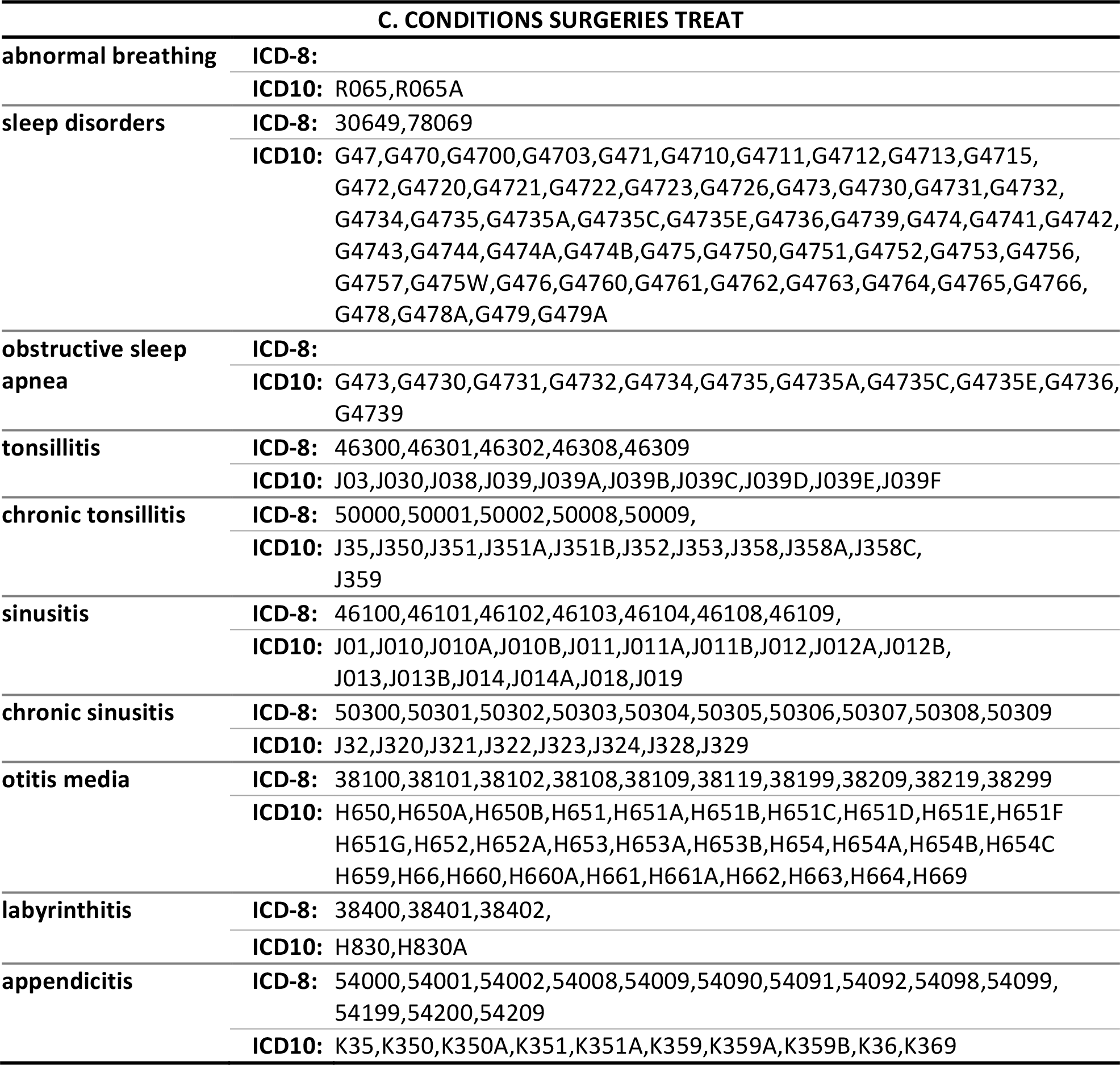
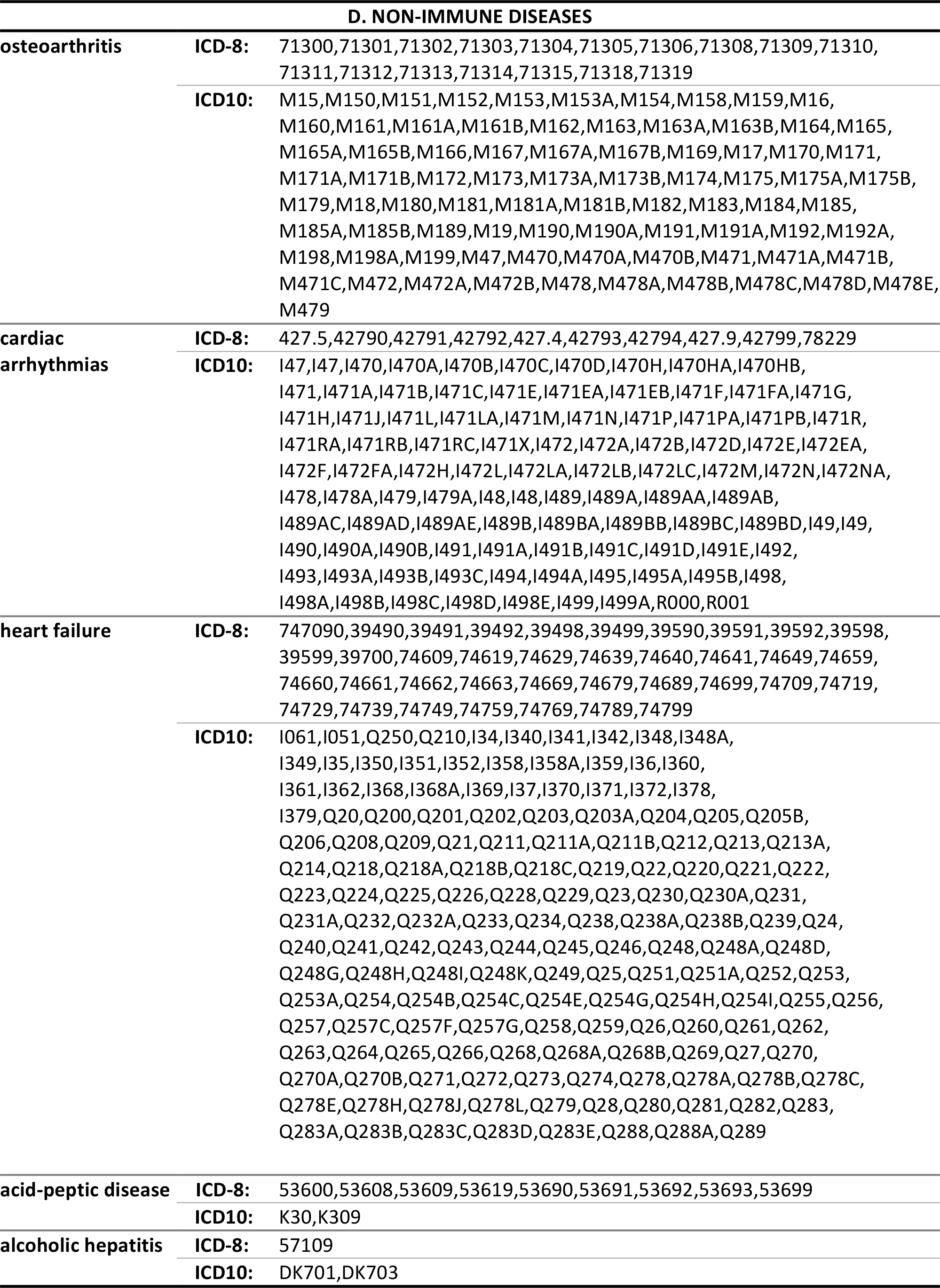
**ICD-8 and ICD-10 classification codes used for A. main diseases tested, B. covariate effects, C. conditions surgeries treat and D. nonimmune diseases.** Surgeries include adenotomy, tonsillectomy and adenotonsillectomy.

**Table S3.** **Relative risk (RR) for non-immune diseases up to age 30 depending on whether any of the three surgeries occurred between birth to 9 years of age.**

|  | adenoidectomy | tonsillectomy | adenotonsillectomy |
| --- | --- | --- | --- |
| Disease | RR | RR | RR |
| osteoarthritis | 1.04(0.85-1.29) | 1.03(0.75-1.41) | 0.87(0.71-1.07) |
| cardiac arrhythmias | 0.99(0.82-1.21) | 0.99(0.75-1.31) | 1.17(0.99-1.37) |
| heart failure | 1.24(0.95-1.61) | 1.28(0.92-1.78) | 1.15(0.93-1.42) |
| acid-peptic disease | 1.17(1.01-1.36) | 1.36(1.10-1.68) | 1.24(1.08-1.41) |
| alcoholic hepatitis | 1.92(0.23-15.84) | - | 5.78(1.21-27.59) |

- RR exclusion: Missing relative risks (RR) occur when the exponentiated coefficient for the Cox regression was infinite or there was insufficient power for hypothesis testing (see methods) - RR p-values: None of the relative risks above were significant after Bonferroni correction for 14 analyses (i.e. $\alpha=0.05/14= 0.00357$ )

**Table S4.**
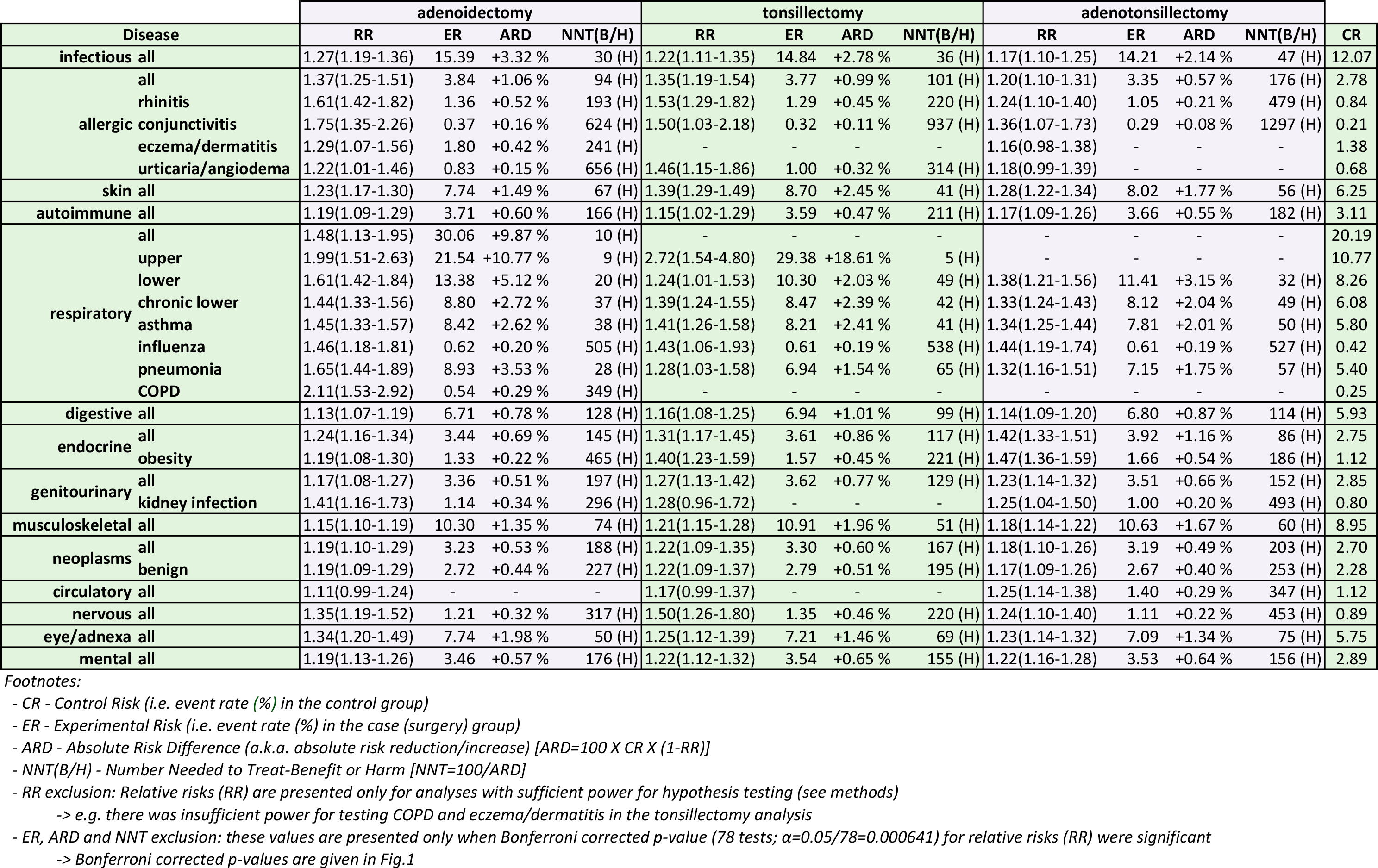
**Accompanying values for Fig. 1 including relative risks (RR), absolute risk difference (ARD), number needed to treat (NNT(B/H) or benefit/harm)**

**Table S5.**
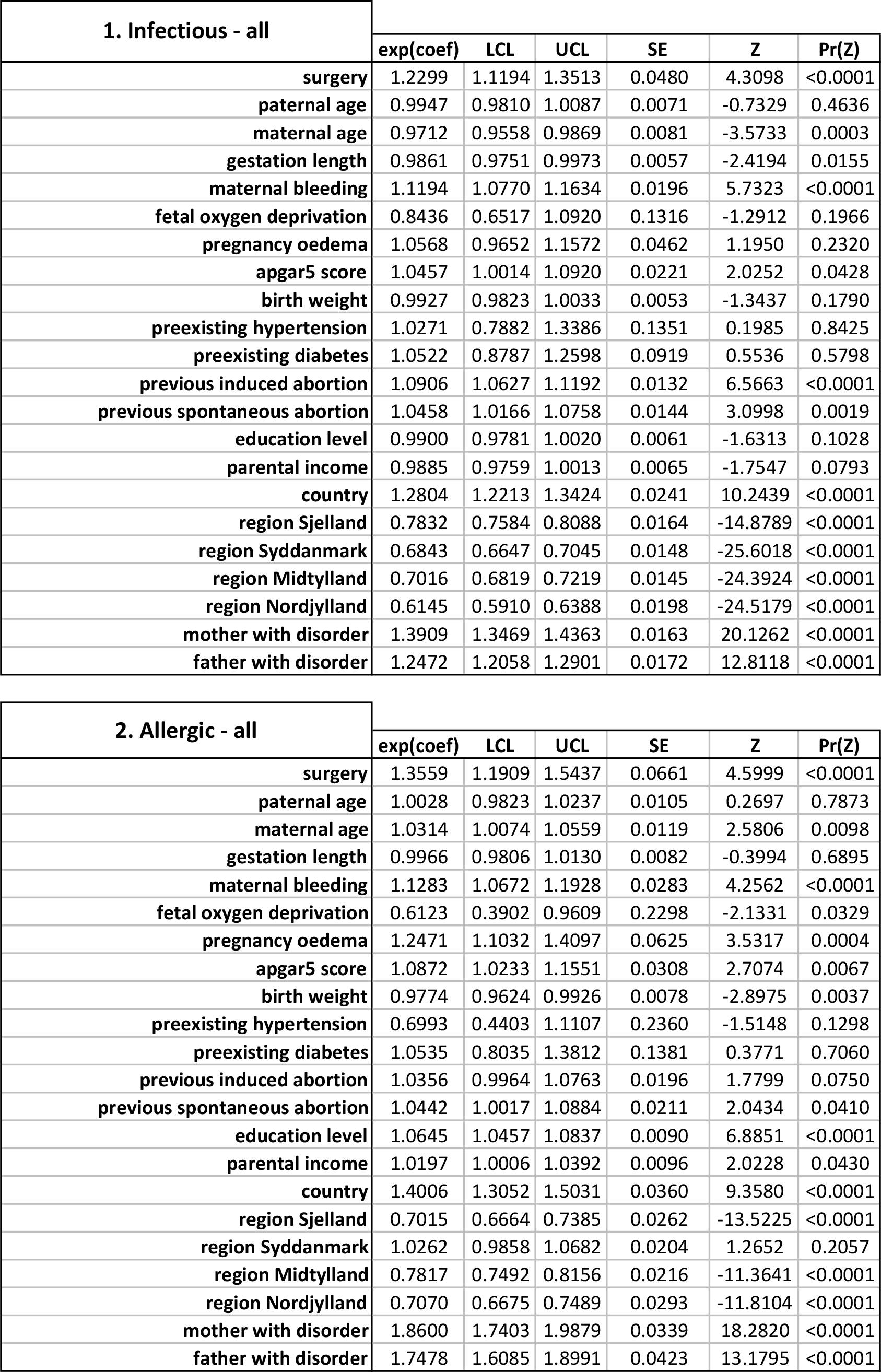
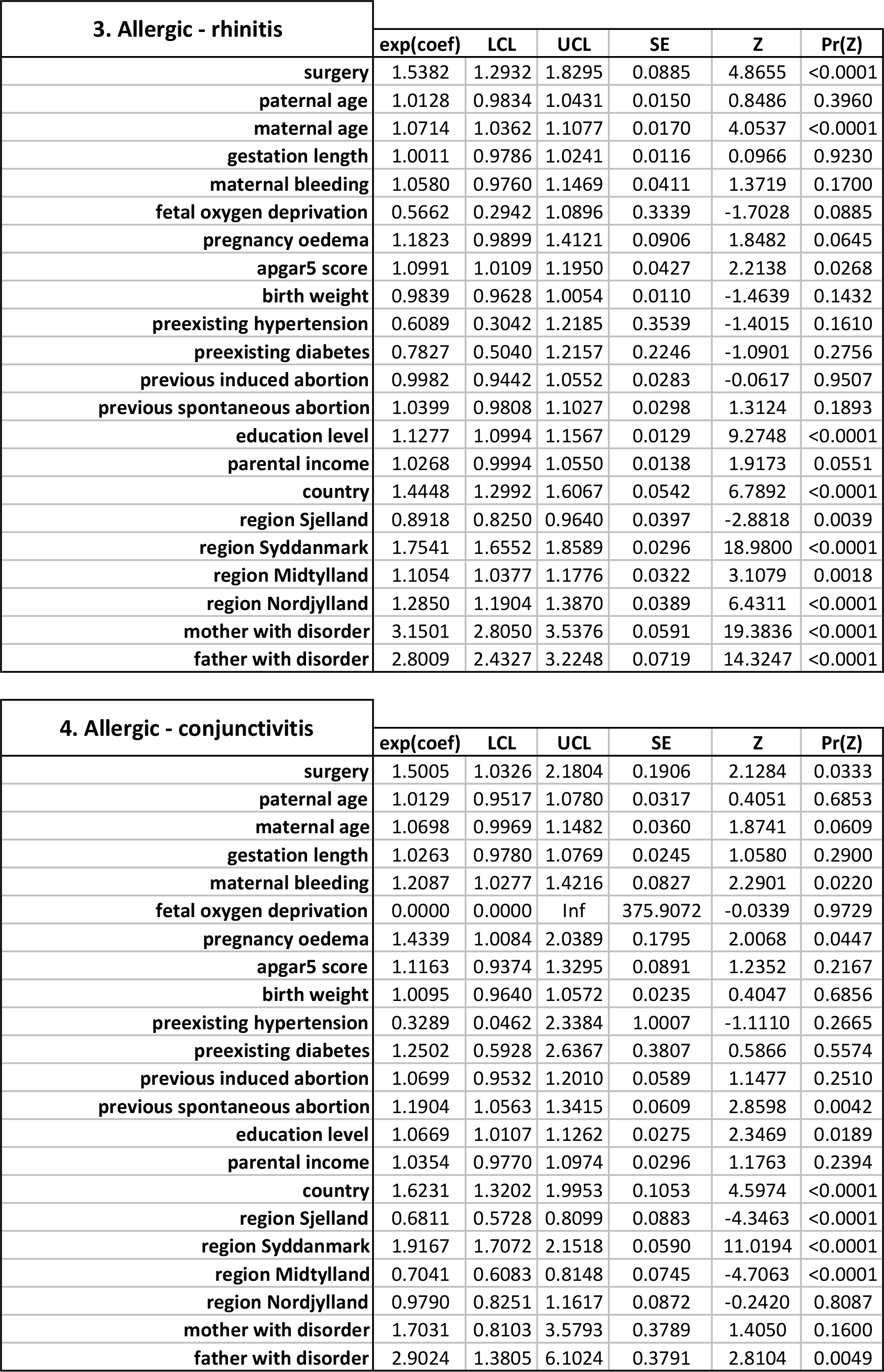
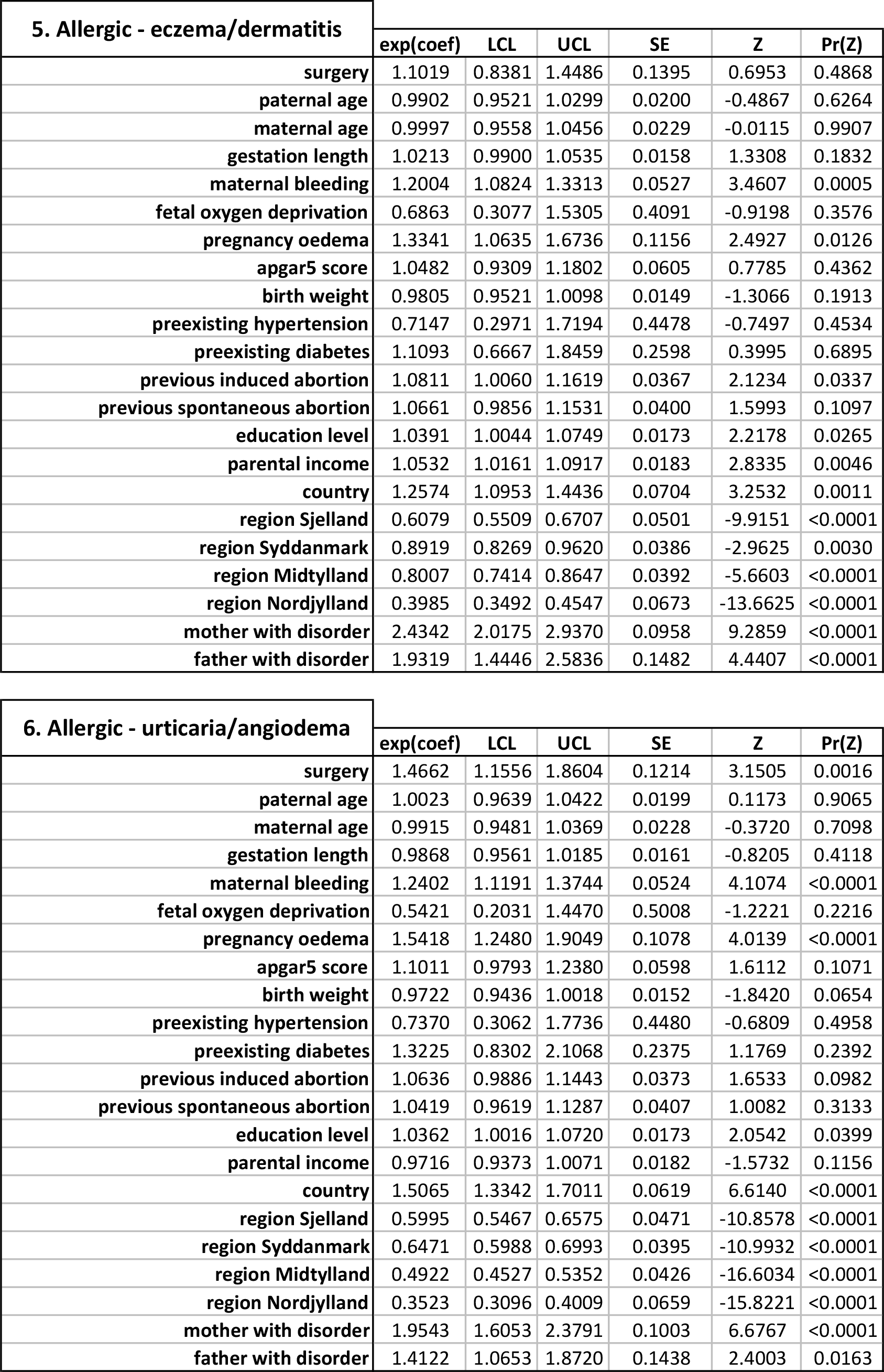
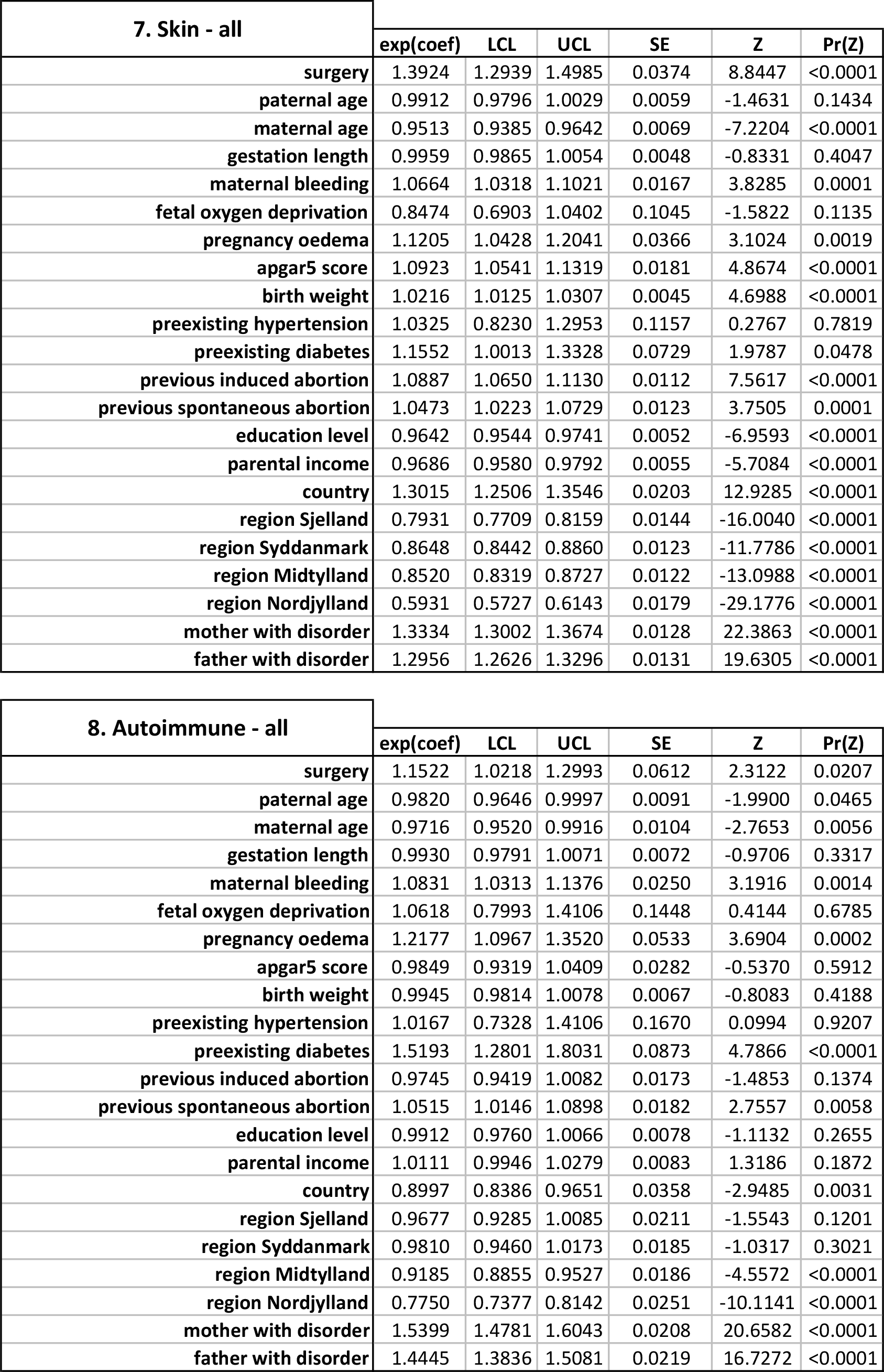
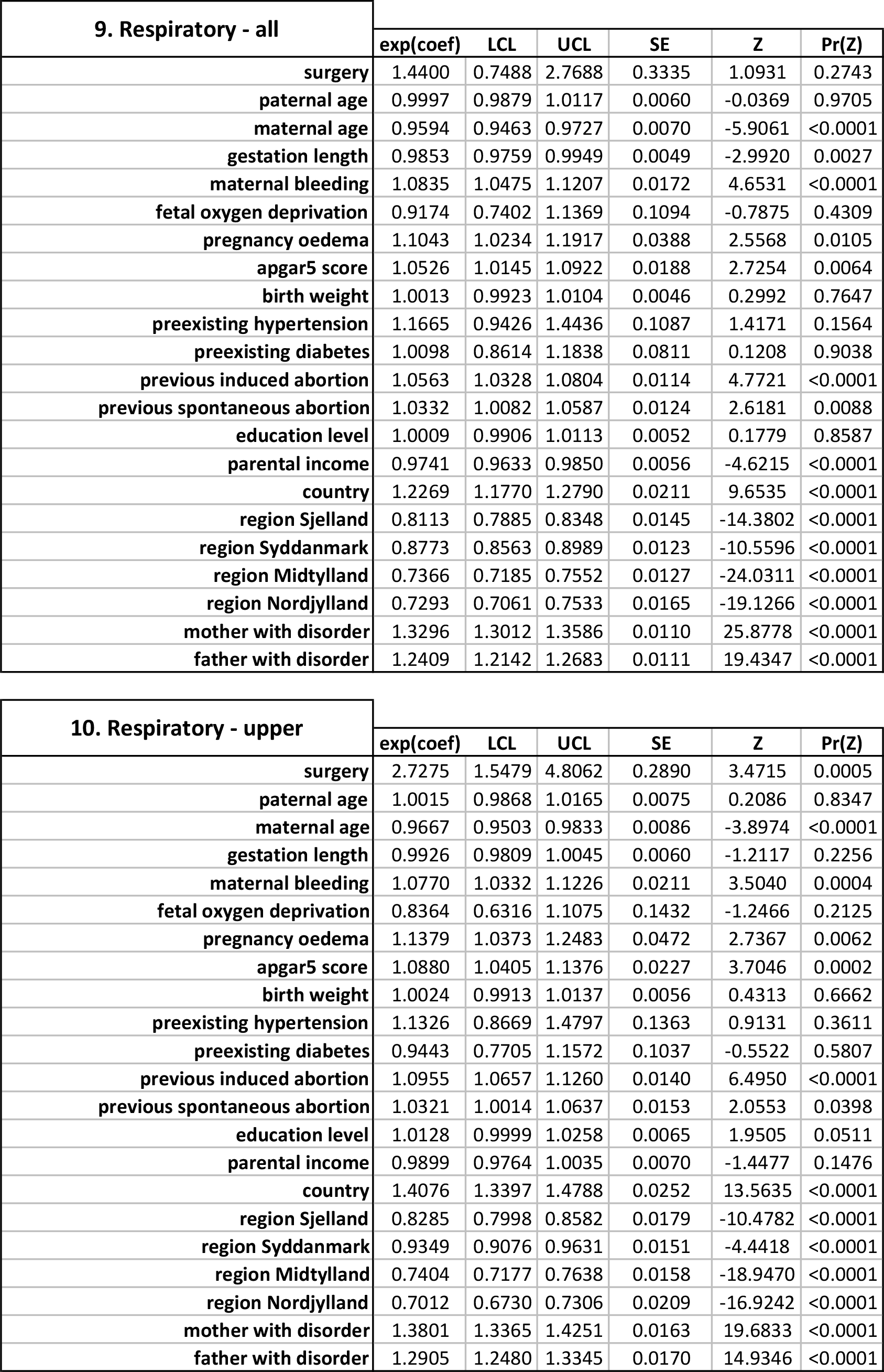
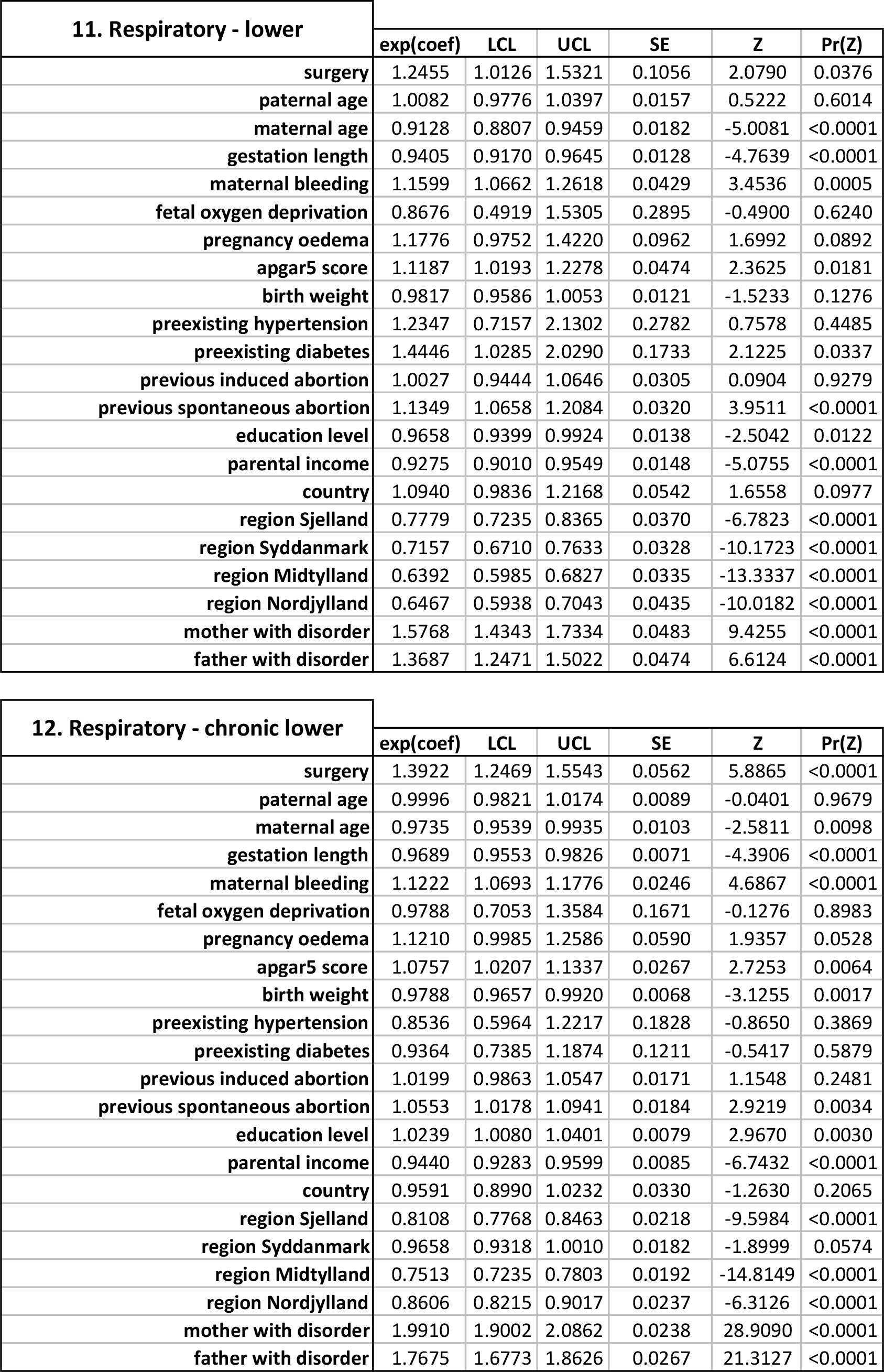
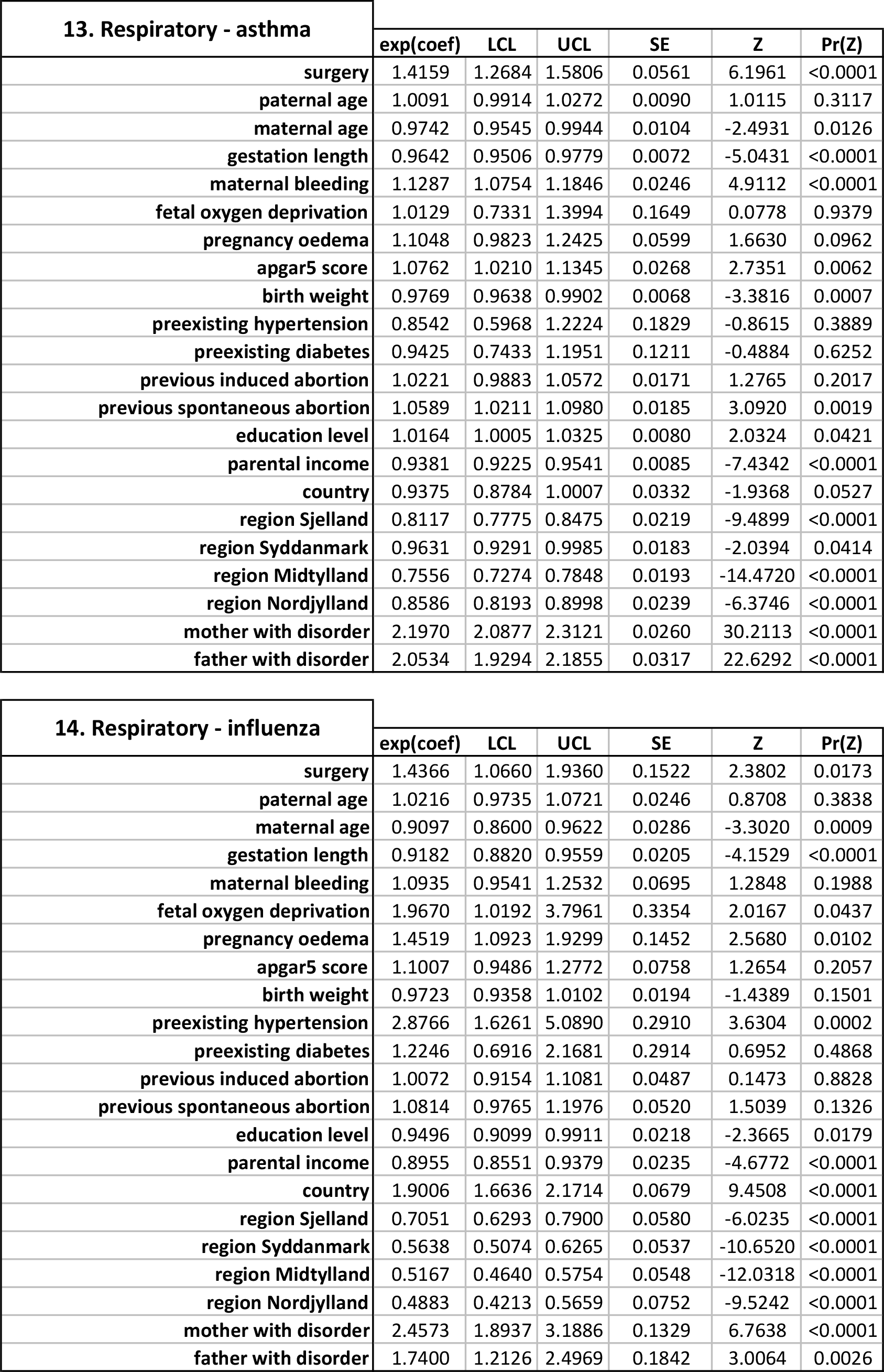
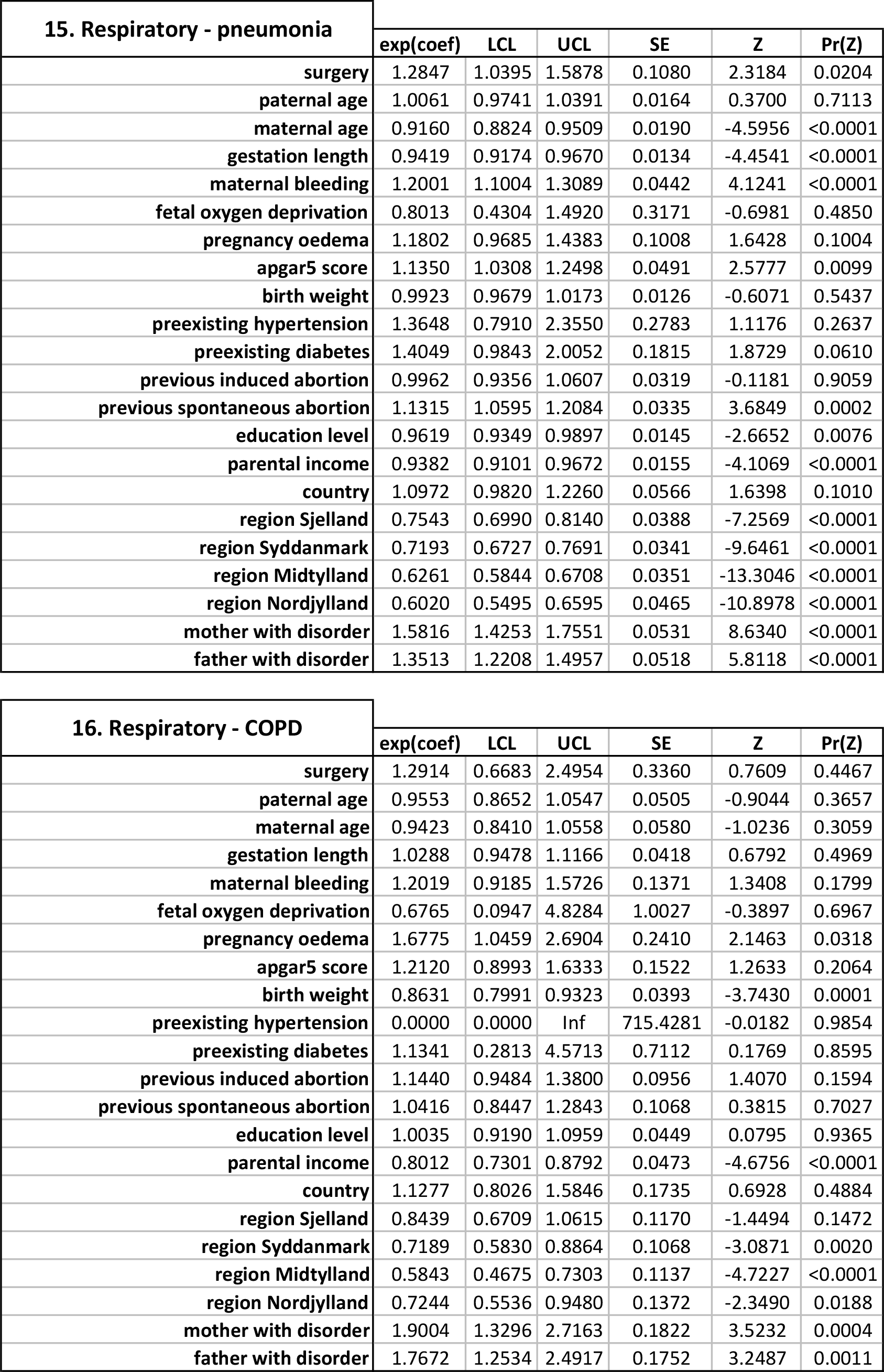
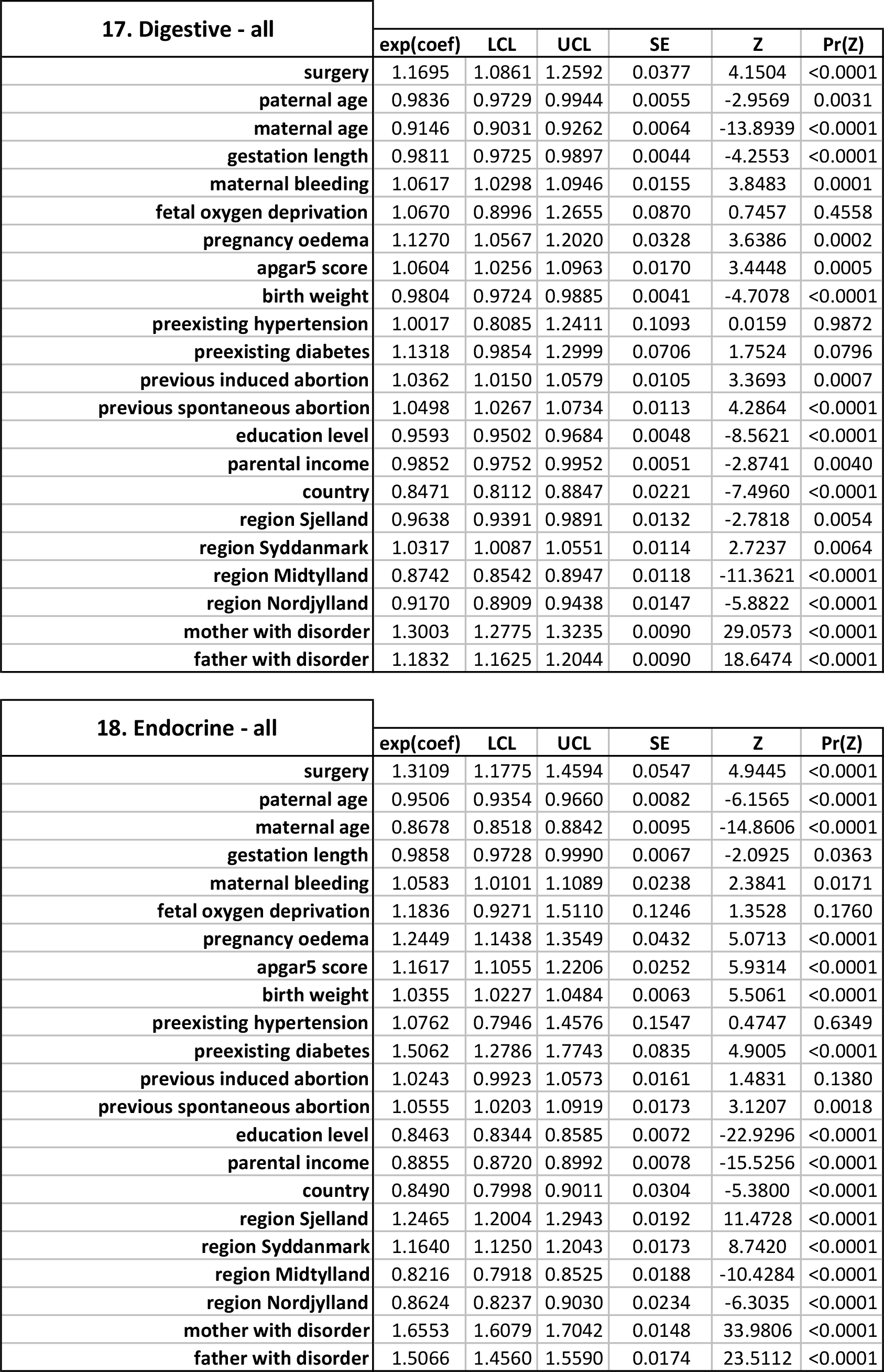
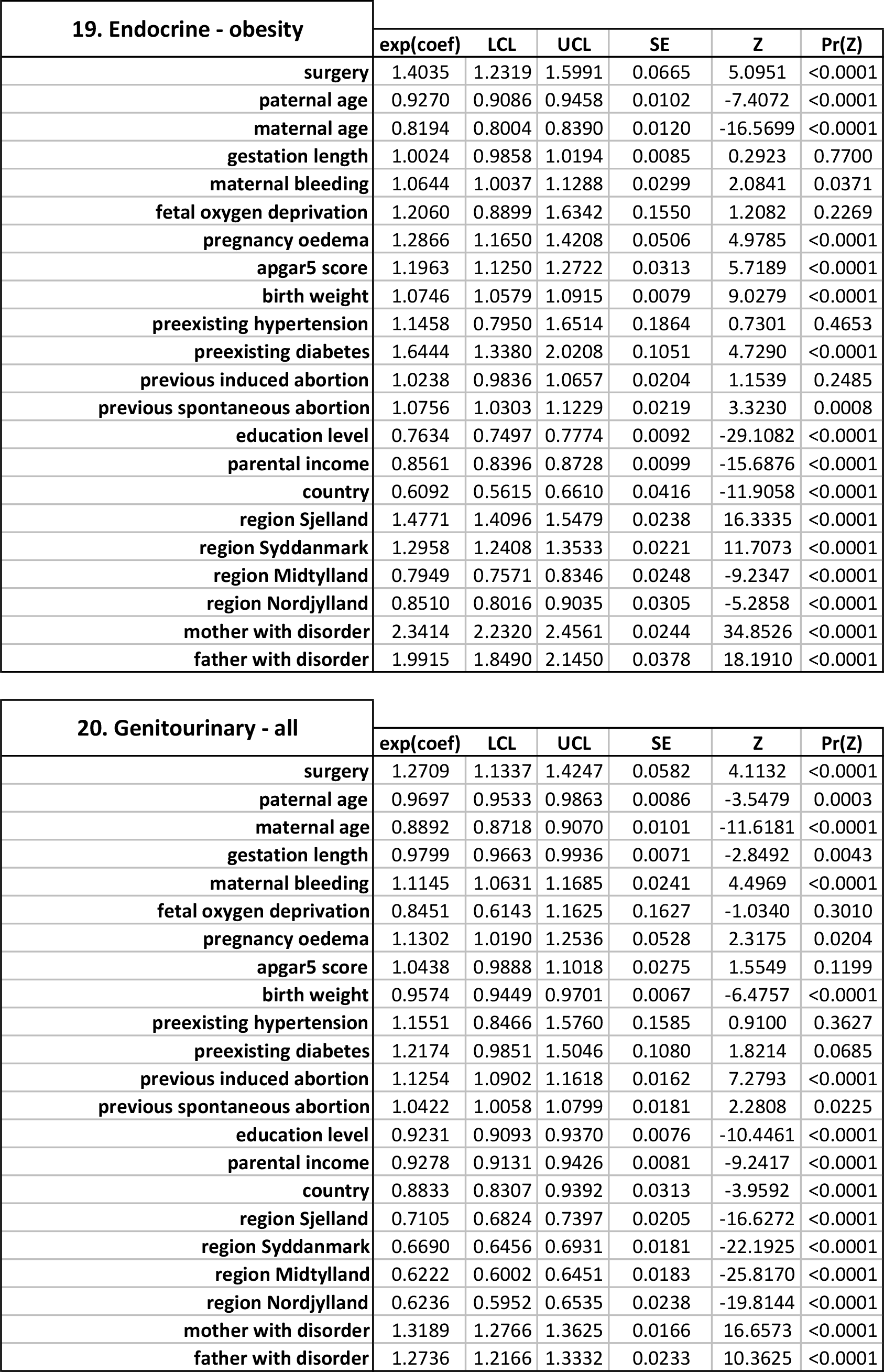
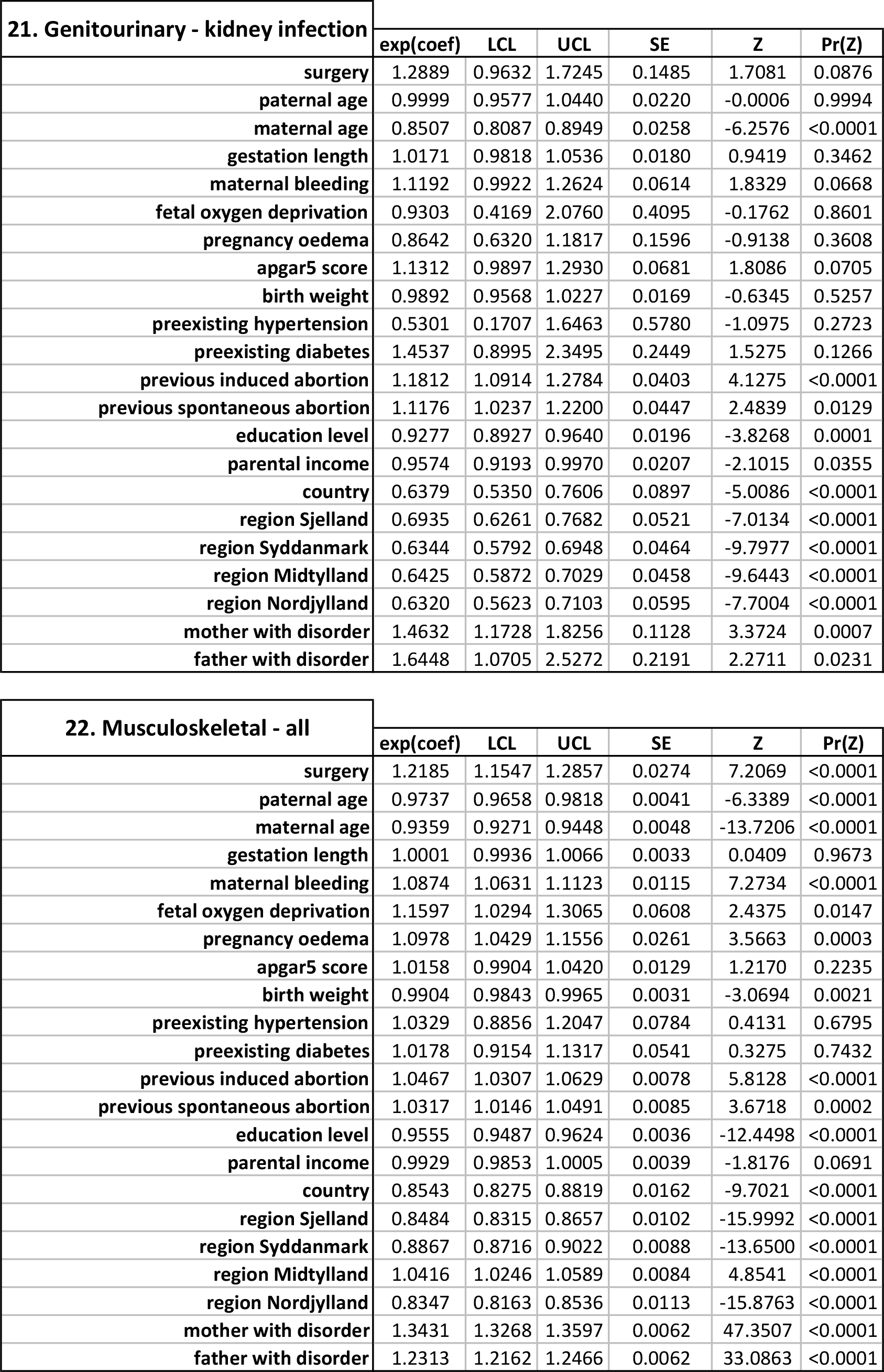
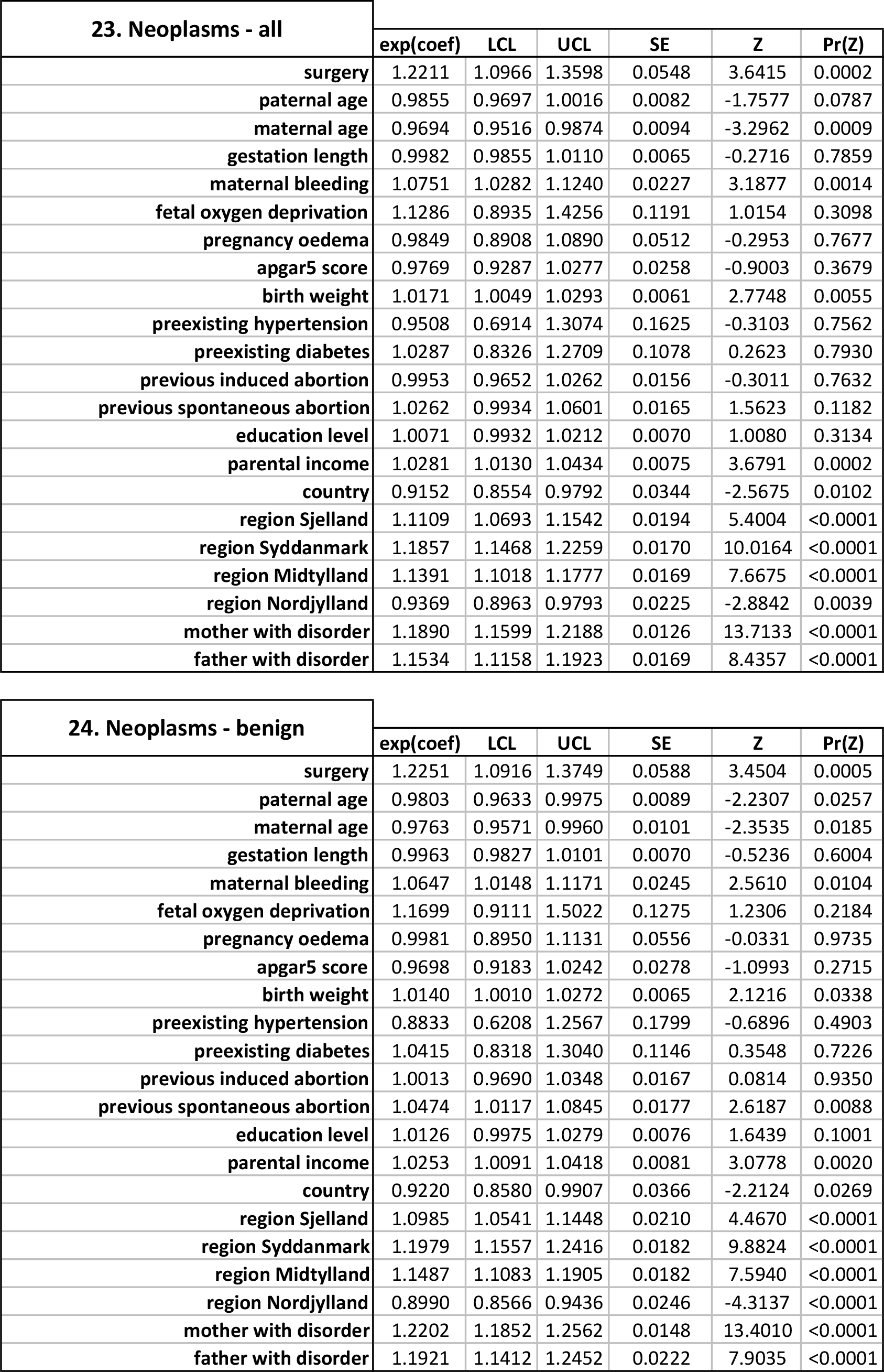
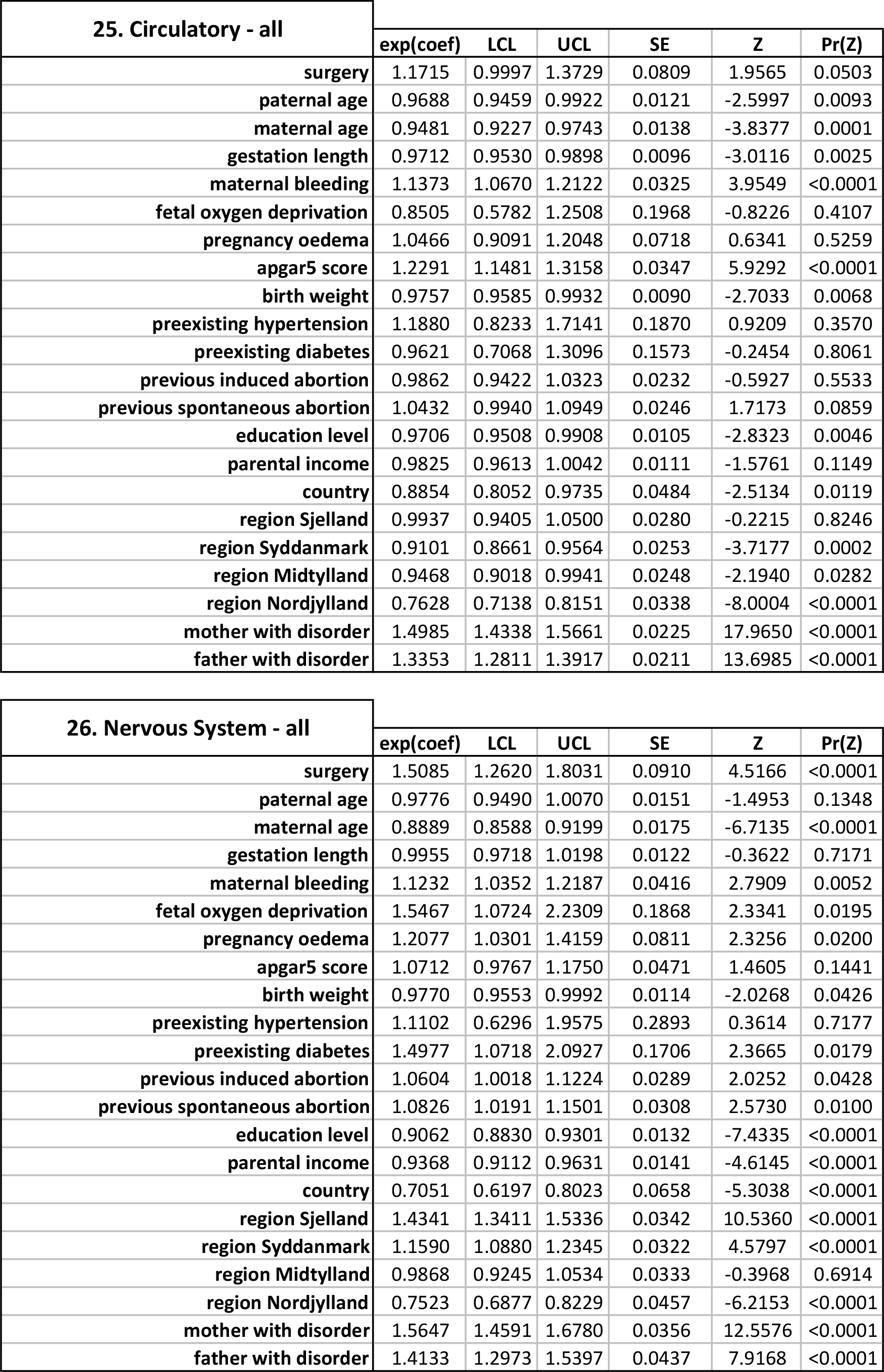
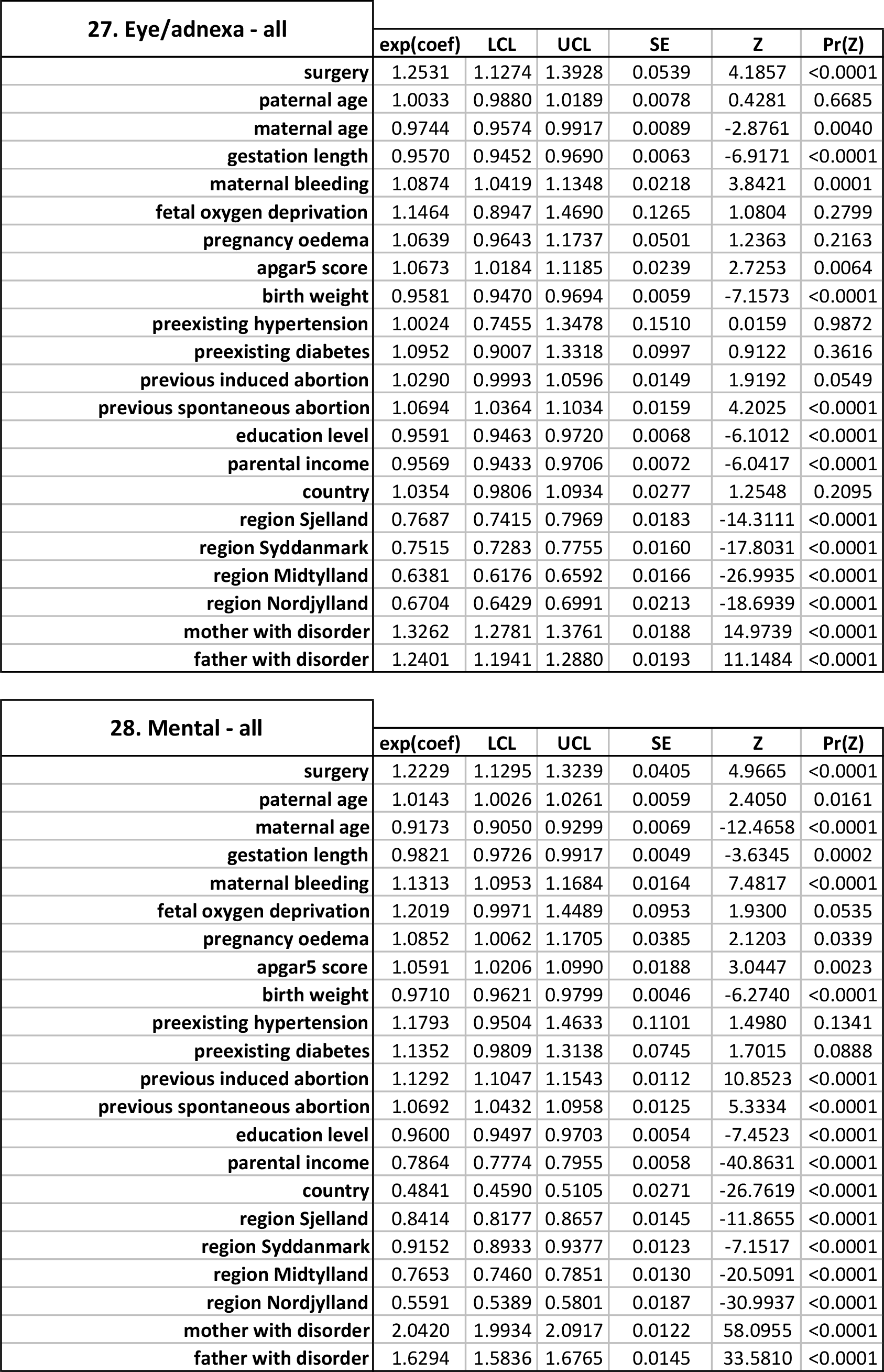
**Full Cox regression outputs - impact of tonsillectomy on risk of later disease.**

| **1. Infectious - all** | **exp(coef)** | **LCL** | **UCL** | **SE** | **Z** | **Pr(Z)** |
| --- | --- | --- | --- | --- | --- | --- |
| surgery | 1.2299 | 1.1194 | 1.3513 | 0.0480 | 4.3098 | <0.0001 |
| paternal age | 0.9947 | 0.9810 | 1.0087 | 0.0071 | -0.7329 | 0.4636 |
| maternal age | 0.9712 | 0.9558 | 0.9869 | 0.0081 | -3.5733 | 0.0003 |
| gestation length | 0.9861 | 0.9751 | 0.9973 | 0.0057 | -2.4194 | 0.0155 |
| maternal bleeding | 1.1194 | 1.0770 | 1.1634 | 0.0196 | 5.7323 | <0.0001 |
| fetal oxygen deprivation | 0.8436 | 0.6517 | 1.0920 | 0.1316 | -1.2912 | 0.1966 |
| pregnancy oedema | 1.0568 | 0.9652 | 1.1572 | 0.0462 | 1.1950 | 0.2320 |
| apgar5 score | 1.0457 | 1.0014 | 1.0920 | 0.0221 | 2.0252 | 0.0428 |
| birth weight | 0.9927 | 0.9823 | 1.0033 | 0.0053 | -1.3437 | 0.1790 |
| preexisting hypertension | 1.0271 | 0.7882 | 1.3386 | 0.1351 | 0.1985 | 0.8425 |
| preexisting diabetes | 1.0522 | 0.8787 | 1.2598 | 0.0919 | 0.5536 | 0.5798 |
| previous induced abortion | 1.0906 | 1.0627 | 1.1192 | 0.0132 | 6.5663 | <0.0001 |
| previous spontaneous abortion | 1.0458 | 1.0166 | 1.0758 | 0.0144 | 3.0998 | 0.0019 |
| education level | 0.9900 | 0.9781 | 1.0020 | 0.0061 | -1.6313 | 0.1028 |
| parental income | 0.9885 | 0.9759 | 1.0013 | 0.0065 | -1.7547 | 0.0793 |
| country | 1.2804 | 1.2213 | 1.3424 | 0.0241 | 10.2439 | <0.0001 |
| region Sjælland | 0.7832 | 0.7584 | 0.8088 | 0.0164 | -14.8789 | <0.0001 |
| region Syddanmark | 0.6843 | 0.6647 | 0.7045 | 0.0148 | -25.6018 | <0.0001 |
| region Midtjylland | 0.7016 | 0.6819 | 0.7219 | 0.0145 | -24.3924 | <0.0001 |
| region Nordjylland | 0.6145 | 0.5910 | 0.6388 | 0.0198 | -24.5179 | <0.0001 |
| mother with disorder | 1.3909 | 1.3469 | 1.4363 | 0.0163 | 20.1262 | <0.0001 |
| father with disorder | 1.2472 | 1.2058 | 1.2901 | 0.0172 | 12.8118 | <0.0001 |

| **2. Allergic - all** | **exp(coef)** | **LCL** | **UCL** | **SE** | **Z** | **Pr(Z)** |
| --- | --- | --- | --- | --- | --- | --- |
| surgery | 1.3559 | 1.1909 | 1.5437 | 0.0661 | 4.5999 | <0.0001 |
| paternal age | 1.0028 | 0.9823 | 1.0237 | 0.0105 | 0.2697 | 0.7873 |
| maternal age | 1.0314 | 1.0074 | 1.0559 | 0.0119 | 2.5806 | 0.0098 |
| gestation length | 0.9966 | 0.9806 | 1.0130 | 0.0082 | -0.3994 | 0.6895 |
| maternal bleeding | 1.1283 | 1.0672 | 1.1928 | 0.0283 | 4.2562 | <0.0001 |
| fetal oxygen deprivation | 0.6123 | 0.3902 | 0.9609 | 0.2298 | -2.1331 | 0.0329 |
| pregnancy oedema | 1.2471 | 1.1032 | 1.4097 | 0.0625 | 3.5317 | 0.0004 |
| apgar5 score | 1.0872 | 1.0233 | 1.1551 | 0.0308 | 2.7074 | 0.0067 |
| birth weight | 0.9774 | 0.9624 | 0.9926 | 0.0078 | -2.8975 | 0.0037 |
| preexisting hypertension | 0.6993 | 0.4403 | 1.1107 | 0.2360 | -1.5148 | 0.1298 |
| preexisting diabetes | 1.0535 | 0.8035 | 1.3812 | 0.1381 | 0.3771 | 0.7060 |
| previous induced abortion | 1.0356 | 0.9964 | 1.0763 | 0.0196 | 1.7799 | 0.0750 |
| previous spontaneous abortion | 1.0442 | 1.0017 | 1.0884 | 0.0211 | 2.0434 | 0.0410 |
| education level | 1.0645 | 1.0457 | 1.0837 | 0.0090 | 6.8851 | <0.0001 |
| parental income | 1.0197 | 1.0006 | 1.0392 | 0.0096 | 2.0228 | 0.0430 |
| country | 1.4006 | 1.3052 | 1.5031 | 0.0360 | 9.3580 | <0.0001 |
| region Sjælland | 0.7015 | 0.6664 | 0.7385 | 0.0262 | -13.5225 | <0.0001 |
| region Syddanmark | 1.0262 | 0.9858 | 1.0682 | 0.0204 | 1.2652 | 0.2057 |
| region Midtjylland | 0.7817 | 0.7492 | 0.8156 | 0.0216 | -11.3641 | <0.0001 |
| region Nordjylland | 0.7070 | 0.6675 | 0.7489 | 0.0293 | -11.8104 | <0.0001 |
| mother with disorder | 1.8600 | 1.7403 | 1.9879 | 0.0339 | 18.2820 | <0.0001 |
| father with disorder | 1.7478 | 1.6085 | 1.8991 | 0.0423 | 13.1795 | <0.0001 |

Table S5
| 3. Allergic - rhinitis | exp(coef) | LCL | UCL | SE | Z | Pr(Z) |
| --- | --- | --- | --- | --- | --- | --- |
| surgery | 1.5382 | 1.2932 | 1.8295 | 0.0885 | 4.8655 | <0.0001 |
| paternal age | 1.0128 | 0.9834 | 1.0431 | 0.0150 | 0.8486 | 0.3960 |
| maternal age | 1.0714 | 1.0362 | 1.1077 | 0.0170 | 4.0537 | <0.0001 |
| gestation length | 1.0011 | 0.9786 | 1.0241 | 0.0116 | 0.0966 | 0.9230 |
| maternal bleeding | 1.0580 | 0.9760 | 1.1469 | 0.0411 | 1.3719 | 0.1700 |
| fetal oxygen deprivation | 0.5662 | 0.2942 | 1.0896 | 0.3339 | -1.7028 | 0.0885 |
| pregnancy oedema | 1.1823 | 0.9899 | 1.4121 | 0.0906 | 1.8482 | 0.0645 |
| apgar5 score | 1.0991 | 1.0109 | 1.1950 | 0.0427 | 2.2138 | 0.0268 |
| birth weight | 0.9839 | 0.9628 | 1.0054 | 0.0110 | -1.4639 | 0.1432 |
| preexisting hypertension | 0.6089 | 0.3042 | 1.2185 | 0.3539 | -1.4015 | 0.1610 |
| preexisting diabetes | 0.7827 | 0.5040 | 1.2157 | 0.2246 | -1.0901 | 0.2756 |
| previous induced abortion | 0.9982 | 0.9442 | 1.0552 | 0.0283 | -0.0617 | 0.9507 |
| previous spontaneous abortion | 1.0399 | 0.9808 | 1.1027 | 0.0298 | 1.3124 | 0.1893 |
| education level | 1.1277 | 1.0994 | 1.1567 | 0.0129 | 9.2748 | <0.0001 |
| parental income | 1.0268 | 0.9994 | 1.0550 | 0.0138 | 1.9173 | 0.0551 |
| country | 1.4448 | 1.2992 | 1.6067 | 0.0542 | 6.7892 | <0.0001 |
| region Sjælland | 0.8918 | 0.8250 | 0.9640 | 0.0397 | -2.8818 | 0.0039 |
| region Syddanmark | 1.7541 | 1.6552 | 1.8589 | 0.0296 | 18.9800 | <0.0001 |
| region Midtjylland | 1.1054 | 1.0377 | 1.1776 | 0.0322 | 3.1079 | 0.0018 |
| region Nordjylland | 1.2850 | 1.1904 | 1.3870 | 0.0389 | 6.4311 | <0.0001 |
| mother with disorder | 3.1501 | 2.8050 | 3.5376 | 0.0591 | 19.3836 | <0.0001 |
| father with disorder | 2.8009 | 2.4327 | 3.2248 | 0.0719 | 14.3247 | <0.0001 |

| 4. Allergic - conjunctivitis | exp(coef) | LCL | UCL | SE | Z | Pr(Z) |
| --- | --- | --- | --- | --- | --- | --- |
| surgery | 1.5005 | 1.0326 | 2.1804 | 0.1906 | 2.1284 | 0.0333 |
| paternal age | 1.0129 | 0.9517 | 1.0780 | 0.0317 | 0.4051 | 0.6853 |
| maternal age | 1.0698 | 0.9969 | 1.1482 | 0.0360 | 1.8741 | 0.0609 |
| gestation length | 1.0263 | 0.9780 | 1.0769 | 0.0245 | 1.0580 | 0.2900 |
| maternal bleeding | 1.2087 | 1.0277 | 1.4216 | 0.0827 | 2.2901 | 0.0220 |
| fetal oxygen deprivation | 0.0000 | 0.0000 | Inf | 375.9072 | -0.0339 | 0.9729 |
| pregnancy oedema | 1.4339 | 1.0084 | 2.0389 | 0.1795 | 2.0068 | 0.0447 |
| apgar5 score | 1.1163 | 0.9374 | 1.3295 | 0.0891 | 1.2352 | 0.2167 |
| birth weight | 1.0095 | 0.9640 | 1.0572 | 0.0235 | 0.4047 | 0.6856 |
| preexisting hypertension | 0.3289 | 0.0462 | 2.3384 | 1.0007 | -1.1110 | 0.2665 |
| preexisting diabetes | 1.2502 | 0.5928 | 2.6367 | 0.3807 | 0.5866 | 0.5574 |
| previous induced abortion | 1.0699 | 0.9532 | 1.2010 | 0.0589 | 1.1477 | 0.2510 |
| previous spontaneous abortion | 1.1904 | 1.0563 | 1.3415 | 0.0609 | 2.8598 | 0.0042 |
| education level | 1.0669 | 1.0107 | 1.1262 | 0.0275 | 2.3469 | 0.0189 |
| parental income | 1.0354 | 0.9770 | 1.0974 | 0.0296 | 1.1763 | 0.2394 |
| country | 1.6231 | 1.3202 | 1.9953 | 0.1053 | 4.5974 | <0.0001 |
| region Sjælland | 0.6811 | 0.5728 | 0.8099 | 0.0883 | -4.3463 | <0.0001 |
| region Syddanmark | 1.9167 | 1.7072 | 2.1518 | 0.0590 | 11.0194 | <0.0001 |
| region Midtjylland | 0.7041 | 0.6083 | 0.8148 | 0.0745 | -4.7063 | <0.0001 |
| region Nordjylland | 0.9790 | 0.8251 | 1.1617 | 0.0872 | -0.2420 | 0.8087 |
| mother with disorder | 1.7031 | 0.8103 | 3.5793 | 0.3789 | 1.4050 | 0.1600 |
| father with disorder | 2.9024 | 1.3805 | 6.1024 | 0.3791 | 2.8104 | 0.0049 |

Table S5
| 5. Allergic - eczema/dermatitis | exp(coef) | LCL | UCL | SE | Z | Pr(Z) |
| --- | --- | --- | --- | --- | --- | --- |
| surgery | 1.1019 | 0.8381 | 1.4486 | 0.1395 | 0.6953 | 0.4868 |
| paternal age | 0.9902 | 0.9521 | 1.0299 | 0.0200 | -0.4867 | 0.6264 |
| maternal age | 0.9997 | 0.9558 | 1.0456 | 0.0229 | -0.0115 | 0.9907 |
| gestation length | 1.0213 | 0.9900 | 1.0535 | 0.0158 | 1.3308 | 0.1832 |
| maternal bleeding | 1.2004 | 1.0824 | 1.3313 | 0.0527 | 3.4607 | 0.0005 |
| fetal oxygen deprivation | 0.6863 | 0.3077 | 1.5305 | 0.4091 | -0.9198 | 0.3576 |
| pregnancy oedema | 1.3341 | 1.0635 | 1.6736 | 0.1156 | 2.4927 | 0.0126 |
| apgar5 score | 1.0482 | 0.9309 | 1.1802 | 0.0605 | 0.7785 | 0.4362 |
| birth weight | 0.9805 | 0.9521 | 1.0098 | 0.0149 | -1.3066 | 0.1913 |
| preexisting hypertension | 0.7147 | 0.2971 | 1.7194 | 0.4478 | -0.7497 | 0.4534 |
| preexisting diabetes | 1.1093 | 0.6667 | 1.8459 | 0.2598 | 0.3995 | 0.6895 |
| previous induced abortion | 1.0811 | 1.0060 | 1.1619 | 0.0367 | 2.1234 | 0.0337 |
| previous spontaneous abortion | 1.0661 | 0.9856 | 1.1531 | 0.0400 | 1.5993 | 0.1097 |
| education level | 1.0391 | 1.0044 | 1.0749 | 0.0173 | 2.2178 | 0.0265 |
| parental income | 1.0532 | 1.0161 | 1.0917 | 0.0183 | 2.8335 | 0.0046 |
| country | 1.2574 | 1.0953 | 1.4436 | 0.0704 | 3.2532 | 0.0011 |
| region Sjælland | 0.6079 | 0.5509 | 0.6707 | 0.0501 | -9.9151 | <0.0001 |
| region Syddanmark | 0.8919 | 0.8269 | 0.9620 | 0.0386 | -2.9625 | 0.0030 |
| region Midtjylland | 0.8007 | 0.7414 | 0.8647 | 0.0392 | -5.6603 | <0.0001 |
| region Nordjylland | 0.3985 | 0.3492 | 0.4547 | 0.0673 | -13.6625 | <0.0001 |
| mother with disorder | 2.4342 | 2.0175 | 2.9370 | 0.0958 | 9.2859 | <0.0001 |
| father with disorder | 1.9319 | 1.4446 | 2.5836 | 0.1482 | 4.4407 | <0.0001 |

| 6. Allergic - urticaria/angiodema | exp(coef) | LCL | UCL | SE | Z | Pr(Z) |
| --- | --- | --- | --- | --- | --- | --- |
| surgery | 1.4662 | 1.1556 | 1.8604 | 0.1214 | 3.1505 | 0.0016 |
| paternal age | 1.0023 | 0.9639 | 1.0422 | 0.0199 | 0.1173 | 0.9065 |
| maternal age | 0.9915 | 0.9481 | 1.0369 | 0.0228 | -0.3720 | 0.7098 |
| gestation length | 0.9868 | 0.9561 | 1.0185 | 0.0161 | -0.8205 | 0.4118 |
| maternal bleeding | 1.2402 | 1.1191 | 1.3744 | 0.0524 | 4.1074 | <0.0001 |
| fetal oxygen deprivation | 0.5421 | 0.2031 | 1.4470 | 0.5008 | -1.2221 | 0.2216 |
| pregnancy oedema | 1.5418 | 1.2480 | 1.9049 | 0.1078 | 4.0139 | <0.0001 |
| apgar5 score | 1.1011 | 0.9793 | 1.2380 | 0.0598 | 1.6112 | 0.1071 |
| birth weight | 0.9722 | 0.9436 | 1.0018 | 0.0152 | -1.8420 | 0.0654 |
| preexisting hypertension | 0.7370 | 0.3062 | 1.7736 | 0.4480 | -0.6809 | 0.4958 |
| preexisting diabetes | 1.3225 | 0.8302 | 2.1068 | 0.2375 | 1.1769 | 0.2392 |
| previous induced abortion | 1.0636 | 0.9886 | 1.1443 | 0.0373 | 1.6533 | 0.0982 |
| previous spontaneous abortion | 1.0419 | 0.9619 | 1.1287 | 0.0407 | 1.0082 | 0.3133 |
| education level | 1.0362 | 1.0016 | 1.0720 | 0.0173 | 2.0542 | 0.0399 |
| parental income | 0.9716 | 0.9373 | 1.0071 | 0.0182 | -1.5732 | 0.1156 |
| country | 1.5065 | 1.3342 | 1.7011 | 0.0619 | 6.6140 | <0.0001 |
| region Sjælland | 0.5995 | 0.5467 | 0.6575 | 0.0471 | -10.8578 | <0.0001 |
| region Syddanmark | 0.6471 | 0.5988 | 0.6993 | 0.0395 | -10.9932 | <0.0001 |
| region Midtjylland | 0.4922 | 0.4527 | 0.5352 | 0.0426 | -16.6034 | <0.0001 |
| region Nordjylland | 0.3523 | 0.3096 | 0.4009 | 0.0659 | -15.8221 | <0.0001 |
| mother with disorder | 1.9543 | 1.6053 | 2.3791 | 0.1003 | 6.6767 | <0.0001 |
| father with disorder | 1.4122 | 1.0653 | 1.8720 | 0.1438 | 2.4003 | 0.0163 |

Table S5
| 7. Skin - all | exp(coef) | LCL | UCL | SE | Z | Pr(Z) |
| --- | --- | --- | --- | --- | --- | --- |
| surgery | 1.3924 | 1.2939 | 1.4985 | 0.0374 | 8.8447 | <0.0001 |
| paternal age | 0.9912 | 0.9796 | 1.0029 | 0.0059 | -1.4631 | 0.1434 |
| maternal age | 0.9513 | 0.9385 | 0.9642 | 0.0069 | -7.2204 | <0.0001 |
| gestation length | 0.9959 | 0.9865 | 1.0054 | 0.0048 | -0.8331 | 0.4047 |
| maternal bleeding | 1.0664 | 1.0318 | 1.1021 | 0.0167 | 3.8285 | 0.0001 |
| fetal oxygen deprivation | 0.8474 | 0.6903 | 1.0402 | 0.1045 | -1.5822 | 0.1135 |
| pregnancy oedema | 1.1205 | 1.0428 | 1.2041 | 0.0366 | 3.1024 | 0.0019 |
| apgar5 score | 1.0923 | 1.0541 | 1.1319 | 0.0181 | 4.8674 | <0.0001 |
| birth weight | 1.0216 | 1.0125 | 1.0307 | 0.0045 | 4.6988 | <0.0001 |
| preexisting hypertension | 1.0325 | 0.8230 | 1.2953 | 0.1157 | 0.2767 | 0.7819 |
| preexisting diabetes | 1.1552 | 1.0013 | 1.3328 | 0.0729 | 1.9787 | 0.0478 |
| previous induced abortion | 1.0887 | 1.0650 | 1.1130 | 0.0112 | 7.5617 | <0.0001 |
| previous spontaneous abortion | 1.0473 | 1.0223 | 1.0729 | 0.0123 | 3.7505 | 0.0001 |
| education level | 0.9642 | 0.9544 | 0.9741 | 0.0052 | -6.9593 | <0.0001 |
| parental income | 0.9686 | 0.9580 | 0.9792 | 0.0055 | -5.7084 | <0.0001 |
| country | 1.3015 | 1.2506 | 1.3546 | 0.0203 | 12.9285 | <0.0001 |
| region Sjælland | 0.7931 | 0.7709 | 0.8159 | 0.0144 | -16.0040 | <0.0001 |
| region Syddanmark | 0.8648 | 0.8442 | 0.8860 | 0.0123 | -11.7786 | <0.0001 |
| region Midtjylland | 0.8520 | 0.8319 | 0.8727 | 0.0122 | -13.0988 | <0.0001 |
| region Nordjylland | 0.5931 | 0.5727 | 0.6143 | 0.0179 | -29.1776 | <0.0001 |
| mother with disorder | 1.3334 | 1.3002 | 1.3674 | 0.0128 | 22.3863 | <0.0001 |
| father with disorder | 1.2956 | 1.2626 | 1.3296 | 0.0131 | 19.6305 | <0.0001 |

| 8. Autoimmune - all | exp(coef) | LCL | UCL | SE | Z | Pr(Z) |
| --- | --- | --- | --- | --- | --- | --- |
| surgery | 1.1522 | 1.0218 | 1.2993 | 0.0612 | 2.3122 | 0.0207 |
| paternal age | 0.9820 | 0.9646 | 0.9997 | 0.0091 | -1.9900 | 0.0465 |
| maternal age | 0.9716 | 0.9520 | 0.9916 | 0.0104 | -2.7653 | 0.0056 |
| gestation length | 0.9930 | 0.9791 | 1.0071 | 0.0072 | -0.9706 | 0.3317 |
| maternal bleeding | 1.0831 | 1.0313 | 1.1376 | 0.0250 | 3.1916 | 0.0014 |
| fetal oxygen deprivation | 1.0618 | 0.7993 | 1.4106 | 0.1448 | 0.4144 | 0.6785 |
| pregnancy oedema | 1.2177 | 1.0967 | 1.3520 | 0.0533 | 3.6904 | 0.0002 |
| apgar5 score | 0.9849 | 0.9319 | 1.0409 | 0.0282 | -0.5370 | 0.5912 |
| birth weight | 0.9945 | 0.9814 | 1.0078 | 0.0067 | -0.8083 | 0.4188 |
| preexisting hypertension | 1.0167 | 0.7328 | 1.4106 | 0.1670 | 0.0994 | 0.9207 |
| preexisting diabetes | 1.5193 | 1.2801 | 1.8031 | 0.0873 | 4.7866 | <0.0001 |
| previous induced abortion | 0.9745 | 0.9419 | 1.0082 | 0.0173 | -1.4853 | 0.1374 |
| previous spontaneous abortion | 1.0515 | 1.0146 | 1.0898 | 0.0182 | 2.7557 | 0.0058 |
| education level | 0.9912 | 0.9760 | 1.0066 | 0.0078 | -1.1132 | 0.2655 |
| parental income | 1.0111 | 0.9946 | 1.0279 | 0.0083 | 1.3186 | 0.1872 |
| country | 0.8997 | 0.8386 | 0.9651 | 0.0358 | -2.9485 | 0.0031 |
| region Sjælland | 0.9677 | 0.9285 | 1.0085 | 0.0211 | -1.5543 | 0.1201 |
| region Syddanmark | 0.9810 | 0.9460 | 1.0173 | 0.0185 | -1.0317 | 0.3021 |
| region Midtjylland | 0.9185 | 0.8855 | 0.9527 | 0.0186 | -4.5572 | <0.0001 |
| region Nordjylland | 0.7750 | 0.7377 | 0.8142 | 0.0251 | -10.1141 | <0.0001 |
| mother with disorder | 1.5399 | 1.4781 | 1.6043 | 0.0208 | 20.6582 | <0.0001 |
| father with disorder | 1.4445 | 1.3836 | 1.5081 | 0.0219 | 16.7272 | <0.0001 |

Table S5
| 9. Respiratory - all | exp(coef) | LCL | UCL | SE | Z | Pr(Z) |
| --- | --- | --- | --- | --- | --- | --- |
| surgery | 1.4400 | 0.7488 | 2.7688 | 0.3335 | 1.0931 | 0.2743 |
| paternal age | 0.9997 | 0.9879 | 1.0117 | 0.0060 | -0.0369 | 0.9705 |
| maternal age | 0.9594 | 0.9463 | 0.9727 | 0.0070 | -5.9061 | <0.0001 |
| gestation length | 0.9853 | 0.9759 | 0.9949 | 0.0049 | -2.9920 | 0.0027 |
| maternal bleeding | 1.0835 | 1.0475 | 1.1207 | 0.0172 | 4.6531 | <0.0001 |
| fetal oxygen deprivation | 0.9174 | 0.7402 | 1.1369 | 0.1094 | -0.7875 | 0.4309 |
| pregnancy oedema | 1.1043 | 1.0234 | 1.1917 | 0.0388 | 2.5568 | 0.0105 |
| apgar5 score | 1.0526 | 1.0145 | 1.0922 | 0.0188 | 2.7254 | 0.0064 |
| birth weight | 1.0013 | 0.9923 | 1.0104 | 0.0046 | 0.2992 | 0.7647 |
| preexisting hypertension | 1.1665 | 0.9426 | 1.4436 | 0.1087 | 1.4171 | 0.1564 |
| preexisting diabetes | 1.0098 | 0.8614 | 1.1838 | 0.0811 | 0.1208 | 0.9038 |
| previous induced abortion | 1.0563 | 1.0328 | 1.0804 | 0.0114 | 4.7721 | <0.0001 |
| previous spontaneous abortion | 1.0332 | 1.0082 | 1.0587 | 0.0124 | 2.6181 | 0.0088 |
| education level | 1.0009 | 0.9906 | 1.0113 | 0.0052 | 0.1779 | 0.8587 |
| parental income | 0.9741 | 0.9633 | 0.9850 | 0.0056 | -4.6215 | <0.0001 |
| country | 1.2269 | 1.1770 | 1.2790 | 0.0211 | 9.6535 | <0.0001 |
| region Sjælland | 0.8113 | 0.7885 | 0.8348 | 0.0145 | -14.3802 | <0.0001 |
| region Syddanmark | 0.8773 | 0.8563 | 0.8989 | 0.0123 | -10.5596 | <0.0001 |
| region Midtjylland | 0.7366 | 0.7185 | 0.7552 | 0.0127 | -24.0311 | <0.0001 |
| region Nordjylland | 0.7293 | 0.7061 | 0.7533 | 0.0165 | -19.1266 | <0.0001 |
| mother with disorder | 1.3296 | 1.3012 | 1.3586 | 0.0110 | 25.8778 | <0.0001 |
| father with disorder | 1.2409 | 1.2142 | 1.2683 | 0.0111 | 19.4347 | <0.0001 |

| 10. Respiratory - upper | exp(coef) | LCL | UCL | SE | Z | Pr(Z) |
| --- | --- | --- | --- | --- | --- | --- |
| surgery | 2.7275 | 1.5479 | 4.8062 | 0.2890 | 3.4715 | 0.0005 |
| paternal age | 1.0015 | 0.9868 | 1.0165 | 0.0075 | 0.2086 | 0.8347 |
| maternal age | 0.9667 | 0.9503 | 0.9833 | 0.0086 | -3.8974 | <0.0001 |
| gestation length | 0.9926 | 0.9809 | 1.0045 | 0.0060 | -1.2117 | 0.2256 |
| maternal bleeding | 1.0770 | 1.0332 | 1.1226 | 0.0211 | 3.5040 | 0.0004 |
| fetal oxygen deprivation | 0.8364 | 0.6316 | 1.1075 | 0.1432 | -1.2466 | 0.2125 |
| pregnancy oedema | 1.1379 | 1.0373 | 1.2483 | 0.0472 | 2.7367 | 0.0062 |
| apgar5 score | 1.0880 | 1.0405 | 1.1376 | 0.0227 | 3.7046 | 0.0002 |
| birth weight | 1.0024 | 0.9913 | 1.0137 | 0.0056 | 0.4313 | 0.6662 |
| preexisting hypertension | 1.1326 | 0.8669 | 1.4797 | 0.1363 | 0.9131 | 0.3611 |
| preexisting diabetes | 0.9443 | 0.7705 | 1.1572 | 0.1037 | -0.5522 | 0.5807 |
| previous induced abortion | 1.0955 | 1.0657 | 1.1260 | 0.0140 | 6.4950 | <0.0001 |
| previous spontaneous abortion | 1.0321 | 1.0014 | 1.0637 | 0.0153 | 2.0553 | 0.0398 |
| education level | 1.0128 | 0.9999 | 1.0258 | 0.0065 | 1.9505 | 0.0511 |
| parental income | 0.9899 | 0.9764 | 1.0035 | 0.0070 | -1.4477 | 0.1476 |
| country | 1.4076 | 1.3397 | 1.4788 | 0.0252 | 13.5635 | <0.0001 |
| region Sjælland | 0.8285 | 0.7998 | 0.8582 | 0.0179 | -10.4782 | <0.0001 |
| region Syddanmark | 0.9349 | 0.9076 | 0.9631 | 0.0151 | -4.4418 | <0.0001 |
| region Midtjylland | 0.7404 | 0.7177 | 0.7638 | 0.0158 | -18.9470 | <0.0001 |
| region Nordjylland | 0.7012 | 0.6730 | 0.7306 | 0.0209 | -16.9242 | <0.0001 |
| mother with disorder | 1.3801 | 1.3365 | 1.4251 | 0.0163 | 19.6833 | <0.0001 |
| father with disorder | 1.2905 | 1.2480 | 1.3345 | 0.0170 | 14.9346 | <0.0001 |

Table S5
| 11. Respiratory - lower | exp(coef) | LCL | UCL | SE | Z | Pr(Z) |
| --- | --- | --- | --- | --- | --- | --- |
| surgery | 1.2455 | 1.0126 | 1.5321 | 0.1056 | 2.0790 | 0.0376 |
| paternal age | 1.0082 | 0.9776 | 1.0397 | 0.0157 | 0.5222 | 0.6014 |
| maternal age | 0.9128 | 0.8807 | 0.9459 | 0.0182 | -5.0081 | <0.0001 |
| gestation length | 0.9405 | 0.9170 | 0.9645 | 0.0128 | -4.7639 | <0.0001 |
| maternal bleeding | 1.1599 | 1.0662 | 1.2618 | 0.0429 | 3.4536 | 0.0005 |
| fetal oxygen deprivation | 0.8676 | 0.4919 | 1.5305 | 0.2895 | -0.4900 | 0.6240 |
| pregnancy oedema | 1.1776 | 0.9752 | 1.4220 | 0.0962 | 1.6992 | 0.0892 |
| apgar5 score | 1.1187 | 1.0193 | 1.2278 | 0.0474 | 2.3625 | 0.0181 |
| birth weight | 0.9817 | 0.9586 | 1.0053 | 0.0121 | -1.5233 | 0.1276 |
| preexisting hypertension | 1.2347 | 0.7157 | 2.1302 | 0.2782 | 0.7578 | 0.4485 |
| preexisting diabetes | 1.4446 | 1.0285 | 2.0290 | 0.1733 | 2.1225 | 0.0337 |
| previous induced abortion | 1.0027 | 0.9444 | 1.0646 | 0.0305 | 0.0904 | 0.9279 |
| previous spontaneous abortion | 1.1349 | 1.0658 | 1.2084 | 0.0320 | 3.9511 | <0.0001 |
| education level | 0.9658 | 0.9399 | 0.9924 | 0.0138 | -2.5042 | 0.0122 |
| parental income | 0.9275 | 0.9010 | 0.9549 | 0.0148 | -5.0755 | <0.0001 |
| country | 1.0940 | 0.9836 | 1.2168 | 0.0542 | 1.6558 | 0.0977 |
| region Sjælland | 0.7779 | 0.7235 | 0.8365 | 0.0370 | -6.7823 | <0.0001 |
| region Syddanmark | 0.7157 | 0.6710 | 0.7633 | 0.0328 | -10.1723 | <0.0001 |
| region Midtjylland | 0.6392 | 0.5985 | 0.6827 | 0.0335 | -13.3337 | <0.0001 |
| region Nordjylland | 0.6467 | 0.5938 | 0.7043 | 0.0435 | -10.0182 | <0.0001 |
| mother with disorder | 1.5768 | 1.4343 | 1.7334 | 0.0483 | 9.4255 | <0.0001 |
| father with disorder | 1.3687 | 1.2471 | 1.5022 | 0.0474 | 6.6124 | <0.0001 |

| 12. Respiratory - chronic lower | exp(coef) | LCL | UCL | SE | Z | Pr(Z) |
| --- | --- | --- | --- | --- | --- | --- |
| surgery | 1.3922 | 1.2469 | 1.5543 | 0.0562 | 5.8865 | <0.0001 |
| paternal age | 0.9996 | 0.9821 | 1.0174 | 0.0089 | -0.0401 | 0.9679 |
| maternal age | 0.9735 | 0.9539 | 0.9935 | 0.0103 | -2.5811 | 0.0098 |
| gestation length | 0.9689 | 0.9553 | 0.9826 | 0.0071 | -4.3906 | <0.0001 |
| maternal bleeding | 1.1222 | 1.0693 | 1.1776 | 0.0246 | 4.6867 | <0.0001 |
| fetal oxygen deprivation | 0.9788 | 0.7053 | 1.3584 | 0.1671 | -0.1276 | 0.8983 |
| pregnancy oedema | 1.1210 | 0.9985 | 1.2586 | 0.0590 | 1.9357 | 0.0528 |
| apgar5 score | 1.0757 | 1.0207 | 1.1337 | 0.0267 | 2.7253 | 0.0064 |
| birth weight | 0.9788 | 0.9657 | 0.9920 | 0.0068 | -3.1255 | 0.0017 |
| preexisting hypertension | 0.8536 | 0.5964 | 1.2217 | 0.1828 | -0.8650 | 0.3869 |
| preexisting diabetes | 0.9364 | 0.7385 | 1.1874 | 0.1211 | -0.5417 | 0.5879 |
| previous induced abortion | 1.0199 | 0.9863 | 1.0547 | 0.0171 | 1.1548 | 0.2481 |
| previous spontaneous abortion | 1.0553 | 1.0178 | 1.0941 | 0.0184 | 2.9219 | 0.0034 |
| education level | 1.0239 | 1.0080 | 1.0401 | 0.0079 | 2.9670 | 0.0030 |
| parental income | 0.9440 | 0.9283 | 0.9599 | 0.0085 | -6.7432 | <0.0001 |
| country | 0.9591 | 0.8990 | 1.0232 | 0.0330 | -1.2630 | 0.2065 |
| region Sjælland | 0.8108 | 0.7768 | 0.8463 | 0.0218 | -9.5984 | <0.0001 |
| region Syddanmark | 0.9658 | 0.9318 | 1.0010 | 0.0182 | -1.8999 | 0.0574 |
| region Midtjylland | 0.7513 | 0.7235 | 0.7803 | 0.0192 | -14.8149 | <0.0001 |
| region Nordjylland | 0.8606 | 0.8215 | 0.9017 | 0.0237 | -6.3126 | <0.0001 |
| mother with disorder | 1.9910 | 1.9002 | 2.0862 | 0.0238 | 28.9090 | <0.0001 |
| father with disorder | 1.7675 | 1.6773 | 1.8626 | 0.0267 | 21.3127 | <0.0001 |

Table S5
| 13. Respiratory - asthma | exp(coef) | LCL | UCL | SE | Z | Pr(Z) |
| --- | --- | --- | --- | --- | --- | --- |
| surgery | 1.4159 | 1.2684 | 1.5806 | 0.0561 | 6.1961 | <0.0001 |
| paternal age | 1.0091 | 0.9914 | 1.0272 | 0.0090 | 1.0115 | 0.3117 |
| maternal age | 0.9742 | 0.9545 | 0.9944 | 0.0104 | -2.4931 | 0.0126 |
| gestation length | 0.9642 | 0.9506 | 0.9779 | 0.0072 | -5.0431 | <0.0001 |
| maternal bleeding | 1.1287 | 1.0754 | 1.1846 | 0.0246 | 4.9112 | <0.0001 |
| fetal oxygen deprivation | 1.0129 | 0.7331 | 1.3994 | 0.1649 | 0.0778 | 0.9379 |
| pregnancy oedema | 1.1048 | 0.9823 | 1.2425 | 0.0599 | 1.6630 | 0.0962 |
| apgar5 score | 1.0762 | 1.0210 | 1.1345 | 0.0268 | 2.7351 | 0.0062 |
| birth weight | 0.9769 | 0.9638 | 0.9902 | 0.0068 | -3.3816 | 0.0007 |
| preexisting hypertension | 0.8542 | 0.5968 | 1.2224 | 0.1829 | -0.8615 | 0.3889 |
| preexisting diabetes | 0.9425 | 0.7433 | 1.1951 | 0.1211 | -0.4884 | 0.6252 |
| previous induced abortion | 1.0221 | 0.9883 | 1.0572 | 0.0171 | 1.2765 | 0.2017 |
| previous spontaneous abortion | 1.0589 | 1.0211 | 1.0980 | 0.0185 | 3.0920 | 0.0019 |
| education level | 1.0164 | 1.0005 | 1.0325 | 0.0080 | 2.0324 | 0.0421 |
| parental income | 0.9381 | 0.9225 | 0.9541 | 0.0085 | -7.4342 | <0.0001 |
| country | 0.9375 | 0.8784 | 1.0007 | 0.0332 | -1.9368 | 0.0527 |
| region Sjælland | 0.8117 | 0.7775 | 0.8475 | 0.0219 | -9.4899 | <0.0001 |
| region Syddanmark | 0.9631 | 0.9291 | 0.9985 | 0.0183 | -2.0394 | 0.0414 |
| region Midtjylland | 0.7556 | 0.7274 | 0.7848 | 0.0193 | -14.4720 | <0.0001 |
| region Nordjylland | 0.8586 | 0.8193 | 0.8998 | 0.0239 | -6.3746 | <0.0001 |
| mother with disorder | 2.1970 | 2.0877 | 2.3121 | 0.0260 | 30.2113 | <0.0001 |
| father with disorder | 2.0534 | 1.9294 | 2.1855 | 0.0317 | 22.6292 | <0.0001 |

| 14. Respiratory - influenza | exp(coef) | LCL | UCL | SE | Z | Pr(Z) |
| --- | --- | --- | --- | --- | --- | --- |
| surgery | 1.4366 | 1.0660 | 1.9360 | 0.1522 | 2.3802 | 0.0173 |
| paternal age | 1.0216 | 0.9735 | 1.0721 | 0.0246 | 0.8708 | 0.3838 |
| maternal age | 0.9097 | 0.8600 | 0.9622 | 0.0286 | -3.3020 | 0.0009 |
| gestation length | 0.9182 | 0.8820 | 0.9559 | 0.0205 | -4.1529 | <0.0001 |
| maternal bleeding | 1.0935 | 0.9541 | 1.2532 | 0.0695 | 1.2848 | 0.1988 |
| fetal oxygen deprivation | 1.9670 | 1.0192 | 3.7961 | 0.3354 | 2.0167 | 0.0437 |
| pregnancy oedema | 1.4519 | 1.0923 | 1.9299 | 0.1452 | 2.5680 | 0.0102 |
| apgar5 score | 1.1007 | 0.9486 | 1.2772 | 0.0758 | 1.2654 | 0.2057 |
| birth weight | 0.9723 | 0.9358 | 1.0102 | 0.0194 | -1.4389 | 0.1501 |
| preexisting hypertension | 2.8766 | 1.6261 | 5.0890 | 0.2910 | 3.6304 | 0.0002 |
| preexisting diabetes | 1.2246 | 0.6916 | 2.1681 | 0.2914 | 0.6952 | 0.4868 |
| previous induced abortion | 1.0072 | 0.9154 | 1.1081 | 0.0487 | 0.1473 | 0.8828 |
| previous spontaneous abortion | 1.0814 | 0.9765 | 1.1976 | 0.0520 | 1.5039 | 0.1326 |
| education level | 0.9496 | 0.9099 | 0.9911 | 0.0218 | -2.3665 | 0.0179 |
| parental income | 0.8955 | 0.8551 | 0.9379 | 0.0235 | -4.6772 | <0.0001 |
| country | 1.9006 | 1.6636 | 2.1714 | 0.0679 | 9.4508 | <0.0001 |
| region Sjælland | 0.7051 | 0.6293 | 0.7900 | 0.0580 | -6.0235 | <0.0001 |
| region Syddanmark | 0.5638 | 0.5074 | 0.6265 | 0.0537 | -10.6520 | <0.0001 |
| region Midtjylland | 0.5167 | 0.4640 | 0.5754 | 0.0548 | -12.0318 | <0.0001 |
| region Nordjylland | 0.4883 | 0.4213 | 0.5659 | 0.0752 | -9.5242 | <0.0001 |
| mother with disorder | 2.4573 | 1.8937 | 3.1886 | 0.1329 | 6.7638 | <0.0001 |
| father with disorder | 1.7400 | 1.2126 | 2.4969 | 0.1842 | 3.0064 | 0.0026 |

Table S5
| 15. Respiratory - pneumonia | exp(coef) | LCL | UCL | SE | Z | Pr(Z) |
| --- | --- | --- | --- | --- | --- | --- |
| surgery | 1.2847 | 1.0395 | 1.5878 | 0.1080 | 2.3184 | 0.0204 |
| paternal age | 1.0061 | 0.9741 | 1.0391 | 0.0164 | 0.3700 | 0.7113 |
| maternal age | 0.9160 | 0.8824 | 0.9509 | 0.0190 | -4.5956 | <0.0001 |
| gestation length | 0.9419 | 0.9174 | 0.9670 | 0.0134 | -4.4541 | <0.0001 |
| maternal bleeding | 1.2001 | 1.1004 | 1.3089 | 0.0442 | 4.1241 | <0.0001 |
| fetal oxygen deprivation | 0.8013 | 0.4304 | 1.4920 | 0.3171 | -0.6981 | 0.4850 |
| pregnancy oedema | 1.1802 | 0.9685 | 1.4383 | 0.1008 | 1.6428 | 0.1004 |
| apgar5 score | 1.1350 | 1.0308 | 1.2498 | 0.0491 | 2.5777 | 0.0099 |
| birth weight | 0.9923 | 0.9679 | 1.0173 | 0.0126 | -0.6071 | 0.5437 |
| preexisting hypertension | 1.3648 | 0.7910 | 2.3550 | 0.2783 | 1.1176 | 0.2637 |
| preexisting diabetes | 1.4049 | 0.9843 | 2.0052 | 0.1815 | 1.8729 | 0.0610 |
| previous induced abortion | 0.9962 | 0.9356 | 1.0607 | 0.0319 | -0.1181 | 0.9059 |
| previous spontaneous abortion | 1.1315 | 1.0595 | 1.2084 | 0.0335 | 3.6849 | 0.0002 |
| education level | 0.9619 | 0.9349 | 0.9897 | 0.0145 | -2.6652 | 0.0076 |
| parental income | 0.9382 | 0.9101 | 0.9672 | 0.0155 | -4.1069 | <0.0001 |
| country | 1.0972 | 0.9820 | 1.2260 | 0.0566 | 1.6398 | 0.1010 |
| region Sjælland | 0.7543 | 0.6990 | 0.8140 | 0.0388 | -7.2569 | <0.0001 |
| region Syddanmark | 0.7193 | 0.6727 | 0.7691 | 0.0341 | -9.6461 | <0.0001 |
| region Midtjylland | 0.6261 | 0.5844 | 0.6708 | 0.0351 | -13.3046 | <0.0001 |
| region Nordjylland | 0.6020 | 0.5495 | 0.6595 | 0.0465 | -10.8978 | <0.0001 |
| mother with disorder | 1.5816 | 1.4253 | 1.7551 | 0.0531 | 8.6340 | <0.0001 |
| father with disorder | 1.3513 | 1.2208 | 1.4957 | 0.0518 | 5.8118 | <0.0001 |

| 16. Respiratory - COPD | exp(coef) | LCL | UCL | SE | Z | Pr(Z) |
| --- | --- | --- | --- | --- | --- | --- |
| surgery | 1.2914 | 0.6683 | 2.4954 | 0.3360 | 0.7609 | 0.4467 |
| paternal age | 0.9553 | 0.8652 | 1.0547 | 0.0505 | -0.9044 | 0.3657 |
| maternal age | 0.9423 | 0.8410 | 1.0558 | 0.0580 | -1.0236 | 0.3059 |
| gestation length | 1.0288 | 0.9478 | 1.1166 | 0.0418 | 0.6792 | 0.4969 |
| maternal bleeding | 1.2019 | 0.9185 | 1.5726 | 0.1371 | 1.3408 | 0.1799 |
| fetal oxygen deprivation | 0.6765 | 0.0947 | 4.8284 | 1.0027 | -0.3897 | 0.6967 |
| pregnancy oedema | 1.6775 | 1.0459 | 2.6904 | 0.2410 | 2.1463 | 0.0318 |
| apgar5 score | 1.2120 | 0.8993 | 1.6333 | 0.1522 | 1.2633 | 0.2064 |
| birth weight | 0.8631 | 0.7991 | 0.9323 | 0.0393 | -3.7430 | 0.0001 |
| preexisting hypertension | 0.0000 | 0.0000 | Inf | 715.4281 | -0.0182 | 0.9854 |
| preexisting diabetes | 1.1341 | 0.2813 | 4.5713 | 0.7112 | 0.1769 | 0.8595 |
| previous induced abortion | 1.1440 | 0.9484 | 1.3800 | 0.0956 | 1.4070 | 0.1594 |
| previous spontaneous abortion | 1.0416 | 0.8447 | 1.2843 | 0.1068 | 0.3815 | 0.7027 |
| education level | 1.0035 | 0.9190 | 1.0959 | 0.0449 | 0.0795 | 0.9365 |
| parental income | 0.8012 | 0.7301 | 0.8792 | 0.0473 | -4.6756 | <0.0001 |
| country | 1.1277 | 0.8026 | 1.5846 | 0.1735 | 0.6928 | 0.4884 |
| region Sjælland | 0.8439 | 0.6709 | 1.0615 | 0.1170 | -1.4494 | 0.1472 |
| region Syddanmark | 0.7189 | 0.5830 | 0.8864 | 0.1068 | -3.0871 | 0.0020 |
| region Midtjylland | 0.5843 | 0.4675 | 0.7303 | 0.1137 | -4.7227 | <0.0001 |
| region Nordjylland | 0.7244 | 0.5536 | 0.9480 | 0.1372 | -2.3490 | 0.0188 |
| mother with disorder | 1.9004 | 1.3296 | 2.7163 | 0.1822 | 3.5232 | 0.0004 |
| father with disorder | 1.7672 | 1.2534 | 2.4917 | 0.1752 | 3.2487 | 0.0011 |

Table S5
| 17. Digestive - all | exp(coef) | LCL | UCL | SE | Z | Pr(Z) |
| --- | --- | --- | --- | --- | --- | --- |
| surgery | 1.1695 | 1.0861 | 1.2592 | 0.0377 | 4.1504 | <0.0001 |
| paternal age | 0.9836 | 0.9729 | 0.9944 | 0.0055 | -2.9569 | 0.0031 |
| maternal age | 0.9146 | 0.9031 | 0.9262 | 0.0064 | -13.8939 | <0.0001 |
| gestation length | 0.9811 | 0.9725 | 0.9897 | 0.0044 | -4.2553 | <0.0001 |
| maternal bleeding | 1.0617 | 1.0298 | 1.0946 | 0.0155 | 3.8483 | 0.0001 |
| fetal oxygen deprivation | 1.0670 | 0.8996 | 1.2655 | 0.0870 | 0.7457 | 0.4558 |
| pregnancy oedema | 1.1270 | 1.0567 | 1.2020 | 0.0328 | 3.6386 | 0.0002 |
| apgar5 score | 1.0604 | 1.0256 | 1.0963 | 0.0170 | 3.4448 | 0.0005 |
| birth weight | 0.9804 | 0.9724 | 0.9885 | 0.0041 | -4.7078 | <0.0001 |
| preexisting hypertension | 1.0017 | 0.8085 | 1.2411 | 0.1093 | 0.0159 | 0.9872 |
| preexisting diabetes | 1.1318 | 0.9854 | 1.2999 | 0.0706 | 1.7524 | 0.0796 |
| previous induced abortion | 1.0362 | 1.0150 | 1.0579 | 0.0105 | 3.3693 | 0.0007 |
| previous spontaneous abortion | 1.0498 | 1.0267 | 1.0734 | 0.0113 | 4.2864 | <0.0001 |
| education level | 0.9593 | 0.9502 | 0.9684 | 0.0048 | -8.5621 | <0.0001 |
| parental income | 0.9852 | 0.9752 | 0.9952 | 0.0051 | -2.8741 | 0.0040 |
| country | 0.8471 | 0.8112 | 0.8847 | 0.0221 | -7.4960 | <0.0001 |
| region Sjælland | 0.9638 | 0.9391 | 0.9891 | 0.0132 | -2.7818 | 0.0054 |
| region Syddanmark | 1.0317 | 1.0087 | 1.0551 | 0.0114 | 2.7237 | 0.0064 |
| region Midtjylland | 0.8742 | 0.8542 | 0.8947 | 0.0118 | -11.3621 | <0.0001 |
| region Nordjylland | 0.9170 | 0.8909 | 0.9438 | 0.0147 | -5.8822 | <0.0001 |
| mother with disorder | 1.3003 | 1.2775 | 1.3235 | 0.0090 | 29.0573 | <0.0001 |
| father with disorder | 1.1832 | 1.1625 | 1.2044 | 0.0090 | 18.6474 | <0.0001 |

| 18. Endocrine - all | exp(coef) | LCL | UCL | SE | Z | Pr(Z) |
| --- | --- | --- | --- | --- | --- | --- |
| surgery | 1.3109 | 1.1775 | 1.4594 | 0.0547 | 4.9445 | <0.0001 |
| paternal age | 0.9506 | 0.9354 | 0.9660 | 0.0082 | -6.1565 | <0.0001 |
| maternal age | 0.8678 | 0.8518 | 0.8842 | 0.0095 | -14.8606 | <0.0001 |
| gestation length | 0.9858 | 0.9728 | 0.9990 | 0.0067 | -2.0925 | 0.0363 |
| maternal bleeding | 1.0583 | 1.0101 | 1.1089 | 0.0238 | 2.3841 | 0.0171 |
| fetal oxygen deprivation | 1.1836 | 0.9271 | 1.5110 | 0.1246 | 1.3528 | 0.1760 |
| pregnancy oedema | 1.2449 | 1.1438 | 1.3549 | 0.0432 | 5.0713 | <0.0001 |
| apgar5 score | 1.1617 | 1.1055 | 1.2206 | 0.0252 | 5.9314 | <0.0001 |
| birth weight | 1.0355 | 1.0227 | 1.0484 | 0.0063 | 5.5061 | <0.0001 |
| preexisting hypertension | 1.0762 | 0.7946 | 1.4576 | 0.1547 | 0.4747 | 0.6349 |
| preexisting diabetes | 1.5062 | 1.2786 | 1.7743 | 0.0835 | 4.9005 | <0.0001 |
| previous induced abortion | 1.0243 | 0.9923 | 1.0573 | 0.0161 | 1.4831 | 0.1380 |
| previous spontaneous abortion | 1.0555 | 1.0203 | 1.0919 | 0.0173 | 3.1207 | 0.0018 |
| education level | 0.8463 | 0.8344 | 0.8585 | 0.0072 | -22.9296 | <0.0001 |
| parental income | 0.8855 | 0.8720 | 0.8992 | 0.0078 | -15.5256 | <0.0001 |
| country | 0.8490 | 0.7998 | 0.9011 | 0.0304 | -5.3800 | <0.0001 |
| region Sjælland | 1.2465 | 1.2004 | 1.2943 | 0.0192 | 11.4728 | <0.0001 |
| region Syddanmark | 1.1640 | 1.1250 | 1.2043 | 0.0173 | 8.7420 | <0.0001 |
| region Midtjylland | 0.8216 | 0.7918 | 0.8525 | 0.0188 | -10.4284 | <0.0001 |
| region Nordjylland | 0.8624 | 0.8237 | 0.9030 | 0.0234 | -6.3035 | <0.0001 |
| mother with disorder | 1.6553 | 1.6079 | 1.7042 | 0.0148 | 33.9806 | <0.0001 |
| father with disorder | 1.5066 | 1.4560 | 1.5590 | 0.0174 | 23.5112 | <0.0001 |

Table S5
| 19. Endocrine - obesity | exp(coef) | LCL | UCL | SE | Z | Pr(Z) |
| --- | --- | --- | --- | --- | --- | --- |
| surgery | 1.4035 | 1.2319 | 1.5991 | 0.0665 | 5.0951 | <0.0001 |
| paternal age | 0.9270 | 0.9086 | 0.9458 | 0.0102 | -7.4072 | <0.0001 |
| maternal age | 0.8194 | 0.8004 | 0.8390 | 0.0120 | -16.5699 | <0.0001 |
| gestation length | 1.0024 | 0.9858 | 1.0194 | 0.0085 | 0.2923 | 0.7700 |
| maternal bleeding | 1.0644 | 1.0037 | 1.1288 | 0.0299 | 2.0841 | 0.0371 |
| fetal oxygen deprivation | 1.2060 | 0.8899 | 1.6342 | 0.1550 | 1.2082 | 0.2269 |
| pregnancy oedema | 1.2866 | 1.1650 | 1.4208 | 0.0506 | 4.9785 | <0.0001 |
| apgar5 score | 1.1963 | 1.1250 | 1.2722 | 0.0313 | 5.7189 | <0.0001 |
| birth weight | 1.0746 | 1.0579 | 1.0915 | 0.0079 | 9.0279 | <0.0001 |
| preexisting hypertension | 1.1458 | 0.7950 | 1.6514 | 0.1864 | 0.7301 | 0.4653 |
| preexisting diabetes | 1.6444 | 1.3380 | 2.0208 | 0.1051 | 4.7290 | <0.0001 |
| previous induced abortion | 1.0238 | 0.9836 | 1.0657 | 0.0204 | 1.1539 | 0.2485 |
| previous spontaneous abortion | 1.0756 | 1.0303 | 1.1229 | 0.0219 | 3.3230 | 0.0008 |
| education level | 0.7634 | 0.7497 | 0.7774 | 0.0092 | -29.1082 | <0.0001 |
| parental income | 0.8561 | 0.8396 | 0.8728 | 0.0099 | -15.6876 | <0.0001 |
| country | 0.6092 | 0.5615 | 0.6610 | 0.0416 | -11.9058 | <0.0001 |
| region Sjælland | 1.4771 | 1.4096 | 1.5479 | 0.0238 | 16.3335 | <0.0001 |
| region Syddanmark | 1.2958 | 1.2408 | 1.3533 | 0.0221 | 11.7073 | <0.0001 |
| region Midtjylland | 0.7949 | 0.7571 | 0.8346 | 0.0248 | -9.2347 | <0.0001 |
| region Nordjylland | 0.8510 | 0.8016 | 0.9035 | 0.0305 | -5.2858 | <0.0001 |
| mother with disorder | 2.3414 | 2.2320 | 2.4561 | 0.0244 | 34.8526 | <0.0001 |
| father with disorder | 1.9915 | 1.8490 | 2.1450 | 0.0378 | 18.1910 | <0.0001 |

| 20. Genitourinary - all | exp(coef) | LCL | UCL | SE | Z | Pr(Z) |
| --- | --- | --- | --- | --- | --- | --- |
| surgery | 1.2709 | 1.1337 | 1.4247 | 0.0582 | 4.1132 | <0.0001 |
| paternal age | 0.9697 | 0.9533 | 0.9863 | 0.0086 | -3.5479 | 0.0003 |
| maternal age | 0.8892 | 0.8718 | 0.9070 | 0.0101 | -11.6181 | <0.0001 |
| gestation length | 0.9799 | 0.9663 | 0.9936 | 0.0071 | -2.8492 | 0.0043 |
| maternal bleeding | 1.1145 | 1.0631 | 1.1685 | 0.0241 | 4.4969 | <0.0001 |
| fetal oxygen deprivation | 0.8451 | 0.6143 | 1.1625 | 0.1627 | -1.0340 | 0.3010 |
| pregnancy oedema | 1.1302 | 1.0190 | 1.2536 | 0.0528 | 2.3175 | 0.0204 |
| apgar5 score | 1.0438 | 0.9888 | 1.1018 | 0.0275 | 1.5549 | 0.1199 |
| birth weight | 0.9574 | 0.9449 | 0.9701 | 0.0067 | -6.4757 | <0.0001 |
| preexisting hypertension | 1.1551 | 0.8466 | 1.5760 | 0.1585 | 0.9100 | 0.3627 |
| preexisting diabetes | 1.2174 | 0.9851 | 1.5046 | 0.1080 | 1.8214 | 0.0685 |
| previous induced abortion | 1.1254 | 1.0902 | 1.1618 | 0.0162 | 7.2793 | <0.0001 |
| previous spontaneous abortion | 1.0422 | 1.0058 | 1.0799 | 0.0181 | 2.2808 | 0.0225 |
| education level | 0.9231 | 0.9093 | 0.9370 | 0.0076 | -10.4461 | <0.0001 |
| parental income | 0.9278 | 0.9131 | 0.9426 | 0.0081 | -9.2417 | <0.0001 |
| country | 0.8833 | 0.8307 | 0.9392 | 0.0313 | -3.9592 | <0.0001 |
| region Sjælland | 0.7105 | 0.6824 | 0.7397 | 0.0205 | -16.6272 | <0.0001 |
| region Syddanmark | 0.6690 | 0.6456 | 0.6931 | 0.0181 | -22.1925 | <0.0001 |
| region Midtjylland | 0.6222 | 0.6002 | 0.6451 | 0.0183 | -25.8170 | <0.0001 |
| region Nordjylland | 0.6236 | 0.5952 | 0.6535 | 0.0238 | -19.8144 | <0.0001 |
| mother with disorder | 1.3189 | 1.2766 | 1.3625 | 0.0166 | 16.6573 | <0.0001 |
| father with disorder | 1.2736 | 1.2166 | 1.3332 | 0.0233 | 10.3625 | <0.0001 |

Table S5
| 21. Genitourinary - kidney infection | exp(coef) | LCL | UCL | SE | Z | Pr(Z) |
| --- | --- | --- | --- | --- | --- | --- |
| surgery | 1.2889 | 0.9632 | 1.7245 | 0.1485 | 1.7081 | 0.0876 |
| paternal age | 0.9999 | 0.9577 | 1.0440 | 0.0220 | -0.0006 | 0.9994 |
| maternal age | 0.8507 | 0.8087 | 0.8949 | 0.0258 | -6.2576 | <0.0001 |
| gestation length | 1.0171 | 0.9818 | 1.0536 | 0.0180 | 0.9419 | 0.3462 |
| maternal bleeding | 1.1192 | 0.9922 | 1.2624 | 0.0614 | 1.8329 | 0.0668 |
| fetal oxygen deprivation | 0.9303 | 0.4169 | 2.0760 | 0.4095 | -0.1762 | 0.8601 |
| pregnancy oedema | 0.8642 | 0.6320 | 1.1817 | 0.1596 | -0.9138 | 0.3608 |
| apgar5 score | 1.1312 | 0.9897 | 1.2930 | 0.0681 | 1.8086 | 0.0705 |
| birth weight | 0.9892 | 0.9568 | 1.0227 | 0.0169 | -0.6345 | 0.5257 |
| preexisting hypertension | 0.5301 | 0.1707 | 1.6463 | 0.5780 | -1.0975 | 0.2723 |
| preexisting diabetes | 1.4537 | 0.8995 | 2.3495 | 0.2449 | 1.5275 | 0.1266 |
| previous induced abortion | 1.1812 | 1.0914 | 1.2784 | 0.0403 | 4.1275 | <0.0001 |
| previous spontaneous abortion | 1.1176 | 1.0237 | 1.2200 | 0.0447 | 2.4839 | 0.0129 |
| education level | 0.9277 | 0.8927 | 0.9640 | 0.0196 | -3.8268 | 0.0001 |
| parental income | 0.9574 | 0.9193 | 0.9970 | 0.0207 | -2.1015 | 0.0355 |
| country | 0.6379 | 0.5350 | 0.7606 | 0.0897 | -5.0086 | <0.0001 |
| region Sjælland | 0.6935 | 0.6261 | 0.7682 | 0.0521 | -7.0134 | <0.0001 |
| region Syddanmark | 0.6344 | 0.5792 | 0.6948 | 0.0464 | -9.7977 | <0.0001 |
| region Midtjylland | 0.6425 | 0.5872 | 0.7029 | 0.0458 | -9.6443 | <0.0001 |
| region Nordjylland | 0.6320 | 0.5623 | 0.7103 | 0.0595 | -7.7004 | <0.0001 |
| mother with disorder | 1.4632 | 1.1728 | 1.8256 | 0.1128 | 3.3724 | 0.0007 |
| father with disorder | 1.6448 | 1.0705 | 2.5272 | 0.2191 | 2.2711 | 0.0231 |

| 22. Musculoskeletal - all | exp(coef) | LCL | UCL | SE | Z | Pr(Z) |
| --- | --- | --- | --- | --- | --- | --- |
| surgery | 1.2185 | 1.1547 | 1.2857 | 0.0274 | 7.2069 | <0.0001 |
| paternal age | 0.9737 | 0.9658 | 0.9818 | 0.0041 | -6.3389 | <0.0001 |
| maternal age | 0.9359 | 0.9271 | 0.9448 | 0.0048 | -13.7206 | <0.0001 |
| gestation length | 1.0001 | 0.9936 | 1.0066 | 0.0033 | 0.0409 | 0.9673 |
| maternal bleeding | 1.0874 | 1.0631 | 1.1123 | 0.0115 | 7.2734 | <0.0001 |
| fetal oxygen deprivation | 1.1597 | 1.0294 | 1.3065 | 0.0608 | 2.4375 | 0.0147 |
| pregnancy oedema | 1.0978 | 1.0429 | 1.1556 | 0.0261 | 3.5663 | 0.0003 |
| apgar5 score | 1.0158 | 0.9904 | 1.0420 | 0.0129 | 1.2170 | 0.2235 |
| birth weight | 0.9904 | 0.9843 | 0.9965 | 0.0031 | -3.0694 | 0.0021 |
| preexisting hypertension | 1.0329 | 0.8856 | 1.2047 | 0.0784 | 0.4131 | 0.6795 |
| preexisting diabetes | 1.0178 | 0.9154 | 1.1317 | 0.0541 | 0.3275 | 0.7432 |
| previous induced abortion | 1.0467 | 1.0307 | 1.0629 | 0.0078 | 5.8128 | <0.0001 |
| previous spontaneous abortion | 1.0317 | 1.0146 | 1.0491 | 0.0085 | 3.6718 | 0.0002 |
| education level | 0.9555 | 0.9487 | 0.9624 | 0.0036 | -12.4498 | <0.0001 |
| parental income | 0.9929 | 0.9853 | 1.0005 | 0.0039 | -1.8176 | 0.0691 |
| country | 0.8543 | 0.8275 | 0.8819 | 0.0162 | -9.7021 | <0.0001 |
| region Sjælland | 0.8484 | 0.8315 | 0.8657 | 0.0102 | -15.9992 | <0.0001 |
| region Syddanmark | 0.8867 | 0.8716 | 0.9022 | 0.0088 | -13.6500 | <0.0001 |
| region Midtjylland | 1.0416 | 1.0246 | 1.0589 | 0.0084 | 4.8541 | <0.0001 |
| region Nordjylland | 0.8347 | 0.8163 | 0.8536 | 0.0113 | -15.8763 | <0.0001 |
| mother with disorder | 1.3431 | 1.3268 | 1.3597 | 0.0062 | 47.3507 | <0.0001 |
| father with disorder | 1.2313 | 1.2162 | 1.2466 | 0.0062 | 33.0863 | <0.0001 |

Table S5
| 23. Neoplasms - all | exp(coef) | LCL | UCL | SE | Z | Pr(Z) |
| --- | --- | --- | --- | --- | --- | --- |
| surgery | 1.2211 | 1.0966 | 1.3598 | 0.0548 | 3.6415 | 0.0002 |
| paternal age | 0.9855 | 0.9697 | 1.0016 | 0.0082 | -1.7577 | 0.0787 |
| maternal age | 0.9694 | 0.9516 | 0.9874 | 0.0094 | -3.2962 | 0.0009 |
| gestation length | 0.9982 | 0.9855 | 1.0110 | 0.0065 | -0.2716 | 0.7859 |
| maternal bleeding | 1.0751 | 1.0282 | 1.1240 | 0.0227 | 3.1877 | 0.0014 |
| fetal oxygen deprivation | 1.1286 | 0.8935 | 1.4256 | 0.1191 | 1.0154 | 0.3098 |
| pregnancy oedema | 0.9849 | 0.8908 | 1.0890 | 0.0512 | -0.2953 | 0.7677 |
| apgar5 score | 0.9769 | 0.9287 | 1.0277 | 0.0258 | -0.9003 | 0.3679 |
| birth weight | 1.0171 | 1.0049 | 1.0293 | 0.0061 | 2.7748 | 0.0055 |
| preexisting hypertension | 0.9508 | 0.6914 | 1.3074 | 0.1625 | -0.3103 | 0.7562 |
| preexisting diabetes | 1.0287 | 0.8326 | 1.2709 | 0.1078 | 0.2623 | 0.7930 |
| previous induced abortion | 0.9953 | 0.9652 | 1.0262 | 0.0156 | -0.3011 | 0.7632 |
| previous spontaneous abortion | 1.0262 | 0.9934 | 1.0601 | 0.0165 | 1.5623 | 0.1182 |
| education level | 1.0071 | 0.9932 | 1.0212 | 0.0070 | 1.0080 | 0.3134 |
| parental income | 1.0281 | 1.0130 | 1.0434 | 0.0075 | 3.6791 | 0.0002 |
| country | 0.9152 | 0.8554 | 0.9792 | 0.0344 | -2.5675 | 0.0102 |
| region Sjælland | 1.1109 | 1.0693 | 1.1542 | 0.0194 | 5.4004 | <0.0001 |
| region Syddanmark | 1.1857 | 1.1468 | 1.2259 | 0.0170 | 10.0164 | <0.0001 |
| region Midtjylland | 1.1391 | 1.1018 | 1.1777 | 0.0169 | 7.6675 | <0.0001 |
| region Nordjylland | 0.9369 | 0.8963 | 0.9793 | 0.0225 | -2.8842 | 0.0039 |
| mother with disorder | 1.1890 | 1.1599 | 1.2188 | 0.0126 | 13.7133 | <0.0001 |
| father with disorder | 1.1534 | 1.1158 | 1.1923 | 0.0169 | 8.4357 | <0.0001 |

| 24. Neoplasms - benign | exp(coef) | LCL | UCL | SE | Z | Pr(Z) |
| --- | --- | --- | --- | --- | --- | --- |
| surgery | 1.2251 | 1.0916 | 1.3749 | 0.0588 | 3.4504 | 0.0005 |
| paternal age | 0.9803 | 0.9633 | 0.9975 | 0.0089 | -2.2307 | 0.0257 |
| maternal age | 0.9763 | 0.9571 | 0.9960 | 0.0101 | -2.3535 | 0.0185 |
| gestation length | 0.9963 | 0.9827 | 1.0101 | 0.0070 | -0.5236 | 0.6004 |
| maternal bleeding | 1.0647 | 1.0148 | 1.1171 | 0.0245 | 2.5610 | 0.0104 |
| fetal oxygen deprivation | 1.1699 | 0.9111 | 1.5022 | 0.1275 | 1.2306 | 0.2184 |
| pregnancy oedema | 0.9981 | 0.8950 | 1.1131 | 0.0556 | -0.0331 | 0.9735 |
| apgar5 score | 0.9698 | 0.9183 | 1.0242 | 0.0278 | -1.0993 | 0.2715 |
| birth weight | 1.0140 | 1.0010 | 1.0272 | 0.0065 | 2.1216 | 0.0338 |
| preexisting hypertension | 0.8833 | 0.6208 | 1.2567 | 0.1799 | -0.6896 | 0.4903 |
| preexisting diabetes | 1.0415 | 0.8318 | 1.3040 | 0.1146 | 0.3548 | 0.7226 |
| previous induced abortion | 1.0013 | 0.9690 | 1.0348 | 0.0167 | 0.0814 | 0.9350 |
| previous spontaneous abortion | 1.0474 | 1.0117 | 1.0845 | 0.0177 | 2.6187 | 0.0088 |
| education level | 1.0126 | 0.9975 | 1.0279 | 0.0076 | 1.6439 | 0.1001 |
| parental income | 1.0253 | 1.0091 | 1.0418 | 0.0081 | 3.0778 | 0.0020 |
| country | 0.9220 | 0.8580 | 0.9907 | 0.0366 | -2.2124 | 0.0269 |
| region Sjælland | 1.0985 | 1.0541 | 1.1448 | 0.0210 | 4.4670 | <0.0001 |
| region Syddanmark | 1.1979 | 1.1557 | 1.2416 | 0.0182 | 9.8824 | <0.0001 |
| region Midtjylland | 1.1487 | 1.1083 | 1.1905 | 0.0182 | 7.5940 | <0.0001 |
| region Nordjylland | 0.8990 | 0.8566 | 0.9436 | 0.0246 | -4.3137 | <0.0001 |
| mother with disorder | 1.2202 | 1.1852 | 1.2562 | 0.0148 | 13.4010 | <0.0001 |
| father with disorder | 1.1921 | 1.1412 | 1.2452 | 0.0222 | 7.9035 | <0.0001 |

Table S5
| 25. Circulatory - all | exp(coef) | LCL | UCL | SE | Z | Pr(Z) |
| --- | --- | --- | --- | --- | --- | --- |
| surgery | 1.1715 | 0.9997 | 1.3729 | 0.0809 | 1.9565 | 0.0503 |
| paternal age | 0.9688 | 0.9459 | 0.9922 | 0.0121 | -2.5997 | 0.0093 |
| maternal age | 0.9481 | 0.9227 | 0.9743 | 0.0138 | -3.8377 | 0.0001 |
| gestation length | 0.9712 | 0.9530 | 0.9898 | 0.0096 | -3.0116 | 0.0025 |
| maternal bleeding | 1.1373 | 1.0670 | 1.2122 | 0.0325 | 3.9549 | <0.0001 |
| fetal oxygen deprivation | 0.8505 | 0.5782 | 1.2508 | 0.1968 | -0.8226 | 0.4107 |
| pregnancy oedema | 1.0466 | 0.9091 | 1.2048 | 0.0718 | 0.6341 | 0.5259 |
| apgar5 score | 1.2291 | 1.1481 | 1.3158 | 0.0347 | 5.9292 | <0.0001 |
| birth weight | 0.9757 | 0.9585 | 0.9932 | 0.0090 | -2.7033 | 0.0068 |
| preexisting hypertension | 1.1880 | 0.8233 | 1.7141 | 0.1870 | 0.9209 | 0.3570 |
| preexisting diabetes | 0.9621 | 0.7068 | 1.3096 | 0.1573 | -0.2454 | 0.8061 |
| previous induced abortion | 0.9862 | 0.9422 | 1.0323 | 0.0232 | -0.5927 | 0.5533 |
| previous spontaneous abortion | 1.0432 | 0.9940 | 1.0949 | 0.0246 | 1.7173 | 0.0859 |
| education level | 0.9706 | 0.9508 | 0.9908 | 0.0105 | -2.8323 | 0.0046 |
| parental income | 0.9825 | 0.9613 | 1.0042 | 0.0111 | -1.5761 | 0.1149 |
| country | 0.8854 | 0.8052 | 0.9735 | 0.0484 | -2.5134 | 0.0119 |
| region Sjælland | 0.9937 | 0.9405 | 1.0500 | 0.0280 | -0.2215 | 0.8246 |
| region Syddanmark | 0.9101 | 0.8661 | 0.9564 | 0.0253 | -3.7177 | 0.0002 |
| region Midtjylland | 0.9468 | 0.9018 | 0.9941 | 0.0248 | -2.1940 | 0.0282 |
| region Nordjylland | 0.7628 | 0.7138 | 0.8151 | 0.0338 | -8.0004 | <0.0001 |
| mother with disorder | 1.4985 | 1.4338 | 1.5661 | 0.0225 | 17.9650 | <0.0001 |
| father with disorder | 1.3353 | 1.2811 | 1.3917 | 0.0211 | 13.6985 | <0.0001 |

| 26. Nervous System - all | exp(coef) | LCL | UCL | SE | Z | Pr(Z) |
| --- | --- | --- | --- | --- | --- | --- |
| surgery | 1.5085 | 1.2620 | 1.8031 | 0.0910 | 4.5166 | <0.0001 |
| paternal age | 0.9776 | 0.9490 | 1.0070 | 0.0151 | -1.4953 | 0.1348 |
| maternal age | 0.8889 | 0.8588 | 0.9199 | 0.0175 | -6.7135 | <0.0001 |
| gestation length | 0.9955 | 0.9718 | 1.0198 | 0.0122 | -0.3622 | 0.7171 |
| maternal bleeding | 1.1232 | 1.0352 | 1.2187 | 0.0416 | 2.7909 | 0.0052 |
| fetal oxygen deprivation | 1.5467 | 1.0724 | 2.2309 | 0.1868 | 2.3341 | 0.0195 |
| pregnancy oedema | 1.2077 | 1.0301 | 1.4159 | 0.0811 | 2.3256 | 0.0200 |
| apgar5 score | 1.0712 | 0.9767 | 1.1750 | 0.0471 | 1.4605 | 0.1441 |
| birth weight | 0.9770 | 0.9553 | 0.9992 | 0.0114 | -2.0268 | 0.0426 |
| preexisting hypertension | 1.1102 | 0.6296 | 1.9575 | 0.2893 | 0.3614 | 0.7177 |
| preexisting diabetes | 1.4977 | 1.0718 | 2.0927 | 0.1706 | 2.3665 | 0.0179 |
| previous induced abortion | 1.0604 | 1.0018 | 1.1224 | 0.0289 | 2.0252 | 0.0428 |
| previous spontaneous abortion | 1.0826 | 1.0191 | 1.1501 | 0.0308 | 2.5730 | 0.0100 |
| education level | 0.9062 | 0.8830 | 0.9301 | 0.0132 | -7.4335 | <0.0001 |
| parental income | 0.9368 | 0.9112 | 0.9631 | 0.0141 | -4.6145 | <0.0001 |
| country | 0.7051 | 0.6197 | 0.8023 | 0.0658 | -5.3038 | <0.0001 |
| region Sjælland | 1.4341 | 1.3411 | 1.5336 | 0.0342 | 10.5360 | <0.0001 |
| region Syddanmark | 1.1590 | 1.0880 | 1.2345 | 0.0322 | 4.5797 | <0.0001 |
| region Midtjylland | 0.9868 | 0.9245 | 1.0534 | 0.0333 | -0.3968 | 0.6914 |
| region Nordjylland | 0.7523 | 0.6877 | 0.8229 | 0.0457 | -6.2153 | <0.0001 |
| mother with disorder | 1.5647 | 1.4591 | 1.6780 | 0.0356 | 12.5576 | <0.0001 |
| father with disorder | 1.4133 | 1.2973 | 1.5397 | 0.0437 | 7.9168 | <0.0001 |

Table S5
| 27. Eye/adnexa - all | exp(coef) | LCL | UCL | SE | Z | Pr(Z) |
| --- | --- | --- | --- | --- | --- | --- |
| surgery | 1.2531 | 1.1274 | 1.3928 | 0.0539 | 4.1857 | <0.0001 |
| paternal age | 1.0033 | 0.9880 | 1.0189 | 0.0078 | 0.4281 | 0.6685 |
| maternal age | 0.9744 | 0.9574 | 0.9917 | 0.0089 | -2.8761 | 0.0040 |
| gestation length | 0.9570 | 0.9452 | 0.9690 | 0.0063 | -6.9171 | <0.0001 |
| maternal bleeding | 1.0874 | 1.0419 | 1.1348 | 0.0218 | 3.8421 | 0.0001 |
| fetal oxygen deprivation | 1.1464 | 0.8947 | 1.4690 | 0.1265 | 1.0804 | 0.2799 |
| pregnancy oedema | 1.0639 | 0.9643 | 1.1737 | 0.0501 | 1.2363 | 0.2163 |
| apgar5 score | 1.0673 | 1.0184 | 1.1185 | 0.0239 | 2.7253 | 0.0064 |
| birth weight | 0.9581 | 0.9470 | 0.9694 | 0.0059 | -7.1573 | <0.0001 |
| preexisting hypertension | 1.0024 | 0.7455 | 1.3478 | 0.1510 | 0.0159 | 0.9872 |
| preexisting diabetes | 1.0952 | 0.9007 | 1.3318 | 0.0997 | 0.9122 | 0.3616 |
| previous induced abortion | 1.0290 | 0.9993 | 1.0596 | 0.0149 | 1.9192 | 0.0549 |
| previous spontaneous abortion | 1.0694 | 1.0364 | 1.1034 | 0.0159 | 4.2025 | <0.0001 |
| education level | 0.9591 | 0.9463 | 0.9720 | 0.0068 | -6.1012 | <0.0001 |
| parental income | 0.9569 | 0.9433 | 0.9706 | 0.0072 | -6.0417 | <0.0001 |
| country | 1.0354 | 0.9806 | 1.0934 | 0.0277 | 1.2548 | 0.2095 |
| region Sjælland | 0.7687 | 0.7415 | 0.7969 | 0.0183 | -14.3111 | <0.0001 |
| region Syddanmark | 0.7515 | 0.7283 | 0.7755 | 0.0160 | -17.8031 | <0.0001 |
| region Midtjylland | 0.6381 | 0.6176 | 0.6592 | 0.0166 | -26.9935 | <0.0001 |
| region Nordjylland | 0.6704 | 0.6429 | 0.6991 | 0.0213 | -18.6939 | <0.0001 |
| mother with disorder | 1.3262 | 1.2781 | 1.3761 | 0.0188 | 14.9739 | <0.0001 |
| father with disorder | 1.2401 | 1.1941 | 1.2880 | 0.0193 | 11.1484 | <0.0001 |

| 28. Mental - all | exp(coef) | LCL | UCL | SE | Z | Pr(Z) |
| --- | --- | --- | --- | --- | --- | --- |
| surgery | 1.2229 | 1.1295 | 1.3239 | 0.0405 | 4.9665 | <0.0001 |
| paternal age | 1.0143 | 1.0026 | 1.0261 | 0.0059 | 2.4050 | 0.0161 |
| maternal age | 0.9173 | 0.9050 | 0.9299 | 0.0069 | -12.4658 | <0.0001 |
| gestation length | 0.9821 | 0.9726 | 0.9917 | 0.0049 | -3.6345 | 0.0002 |
| maternal bleeding | 1.1313 | 1.0953 | 1.1684 | 0.0164 | 7.4817 | <0.0001 |
| fetal oxygen deprivation | 1.2019 | 0.9971 | 1.4489 | 0.0953 | 1.9300 | 0.0535 |
| pregnancy oedema | 1.0852 | 1.0062 | 1.1705 | 0.0385 | 2.1203 | 0.0339 |
| apgar5 score | 1.0591 | 1.0206 | 1.0990 | 0.0188 | 3.0447 | 0.0023 |
| birth weight | 0.9710 | 0.9621 | 0.9799 | 0.0046 | -6.2740 | <0.0001 |
| preexisting hypertension | 1.1793 | 0.9504 | 1.4633 | 0.1101 | 1.4980 | 0.1341 |
| preexisting diabetes | 1.1352 | 0.9809 | 1.3138 | 0.0745 | 1.7015 | 0.0888 |
| previous induced abortion | 1.1292 | 1.1047 | 1.1543 | 0.0112 | 10.8523 | <0.0001 |
| previous spontaneous abortion | 1.0692 | 1.0432 | 1.0958 | 0.0125 | 5.3334 | <0.0001 |
| education level | 0.9600 | 0.9497 | 0.9703 | 0.0054 | -7.4523 | <0.0001 |
| parental income | 0.7864 | 0.7774 | 0.7955 | 0.0058 | -40.8631 | <0.0001 |
| country | 0.4841 | 0.4590 | 0.5105 | 0.0271 | -26.7619 | <0.0001 |
| region Sjælland | 0.8414 | 0.8177 | 0.8657 | 0.0145 | -11.8655 | <0.0001 |
| region Syddanmark | 0.9152 | 0.8933 | 0.9377 | 0.0123 | -7.1517 | <0.0001 |
| region Midtjylland | 0.7653 | 0.7460 | 0.7851 | 0.0130 | -20.5091 | <0.0001 |
| region Nordjylland | 0.5591 | 0.5389 | 0.5801 | 0.0187 | -30.9937 | <0.0001 |
| mother with disorder | 2.0420 | 1.9934 | 2.0917 | 0.0122 | 58.0955 | <0.0001 |
| father with disorder | 1.6294 | 1.5836 | 1.6765 | 0.0145 | 33.5810 | <0.0001 |

**Table S6.** **Full Cox regression outputs - impact of adenoidectomy on risk of later disease.**

| 1. Infectious - all |  |  |  |  |  |  |
| --- | --- | --- | --- | --- | --- | --- |
|  | exp(coef) | LCL | UCL | SE | Z | Pr(Z) |
| surgery | 1.2751 | 1.1931 | 1.3627 | 0.0339 | 7.1651 | <0.0001 |
| paternal age | 0.9933 | 0.9796 | 1.0072 | 0.0070 | -0.9405 | 0.3469 |
| maternal age | 0.9731 | 0.9578 | 0.9887 | 0.0081 | -3.3560 | 0.0007 |
| gestation length | 0.9878 | 0.9768 | 0.9989 | 0.0057 | -2.1485 | 0.0316 |
| maternal bleeding | 1.1079 | 1.0661 | 1.1513 | 0.0196 | 5.2296 | <0.0001 |
| fetal oxygen deprivation | 0.8918 | 0.6948 | 1.1448 | 0.1273 | -0.8980 | 0.3691 |
| pregnancy oedema | 1.0579 | 0.9673 | 1.1570 | 0.0456 | 1.2333 | 0.2174 |
| apgar5 score | 1.0433 | 0.9994 | 1.0892 | 0.0219 | 1.9343 | 0.0530 |
| birth weight | 0.9913 | 0.9810 | 1.0018 | 0.0053 | -1.6162 | 0.1060 |
| preexisting hypertension | 1.0357 | 0.7966 | 1.3465 | 0.1338 | 0.2625 | 0.7929 |
| preexisting diabetes | 1.0546 | 0.8814 | 1.2618 | 0.0915 | 0.5809 | 0.5612 |
| previous induced abortion | 1.0931 | 1.0654 | 1.1216 | 0.0131 | 6.7951 | <0.0001 |
| previous spontaneous abortion | 1.0461 | 1.0171 | 1.0760 | 0.0143 | 3.1438 | 0.0016 |
| education level | 0.9923 | 0.9805 | 1.0043 | 0.0061 | -1.2552 | 0.2093 |
| parental income | 0.9864 | 0.9739 | 0.9991 | 0.0065 | -2.0943 | 0.0362 |
| country | 1.2691 | 1.2104 | 1.3306 | 0.0241 | 9.8712 | <0.0001 |
| region Sjælland | 0.7845 | 0.7599 | 0.8099 | 0.0162 | -14.9030 | <0.0001 |
| region Syddanmark | 0.6846 | 0.6651 | 0.7046 | 0.0147 | -25.7642 | <0.0001 |
| region Midtjylland | 0.7020 | 0.6824 | 0.7223 | 0.0144 | -24.4181 | <0.0001 |
| region Nordjylland | 0.6156 | 0.5923 | 0.6399 | 0.0196 | -24.6213 | <0.0001 |
| mother with disorder | 1.3780 | 1.3346 | 1.4229 | 0.0163 | 19.6287 | <0.0001 |
| father with disorder | 1.2447 | 1.2036 | 1.2873 | 0.0171 | 12.7714 | <0.0001 |

| 2. Allergic - all |  |  |  |  |  |  |
| --- | --- | --- | --- | --- | --- | --- |
|  | exp(coef) | LCL | UCL | SE | Z | Pr(Z) |
| surgery | 1.3799 | 1.2562 | 1.5159 | 0.0479 | 6.7173 | <0.0001 |
| paternal age | 1.0061 | 0.9857 | 1.0269 | 0.0104 | 0.5832 | 0.5597 |
| maternal age | 1.0268 | 1.0032 | 1.0511 | 0.0118 | 2.2304 | 0.0257 |
| gestation length | 0.9972 | 0.9812 | 1.0134 | 0.0082 | -0.3397 | 0.7340 |
| maternal bleeding | 1.1426 | 1.0816 | 1.2070 | 0.0279 | 4.7651 | <0.0001 |
| fetal oxygen deprivation | 0.6034 | 0.3845 | 0.9468 | 0.2298 | -2.1975 | 0.0279 |
| pregnancy oedema | 1.2391 | 1.0974 | 1.3990 | 0.0619 | 3.4611 | 0.0005 |
| apgar5 score | 1.0825 | 1.0193 | 1.1495 | 0.0306 | 2.5846 | 0.0097 |
| birth weight | 0.9780 | 0.9632 | 0.9931 | 0.0078 | -2.8362 | 0.0045 |
| preexisting hypertension | 0.7289 | 0.4646 | 1.1435 | 0.2297 | -1.3763 | 0.1687 |
| preexisting diabetes | 1.0399 | 0.7932 | 1.3634 | 0.1381 | 0.2834 | 0.7768 |
| previous induced abortion | 1.0402 | 1.0012 | 1.0808 | 0.0195 | 2.0242 | 0.0429 |
| previous spontaneous abortion | 1.0433 | 1.0012 | 1.0872 | 0.0210 | 2.0175 | 0.0436 |
| education level | 1.0654 | 1.0467 | 1.0844 | 0.0090 | 7.0149 | <0.0001 |
| parental income | 1.0196 | 1.0006 | 1.0389 | 0.0095 | 2.0307 | 0.0422 |
| country | 1.3856 | 1.2911 | 1.4871 | 0.0360 | 9.0506 | <0.0001 |
| region Sjælland | 0.7125 | 0.6772 | 0.7497 | 0.0259 | -13.0736 | <0.0001 |
| region Syddanmark | 1.0327 | 0.9923 | 1.0747 | 0.0203 | 1.5842 | 0.1131 |
| region Midtjylland | 0.7876 | 0.7550 | 0.8217 | 0.0216 | -11.0424 | <0.0001 |
| region Nordjylland | 0.7121 | 0.6726 | 0.7540 | 0.0291 | -11.6537 | <0.0001 |
| mother with disorder | 1.8645 | 1.7454 | 1.9918 | 0.0336 | 18.5010 | <0.0001 |
| father with disorder | 1.7644 | 1.6250 | 1.9158 | 0.0420 | 13.5176 | <0.0001 |

Table S6
| 3. Allergic - rhinitis |  |  |  |  |  |  |
| --- | --- | --- | --- | --- | --- | --- |
|  | exp(coef) | LCL | UCL | SE | Z | Pr(Z) |
| surgery | 1.6148 | 1.4252 | 1.8296 | 0.0637 | 7.5203 | <0.0001 |
| paternal age | 1.0191 | 0.9898 | 1.0492 | 0.0148 | 1.2732 | 0.2029 |
| maternal age | 1.0643 | 1.0297 | 1.1001 | 0.0168 | 3.6957 | 0.0002 |
| gestation length | 0.9994 | 0.9771 | 1.0222 | 0.0115 | -0.0517 | 0.9587 |
| maternal bleeding | 1.0643 | 0.9828 | 1.1526 | 0.0406 | 1.5342 | 0.1249 |
| fetal oxygen deprivation | 0.5550 | 0.2884 | 1.0680 | 0.3339 | -1.7629 | 0.0779 |
| pregnancy oedema | 1.1629 | 0.9751 | 1.3870 | 0.0899 | 1.6795 | 0.0930 |
| apgar5 score | 1.0914 | 1.0044 | 1.1860 | 0.0423 | 2.0647 | 0.0389 |
| birth weight | 0.9888 | 0.9678 | 1.0103 | 0.0109 | -1.0195 | 0.3079 |
| preexisting hypertension | 0.6024 | 0.3010 | 1.2056 | 0.3539 | -1.4315 | 0.1522 |
| preexisting diabetes | 0.7667 | 0.4937 | 1.1908 | 0.2246 | -1.1823 | 0.2370 |
| previous induced abortion | 1.0042 | 0.9505 | 1.0610 | 0.0280 | 0.1525 | 0.8787 |
| previous spontaneous abortion | 1.0328 | 0.9744 | 1.0947 | 0.0297 | 1.0883 | 0.2764 |
| education level | 1.1294 | 1.1012 | 1.1582 | 0.0128 | 9.4599 | <0.0001 |
| parental income | 1.0258 | 0.9986 | 1.0538 | 0.0137 | 1.8620 | 0.0625 |
| country | 1.4185 | 1.2751 | 1.5780 | 0.0543 | 6.4295 | <0.0001 |
| region Sjælland | 0.9071 | 0.8400 | 0.9795 | 0.0391 | -2.4869 | 0.0128 |
| region Syddanmark | 1.7590 | 1.6605 | 1.8633 | 0.0293 | 19.2115 | <0.0001 |
| region Midtjylland | 1.1154 | 1.0473 | 1.1879 | 0.0321 | 3.3992 | 0.0006 |
| region Nordjylland | 1.2836 | 1.1897 | 1.3849 | 0.0387 | 6.4435 | <0.0001 |
| mother with disorder | 3.1455 | 2.8026 | 3.5304 | 0.0588 | 19.4586 | <0.0001 |
| father with disorder | 2.8219 | 2.4535 | 3.2456 | 0.0713 | 14.5365 | <0.0001 |

| 4. Allergic - conjunctivitis |  |  |  |  |  |  |
| --- | --- | --- | --- | --- | --- | --- |
|  | exp(coef) | LCL | UCL | SE | Z | Pr(Z) |
| surgery | 1.7514 | 1.3518 | 2.2691 | 0.1321 | 4.2421 | <0.0001 |
| paternal age | 1.0119 | 0.9514 | 1.0762 | 0.0314 | 0.3770 | 0.7061 |
| maternal age | 1.0713 | 0.9990 | 1.1489 | 0.0356 | 1.9319 | 0.0533 |
| gestation length | 1.0248 | 0.9770 | 1.0749 | 0.0243 | 1.0057 | 0.3145 |
| maternal bleeding | 1.2086 | 1.0295 | 1.4188 | 0.0818 | 2.3155 | 0.0205 |
| fetal oxygen deprivation | 0.0000 | 0.0000 | Inf | 370.1012 | -0.0344 | 0.9724 |
| pregnancy oedema | 1.4792 | 1.0510 | 2.0818 | 0.1743 | 2.2457 | 0.0247 |
| apgar5 score | 1.0437 | 0.8738 | 1.2466 | 0.0906 | 0.4722 | 0.6367 |
| birth weight | 1.0132 | 0.9678 | 1.0606 | 0.0233 | 0.5616 | 0.5743 |
| preexisting hypertension | 0.3226 | 0.0453 | 2.2934 | 1.0006 | -1.1304 | 0.2582 |
| preexisting diabetes | 1.2298 | 0.5832 | 2.5932 | 0.3806 | 0.5434 | 0.5868 |
| previous induced abortion | 1.1018 | 0.9837 | 1.2340 | 0.0578 | 1.6766 | 0.0936 |
| previous spontaneous abortion | 1.1712 | 1.0398 | 1.3192 | 0.0607 | 2.6032 | 0.0092 |
| education level | 1.0673 | 1.0116 | 1.1261 | 0.0273 | 2.3824 | 0.0171 |
| parental income | 1.0253 | 0.9680 | 1.0860 | 0.0293 | 0.8523 | 0.3940 |
| country | 1.5564 | 1.2648 | 1.9152 | 0.1058 | 4.1797 | <0.0001 |
| region Sjælland | 0.6708 | 0.5653 | 0.7960 | 0.0873 | -4.5727 | <0.0001 |
| region Syddanmark | 1.8783 | 1.6752 | 2.1060 | 0.0583 | 10.7975 | <0.0001 |
| region Midtjylland | 0.6978 | 0.6035 | 0.8067 | 0.0740 | -4.8608 | <0.0001 |
| region Nordjylland | 0.9588 | 0.8091 | 1.1361 | 0.0865 | -0.4857 | 0.6271 |
| mother with disorder | 1.6499 | 0.7851 | 3.4676 | 0.3789 | 1.3215 | 0.1863 |
| father with disorder | 2.8925 | 1.3758 | 6.0812 | 0.3791 | 2.8014 | 0.0050 |

Table S6
| 5. Allergic - eczema/dermatitis |  |  |  |  |  |  |
| --- | --- | --- | --- | --- | --- | --- |
|  | exp(coef) | LCL | UCL | SE | Z | Pr(Z) |
| surgery | 1.2997 | 1.0782 | 1.5666 | 0.0953 | 2.7505 | 0.0059 |
| paternal age | 0.9893 | 0.9515 | 1.0287 | 0.0199 | -0.5354 | 0.5923 |
| maternal age | 0.9966 | 0.9531 | 1.0422 | 0.0227 | -0.1455 | 0.8842 |
| gestation length | 1.0211 | 0.9901 | 1.0531 | 0.0157 | 1.3314 | 0.1830 |
| maternal bleeding | 1.2056 | 1.0883 | 1.3356 | 0.0522 | 3.5796 | 0.0003 |
| fetal oxygen deprivation | 0.6818 | 0.3057 | 1.5204 | 0.4091 | -0.9358 | 0.3493 |
| pregnancy oedema | 1.2988 | 1.0354 | 1.6291 | 0.1156 | 2.2610 | 0.0237 |
| apgar5 score | 1.0580 | 0.9411 | 1.1895 | 0.0597 | 0.9446 | 0.3448 |
| birth weight | 0.9816 | 0.9534 | 1.0107 | 0.0148 | -1.2429 | 0.2138 |
| preexisting hypertension | 0.7051 | 0.2931 | 1.6961 | 0.4478 | -0.7801 | 0.4353 |
| preexisting diabetes | 1.0918 | 0.6561 | 1.8166 | 0.2597 | 0.3381 | 0.7352 |
| previous induced abortion | 1.0789 | 1.0044 | 1.1590 | 0.0365 | 2.0816 | 0.0373 |
| previous spontaneous abortion | 1.0723 | 0.9920 | 1.1591 | 0.0397 | 1.7594 | 0.0785 |
| education level | 1.0411 | 1.0066 | 1.0768 | 0.0171 | 2.3460 | 0.0189 |
| parental income | 1.0582 | 1.0212 | 1.0966 | 0.0181 | 3.1159 | 0.0018 |
| country | 1.2608 | 1.0985 | 1.4471 | 0.0703 | 3.2970 | 0.0009 |
| region Sjælland | 0.6128 | 0.5559 | 0.6755 | 0.0497 | -9.8518 | <0.0001 |
| region Syddanmark | 0.9003 | 0.8352 | 0.9704 | 0.0382 | -2.7429 | 0.0060 |
| region Midtjylland | 0.8009 | 0.7416 | 0.8648 | 0.0391 | -5.6631 | <0.0001 |
| region Nordjylland | 0.4031 | 0.3538 | 0.4594 | 0.0666 | -13.6332 | <0.0001 |
| mother with disorder | 2.4876 | 2.0684 | 2.9919 | 0.0941 | 9.6778 | <0.0001 |
| father with disorder | 2.0043 | 1.5079 | 2.6641 | 0.1451 | 4.7886 | <0.0001 |

| 6. Allergic - urticaria/angiodema |  |  |  |  |  |  |
| --- | --- | --- | --- | --- | --- | --- |
|  | exp(coef) | LCL | UCL | SE | Z | Pr(Z) |
| surgery | 1.2233 | 1.0186 | 1.4691 | 0.0934 | 2.1572 | 0.0309 |
| paternal age | 1.0063 | 0.9680 | 1.0461 | 0.0197 | 0.3201 | 0.7488 |
| maternal age | 0.9894 | 0.9464 | 1.0344 | 0.0226 | -0.4679 | 0.6398 |
| gestation length | 0.9903 | 0.9597 | 1.0218 | 0.0160 | -0.6076 | 0.5434 |
| maternal bleeding | 1.2773 | 1.1548 | 1.4127 | 0.0514 | 4.7600 | <0.0001 |
| fetal oxygen deprivation | 0.5341 | 0.2001 | 1.4256 | 0.5008 | -1.2519 | 0.2106 |
| pregnancy oedema | 1.5592 | 1.2664 | 1.9197 | 0.1061 | 4.1854 | <0.0001 |
| apgar5 score | 1.1153 | 0.9935 | 1.2520 | 0.0589 | 1.8507 | 0.0642 |
| birth weight | 0.9683 | 0.9400 | 0.9975 | 0.0151 | -2.1207 | 0.0339 |
| preexisting hypertension | 0.7246 | 0.3011 | 1.7436 | 0.4480 | -0.7189 | 0.4721 |
| preexisting diabetes | 1.3165 | 0.8266 | 2.0970 | 0.2374 | 1.1581 | 0.2468 |
| previous induced abortion | 1.0613 | 0.9869 | 1.1414 | 0.0370 | 1.6055 | 0.1083 |
| previous spontaneous abortion | 1.0461 | 0.9663 | 1.1325 | 0.0404 | 1.1153 | 0.2647 |
| education level | 1.0288 | 0.9947 | 1.0641 | 0.0172 | 1.6529 | 0.0983 |
| parental income | 0.9741 | 0.9400 | 1.0094 | 0.0181 | -1.4421 | 0.1492 |
| country | 1.4941 | 1.3234 | 1.6868 | 0.0619 | 6.4866 | <0.0001 |
| region Sjælland | 0.6060 | 0.5530 | 0.6640 | 0.0466 | -10.7258 | <0.0001 |
| region Syddanmark | 0.6497 | 0.6015 | 0.7018 | 0.0393 | -10.9571 | <0.0001 |
| region Midtjylland | 0.4924 | 0.4528 | 0.5354 | 0.0427 | -16.5806 | <0.0001 |
| region Nordjylland | 0.3630 | 0.3197 | 0.4121 | 0.0647 | -15.6455 | <0.0001 |
| mother with disorder | 1.9275 | 1.5848 | 2.3443 | 0.0998 | 6.5704 | <0.0001 |
| father with disorder | 1.4300 | 1.0818 | 1.8903 | 0.1423 | 2.5128 | 0.0119 |

Table S6
| 7. Skin - all | exp(coef) | LCL | UCL | SE | Z | Pr(Z) |
| --- | --- | --- | --- | --- | --- | --- |
| surgery | 1.2386 | 1.1730 | 1.3078 | 0.0277 | 7.7116 | <0.0001 |
| paternal age | 0.9884 | 0.9769 | 1.0000 | 0.0059 | -1.9541 | 0.0506 |
| maternal age | 0.9546 | 0.9418 | 0.9675 | 0.0068 | -6.7703 | <0.0001 |
| gestation length | 0.9965 | 0.9872 | 1.0059 | 0.0047 | -0.7222 | 0.4701 |
| maternal bleeding | 1.0696 | 1.0353 | 1.1051 | 0.0166 | 4.0493 | <0.0001 |
| fetal oxygen deprivation | 0.8429 | 0.6874 | 1.0336 | 0.1040 | -1.6420 | 0.1005 |
| pregnancy oedema | 1.1263 | 1.0492 | 1.2090 | 0.0361 | 3.2901 | 0.0010 |
| apgar5 score | 1.0963 | 1.0584 | 1.1356 | 0.0179 | 5.1213 | <0.0001 |
| birth weight | 1.0224 | 1.0134 | 1.0315 | 0.0045 | 4.9134 | <0.0001 |
| preexisting hypertension | 1.0731 | 0.8603 | 1.3385 | 0.1127 | 0.6261 | 0.5311 |
| preexisting diabetes | 1.1615 | 1.0079 | 1.3386 | 0.0723 | 2.0690 | 0.0385 |
| previous induced abortion | 1.0888 | 1.0652 | 1.1129 | 0.0111 | 7.6198 | <0.0001 |
| previous spontaneous abortion | 1.0468 | 1.0220 | 1.0722 | 0.0122 | 3.7382 | 0.0001 |
| education level | 0.9623 | 0.9526 | 0.9722 | 0.0051 | -7.3767 | <0.0001 |
| parental income | 0.9707 | 0.9602 | 0.9813 | 0.0055 | -5.3601 | <0.0001 |
| country | 1.2997 | 1.2489 | 1.3527 | 0.0203 | 12.8804 | <0.0001 |
| region Sjælland | 0.7951 | 0.7730 | 0.8178 | 0.0143 | -15.9584 | <0.0001 |
| region Syddanmark | 0.8676 | 0.8470 | 0.8886 | 0.0122 | -11.6087 | <0.0001 |
| region Midtjylland | 0.8544 | 0.8342 | 0.8751 | 0.0121 | -12.9038 | <0.0001 |
| region Nordjylland | 0.5976 | 0.5772 | 0.6188 | 0.0177 | -29.0272 | <0.0001 |
| mother with disorder | 1.3345 | 1.3016 | 1.3683 | 0.0127 | 22.6340 | <0.0001 |
| father with disorder | 1.2917 | 1.2589 | 1.3254 | 0.0131 | 19.5177 | <0.0001 |

| 8. Autoimmune - all | exp(coef) | LCL | UCL | SE | Z | Pr(Z) |
| --- | --- | --- | --- | --- | --- | --- |
| surgery | 1.1928 | 1.0955 | 1.2987 | 0.0433 | 4.0637 | <0.0001 |
| paternal age | 0.9839 | 0.9666 | 1.0015 | 0.0090 | -1.7920 | 0.0731 |
| maternal age | 0.9715 | 0.9521 | 0.9914 | 0.0103 | -2.7895 | 0.0052 |
| gestation length | 0.9934 | 0.9796 | 1.0074 | 0.0071 | -0.9192 | 0.3579 |
| maternal bleeding | 1.0847 | 1.0333 | 1.1388 | 0.0248 | 3.2813 | 0.0010 |
| fetal oxygen deprivation | 1.0483 | 0.7891 | 1.3925 | 0.1448 | 0.3256 | 0.7446 |
| pregnancy oedema | 1.2107 | 1.0915 | 1.3429 | 0.0528 | 3.6166 | 0.0002 |
| apgar5 score | 0.9841 | 0.9316 | 1.0396 | 0.0279 | -0.5697 | 0.5688 |
| birth weight | 0.9950 | 0.9819 | 1.0082 | 0.0067 | -0.7438 | 0.4569 |
| preexisting hypertension | 1.0326 | 0.7476 | 1.4263 | 0.1647 | 0.1951 | 0.8452 |
| preexisting diabetes | 1.5553 | 1.3135 | 1.8415 | 0.0861 | 5.1250 | <0.0001 |
| previous induced abortion | 0.9742 | 0.9419 | 1.0077 | 0.0172 | -1.5116 | 0.1306 |
| previous spontaneous abortion | 1.0520 | 1.0153 | 1.0901 | 0.0181 | 2.8010 | 0.0050 |
| education level | 0.9911 | 0.9760 | 1.0064 | 0.0078 | -1.1384 | 0.2549 |
| parental income | 1.0117 | 0.9953 | 1.0284 | 0.0083 | 1.3997 | 0.1615 |
| country | 0.8921 | 0.8315 | 0.9570 | 0.0358 | -3.1834 | 0.0014 |
| region Sjælland | 0.9654 | 0.9266 | 1.0058 | 0.0209 | -1.6790 | 0.0931 |
| region Syddanmark | 0.9770 | 0.9424 | 1.0129 | 0.0184 | -1.2608 | 0.2073 |
| region Midtjylland | 0.9182 | 0.8854 | 0.9522 | 0.0185 | -4.5920 | <0.0001 |
| region Nordjylland | 0.7767 | 0.7396 | 0.8156 | 0.0249 | -10.1193 | <0.0001 |
| mother with disorder | 1.5299 | 1.4688 | 1.5935 | 0.0207 | 20.4572 | <0.0001 |
| father with disorder | 1.4422 | 1.3817 | 1.5053 | 0.0218 | 16.7602 | <0.0001 |

Table S6
| 9. Respiratory - all |  |  |  |  |  |  |
| --- | --- | --- | --- | --- | --- | --- |
|  | exp(coef) | LCL | UCL | SE | Z | Pr(Z) |
| surgery | 1.4886 | 1.1309 | 1.9594 | 0.1402 | 2.8377 | 0.0045 |
| paternal age | 0.9996 | 0.9877 | 1.0116 | 0.0060 | -0.0615 | 0.9509 |
| maternal age | 0.9595 | 0.9464 | 0.9728 | 0.0070 | -5.8935 | <0.0001 |
| gestation length | 0.9852 | 0.9757 | 0.9947 | 0.0049 | -3.0253 | 0.0024 |
| maternal bleeding | 1.0834 | 1.0474 | 1.1206 | 0.0172 | 4.6513 | <0.0001 |
| fetal oxygen deprivation | 0.9176 | 0.7404 | 1.1371 | 0.1094 | -0.7855 | 0.4321 |
| pregnancy oedema | 1.1050 | 1.0241 | 1.1923 | 0.0388 | 2.5740 | 0.0100 |
| apgar5 score | 1.0527 | 1.0146 | 1.0923 | 0.0188 | 2.7320 | 0.0062 |
| birth weight | 1.0015 | 0.9925 | 1.0106 | 0.0046 | 0.3321 | 0.7397 |
| preexisting hypertension | 1.1662 | 0.9424 | 1.4432 | 0.1087 | 1.4145 | 0.1572 |
| preexisting diabetes | 1.0094 | 0.8610 | 1.1833 | 0.0811 | 0.1158 | 0.9077 |
| previous induced abortion | 1.0560 | 1.0324 | 1.0800 | 0.0114 | 4.7433 | <0.0001 |
| previous spontaneous abortion | 1.0331 | 1.0081 | 1.0586 | 0.0124 | 2.6136 | 0.0089 |
| education level | 1.0008 | 0.9904 | 1.0112 | 0.0052 | 0.1548 | 0.8769 |
| parental income | 0.9741 | 0.9634 | 0.9850 | 0.0056 | -4.6150 | <0.0001 |
| country | 1.2267 | 1.1768 | 1.2787 | 0.0211 | 9.6487 | <0.0001 |
| region Sjælland | 0.8114 | 0.7886 | 0.8348 | 0.0145 | -14.3841 | <0.0001 |
| region Syddanmark | 0.8771 | 0.8561 | 0.8986 | 0.0123 | -10.5862 | <0.0001 |
| region Midtjylland | 0.7360 | 0.7179 | 0.7546 | 0.0127 | -24.0978 | <0.0001 |
| region Nordjylland | 0.7285 | 0.7053 | 0.7524 | 0.0164 | -19.1977 | <0.0001 |
| mother with disorder | 1.3296 | 1.3013 | 1.3586 | 0.0110 | 25.8926 | <0.0001 |
| father with disorder | 1.2407 | 1.2140 | 1.2680 | 0.0111 | 19.4280 | <0.0001 |

| 10. Respiratory - upper |  |  |  |  |  |  |
| --- | --- | --- | --- | --- | --- | --- |
|  | exp(coef) | LCL | UCL | SE | Z | Pr(Z) |
| surgery | 1.9998 | 1.5191 | 2.6326 | 0.1402 | 4.9410 | <0.0001 |
| paternal age | 1.0013 | 0.9866 | 1.0162 | 0.0075 | 0.1786 | 0.8581 |
| maternal age | 0.9668 | 0.9504 | 0.9833 | 0.0086 | -3.8879 | 0.0001 |
| gestation length | 0.9923 | 0.9806 | 1.0041 | 0.0060 | -1.2674 | 0.2049 |
| maternal bleeding | 1.0761 | 1.0324 | 1.1217 | 0.0211 | 3.4681 | 0.0005 |
| fetal oxygen deprivation | 0.8362 | 0.6315 | 1.1073 | 0.1432 | -1.2482 | 0.2119 |
| pregnancy oedema | 1.1372 | 1.0366 | 1.2475 | 0.0472 | 2.7225 | 0.0064 |
| apgar5 score | 1.0902 | 1.0427 | 1.1399 | 0.0227 | 3.8019 | 0.0001 |
| birth weight | 1.0024 | 0.9913 | 1.0137 | 0.0056 | 0.4308 | 0.6665 |
| preexisting hypertension | 1.1318 | 0.8663 | 1.4787 | 0.1363 | 0.9082 | 0.3637 |
| preexisting diabetes | 0.9439 | 0.7702 | 1.1568 | 0.1037 | -0.5558 | 0.5783 |
| previous induced abortion | 1.0949 | 1.0652 | 1.1254 | 0.0140 | 6.4613 | <0.0001 |
| previous spontaneous abortion | 1.0316 | 1.0010 | 1.0632 | 0.0153 | 2.0251 | 0.0428 |
| education level | 1.0127 | 0.9998 | 1.0257 | 0.0065 | 1.9369 | 0.0527 |
| parental income | 0.9900 | 0.9765 | 1.0036 | 0.0069 | -1.4359 | 0.1510 |
| country | 1.4063 | 1.3386 | 1.4775 | 0.0251 | 13.5348 | <0.0001 |
| region Sjælland | 0.8276 | 0.7990 | 0.8572 | 0.0179 | -10.5500 | <0.0001 |
| region Syddanmark | 0.9339 | 0.9066 | 0.9620 | 0.0151 | -4.5181 | <0.0001 |
| region Midtjylland | 0.7395 | 0.7169 | 0.7628 | 0.0158 | -19.0335 | <0.0001 |
| region Nordjylland | 0.6998 | 0.6716 | 0.7291 | 0.0209 | -17.0259 | <0.0001 |
| mother with disorder | 1.3807 | 1.3371 | 1.4257 | 0.0163 | 19.7293 | <0.0001 |
| father with disorder | 1.2904 | 1.2479 | 1.3343 | 0.0170 | 14.9377 | <0.0001 |

Table S6
| 11. Respiratory - lower | exp(coef) | LCL | UCL | SE | Z | Pr(Z) |
| --- | --- | --- | --- | --- | --- | --- |
| surgery | 1.6187 | 1.4204 | 1.8446 | 0.0666 | 7.2258 | <0.0001 |
| paternal age | 1.0031 | 0.9730 | 1.0342 | 0.0155 | 0.2016 | 0.8401 |
| maternal age | 0.9158 | 0.8840 | 0.9487 | 0.0180 | -4.8791 | <0.0001 |
| gestation length | 0.9408 | 0.9176 | 0.9646 | 0.0127 | -4.7868 | <0.0001 |
| maternal bleeding | 1.1543 | 1.0620 | 1.2546 | 0.0425 | 3.3752 | 0.0007 |
| fetal oxygen deprivation | 0.9896 | 0.5850 | 1.6740 | 0.2682 | -0.0388 | 0.9690 |
| pregnancy oedema | 1.1558 | 0.9588 | 1.3934 | 0.0953 | 1.5190 | 0.1287 |
| apgar5 score | 1.1228 | 1.0242 | 1.2309 | 0.0468 | 2.4716 | 0.0134 |
| birth weight | 0.9824 | 0.9596 | 1.0058 | 0.0119 | -1.4761 | 0.1399 |
| preexisting hypertension | 1.2103 | 0.7015 | 2.0879 | 0.2782 | 0.6861 | 0.4926 |
| preexisting diabetes | 1.4144 | 1.0071 | 1.9864 | 0.1732 | 2.0010 | 0.0453 |
| previous induced abortion | 1.0015 | 0.9439 | 1.0626 | 0.0302 | 0.0497 | 0.9603 |
| previous spontaneous abortion | 1.1303 | 1.0622 | 1.2029 | 0.0317 | 3.8630 | 0.0001 |
| education level | 0.9675 | 0.9418 | 0.9939 | 0.0137 | -2.4023 | 0.0162 |
| parental income | 0.9277 | 0.9014 | 0.9547 | 0.0146 | -5.1210 | <0.0001 |
| country | 1.1097 | 0.9986 | 1.2332 | 0.0538 | 1.9339 | 0.0531 |
| region Sjælland | 0.7978 | 0.7428 | 0.8569 | 0.0364 | -6.1951 | <0.0001 |
| region Syddanmark | 0.7261 | 0.6812 | 0.7740 | 0.0325 | -9.8266 | <0.0001 |
| region Midtjylland | 0.6505 | 0.6093 | 0.6945 | 0.0333 | -12.8737 | <0.0001 |
| region Nordjylland | 0.6551 | 0.6021 | 0.7128 | 0.0430 | -9.8243 | <0.0001 |
| mother with disorder | 1.5851 | 1.4441 | 1.7399 | 0.0475 | 9.6899 | <0.0001 |
| father with disorder | 1.3628 | 1.2430 | 1.4941 | 0.0469 | 6.5931 | <0.0001 |

| 12. Respiratory - chronic lower | exp(coef) | LCL | UCL | SE | Z | Pr(Z) |
| --- | --- | --- | --- | --- | --- | --- |
| surgery | 1.4465 | 1.3335 | 1.5690 | 0.0414 | 8.8969 | <0.0001 |
| paternal age | 0.9980 | 0.9807 | 1.0156 | 0.0089 | -0.2172 | 0.8280 |
| maternal age | 0.9755 | 0.9560 | 0.9954 | 0.0102 | -2.4069 | 0.0160 |
| gestation length | 0.9670 | 0.9535 | 0.9806 | 0.0071 | -4.7003 | <0.0001 |
| maternal bleeding | 1.1178 | 1.0655 | 1.1726 | 0.0244 | 4.5602 | <0.0001 |
| fetal oxygen deprivation | 1.0408 | 0.7597 | 1.4261 | 0.1606 | 0.2493 | 0.8030 |
| pregnancy oedema | 1.1233 | 1.0023 | 1.2590 | 0.0581 | 2.0000 | 0.0454 |
| apgar5 score | 1.0730 | 1.0185 | 1.1303 | 0.0265 | 2.6522 | 0.0079 |
| birth weight | 0.9772 | 0.9643 | 0.9903 | 0.0068 | -3.3801 | 0.0007 |
| preexisting hypertension | 0.8698 | 0.6113 | 1.2376 | 0.1799 | -0.7747 | 0.4384 |
| preexisting diabetes | 0.9545 | 0.7553 | 1.2062 | 0.1194 | -0.3899 | 0.6965 |
| previous induced abortion | 1.0226 | 0.9891 | 1.0572 | 0.0169 | 1.3194 | 0.1870 |
| previous spontaneous abortion | 1.0530 | 1.0158 | 1.0914 | 0.0183 | 2.8211 | 0.0047 |
| education level | 1.0239 | 1.0081 | 1.0400 | 0.0079 | 2.9890 | 0.0027 |
| parental income | 0.9431 | 0.9276 | 0.9589 | 0.0084 | -6.9059 | <0.0001 |
| country | 0.9559 | 0.8960 | 1.0197 | 0.0329 | -1.3669 | 0.1716 |
| region Sjælland | 0.8193 | 0.7853 | 0.8547 | 0.0215 | -9.2252 | <0.0001 |
| region Syddanmark | 0.9663 | 0.9325 | 1.0013 | 0.0181 | -1.8852 | 0.0593 |
| region Midtjylland | 0.7513 | 0.7235 | 0.7802 | 0.0192 | -14.8478 | <0.0001 |
| region Nordjylland | 0.8561 | 0.8173 | 0.8967 | 0.0236 | -6.5653 | <0.0001 |
| mother with disorder | 1.9777 | 1.8882 | 2.0715 | 0.0236 | 28.8591 | <0.0001 |
| father with disorder | 1.7727 | 1.6831 | 1.8671 | 0.0264 | 21.6381 | <0.0001 |

Table S6
| 13. Respiratory - asthma |  |  |  |  |  |  |
| --- | --- | --- | --- | --- | --- | --- |
|  | exp(coef) | LCL | UCL | SE | Z | Pr(Z) |
| surgery | 1.4513 | 1.3374 | 1.5748 | 0.0416 | 8.9332 | <0.0001 |
| paternal age | 1.0082 | 0.9906 | 1.0261 | 0.0089 | 0.9119 | 0.3618 |
| maternal age | 0.9757 | 0.9560 | 0.9957 | 0.0103 | -2.3732 | 0.0176 |
| gestation length | 0.9623 | 0.9488 | 0.9759 | 0.0071 | -5.3570 | <0.0001 |
| maternal bleeding | 1.1237 | 1.0710 | 1.1789 | 0.0244 | 4.7644 | <0.0001 |
| fetal oxygen deprivation | 1.0800 | 0.7913 | 1.4739 | 0.1586 | 0.4853 | 0.6274 |
| pregnancy oedema | 1.1080 | 0.9869 | 1.2439 | 0.0590 | 1.7372 | 0.0823 |
| apgar5 score | 1.0726 | 1.0180 | 1.1302 | 0.0266 | 2.6305 | 0.0085 |
| birth weight | 0.9758 | 0.9628 | 0.9890 | 0.0068 | -3.5724 | 0.0003 |
| preexisting hypertension | 0.8696 | 0.6111 | 1.2373 | 0.1799 | -0.7763 | 0.4375 |
| preexisting diabetes | 0.9582 | 0.7582 | 1.2109 | 0.1194 | -0.3573 | 0.7208 |
| previous induced abortion | 1.0245 | 0.9908 | 1.0594 | 0.0170 | 1.4236 | 0.1545 |
| previous spontaneous abortion | 1.0565 | 1.0191 | 1.0953 | 0.0184 | 2.9903 | 0.0027 |
| education level | 1.0159 | 1.0002 | 1.0319 | 0.0079 | 1.9914 | 0.0464 |
| parental income | 0.9375 | 0.9219 | 0.9532 | 0.0085 | -7.5791 | <0.0001 |
| country | 0.9329 | 0.8740 | 0.9957 | 0.0332 | -2.0893 | 0.0366 |
| region Sjælland | 0.8203 | 0.7861 | 0.8560 | 0.0217 | -9.1132 | <0.0001 |
| region Syddanmark | 0.9636 | 0.9297 | 0.9987 | 0.0182 | -2.0269 | 0.0426 |
| region Midtjylland | 0.7551 | 0.7271 | 0.7843 | 0.0193 | -14.5290 | <0.0001 |
| region Nordjylland | 0.8550 | 0.8161 | 0.8958 | 0.0237 | -6.5858 | <0.0001 |
| mother with disorder | 2.1842 | 2.0761 | 2.2978 | 0.0258 | 30.1915 | <0.0001 |
| father with disorder | 2.0609 | 1.9372 | 2.1923 | 0.0315 | 22.9156 | <0.0001 |

| 14. Respiratory - influenza |  |  |  |  |  |  |
| --- | --- | --- | --- | --- | --- | --- |
|  | exp(coef) | LCL | UCL | SE | Z | Pr(Z) |
| surgery | 1.4652 | 1.1845 | 1.8124 | 0.1085 | 3.5204 | 0.0004 |
| paternal age | 1.0202 | 0.9726 | 1.0701 | 0.0243 | 0.8219 | 0.4111 |
| maternal age | 0.9128 | 0.8634 | 0.9651 | 0.0283 | -3.2111 | 0.0013 |
| gestation length | 0.9109 | 0.8753 | 0.9479 | 0.0203 | -4.5871 | <0.0001 |
| maternal bleeding | 1.0735 | 0.9375 | 1.2293 | 0.0691 | 1.0268 | 0.3044 |
| fetal oxygen deprivation | 1.9432 | 1.0070 | 3.7499 | 0.3354 | 1.9807 | 0.0476 |
| pregnancy oedema | 1.3781 | 1.0339 | 1.8368 | 0.1465 | 2.1877 | 0.0286 |
| apgar5 score | 1.1003 | 0.9500 | 1.2744 | 0.0749 | 1.2756 | 0.2020 |
| birth weight | 0.9716 | 0.9355 | 1.0091 | 0.0193 | -1.4885 | 0.1365 |
| preexisting hypertension | 2.8172 | 1.5926 | 4.9834 | 0.2910 | 3.5592 | 0.0003 |
| preexisting diabetes | 1.3105 | 0.7567 | 2.2694 | 0.2801 | 0.9653 | 0.3343 |
| previous induced abortion | 1.0225 | 0.9306 | 1.1235 | 0.0480 | 0.4634 | 0.6430 |
| previous spontaneous abortion | 1.0899 | 0.9852 | 1.2056 | 0.0514 | 1.6719 | 0.0945 |
| education level | 0.9502 | 0.9107 | 0.9913 | 0.0216 | -2.3608 | 0.0182 |
| parental income | 0.8971 | 0.8570 | 0.9392 | 0.0233 | -4.6444 | <0.0001 |
| country | 1.9293 | 1.6906 | 2.2018 | 0.0673 | 9.7526 | <0.0001 |
| region Sjælland | 0.7028 | 0.6278 | 0.7868 | 0.0575 | -6.1223 | <0.0001 |
| region Syddanmark | 0.5771 | 0.5202 | 0.6403 | 0.0529 | -10.3719 | <0.0001 |
| region Midtjylland | 0.5170 | 0.4643 | 0.5756 | 0.0547 | -12.0414 | <0.0001 |
| region Nordjylland | 0.4885 | 0.4219 | 0.5656 | 0.0747 | -9.5847 | <0.0001 |
| mother with disorder | 2.5029 | 1.9371 | 3.2339 | 0.1307 | 7.0172 | <0.0001 |
| father with disorder | 1.7058 | 1.1888 | 2.4475 | 0.1842 | 2.8990 | 0.0037 |

Table S6
| 15. Respiratory - pneumonia | exp(coef) | LCL | UCL | SE | Z | Pr(Z) |
| --- | --- | --- | --- | --- | --- | --- |
| surgery | 1.6536 | 1.4459 | 1.8912 | 0.0684 | 7.3452 | <0.0001 |
| paternal age | 0.9994 | 0.9680 | 1.0319 | 0.0163 | -0.0319 | 0.9745 |
| maternal age | 0.9195 | 0.8861 | 0.9541 | 0.0188 | -4.4449 | <0.0001 |
| gestation length | 0.9415 | 0.9173 | 0.9663 | 0.0132 | -4.5359 | <0.0001 |
| maternal bleeding | 1.1920 | 1.0939 | 1.2988 | 0.0437 | 4.0101 | <0.0001 |
| fetal oxygen deprivation | 0.7814 | 0.4197 | 1.4549 | 0.3171 | -0.7775 | 0.4368 |
| pregnancy oedema | 1.1725 | 0.9649 | 1.4247 | 0.0994 | 1.6006 | 0.1094 |
| apgar5 score | 1.1345 | 1.0314 | 1.2478 | 0.0485 | 2.5983 | 0.0093 |
| birth weight | 0.9902 | 0.9662 | 1.0149 | 0.0125 | -0.7787 | 0.4361 |
| preexisting hypertension | 1.3374 | 0.7752 | 2.3076 | 0.2782 | 1.0449 | 0.2960 |
| preexisting diabetes | 1.3807 | 0.9675 | 1.9705 | 0.1814 | 1.7780 | 0.0753 |
| previous induced abortion | 0.9952 | 0.9354 | 1.0589 | 0.0316 | -0.1507 | 0.8801 |
| previous spontaneous abortion | 1.1223 | 1.0514 | 1.1979 | 0.0332 | 3.4694 | 0.0005 |
| education level | 0.9627 | 0.9359 | 0.9902 | 0.0144 | -2.6368 | 0.0083 |
| parental income | 0.9391 | 0.9112 | 0.9677 | 0.0153 | -4.0913 | <0.0001 |
| country | 1.1092 | 0.9936 | 1.2383 | 0.0561 | 1.8459 | 0.0648 |
| region Sjælland | 0.7750 | 0.7191 | 0.8353 | 0.0381 | -6.6694 | <0.0001 |
| region Syddanmark | 0.7303 | 0.6835 | 0.7803 | 0.0337 | -9.3001 | <0.0001 |
| region Midtjylland | 0.6380 | 0.5957 | 0.6833 | 0.0349 | -12.8422 | <0.0001 |
| region Nordjylland | 0.6024 | 0.5502 | 0.6596 | 0.0462 | -10.9559 | <0.0001 |
| mother with disorder | 1.5898 | 1.4350 | 1.7613 | 0.0522 | 8.8721 | <0.0001 |
| father with disorder | 1.3477 | 1.2190 | 1.4899 | 0.0511 | 5.8299 | <0.0001 |

| 16. Respiratory - COPD | exp(coef) | LCL | UCL | SE | Z | Pr(Z) |
| --- | --- | --- | --- | --- | --- | --- |
| surgery | 2.1195 | 1.5349 | 2.9268 | 0.1646 | 4.5620 | <0.0001 |
| paternal age | 0.9527 | 0.8649 | 1.0495 | 0.0493 | -0.9807 | 0.3267 |
| maternal age | 0.9454 | 0.8460 | 1.0566 | 0.0567 | -0.9888 | 0.3227 |
| gestation length | 1.0242 | 0.9455 | 1.1094 | 0.0407 | 0.5867 | 0.5573 |
| maternal bleeding | 1.2796 | 0.9911 | 1.6520 | 0.1303 | 1.8918 | 0.0585 |
| fetal oxygen deprivation | 0.6305 | 0.0883 | 4.5002 | 1.0026 | -0.4598 | 0.6456 |
| pregnancy oedema | 1.5897 | 0.9917 | 2.5483 | 0.2407 | 1.9255 | 0.0541 |
| apgar5 score | 1.1194 | 0.8310 | 1.5080 | 0.1520 | 0.7424 | 0.4578 |
| birth weight | 0.8669 | 0.8040 | 0.9347 | 0.0384 | -3.7147 | 0.0002 |
| preexisting hypertension | 0.0000 | 0.0000 | Inf | 699.1457 | -0.0186 | 0.9851 |
| preexisting diabetes | 1.1143 | 0.2766 | 4.4894 | 0.7109 | 0.1523 | 0.8789 |
| previous induced abortion | 1.1268 | 0.9378 | 1.3540 | 0.0936 | 1.2752 | 0.2022 |
| previous spontaneous abortion | 1.0182 | 0.8283 | 1.2515 | 0.1052 | 0.1713 | 0.8639 |
| education level | 0.9994 | 0.9169 | 1.0892 | 0.0439 | -0.0136 | 0.9891 |
| parental income | 0.8112 | 0.7408 | 0.8883 | 0.0463 | -4.5148 | <0.0001 |
| country | 1.1506 | 0.8256 | 1.6037 | 0.1693 | 0.8285 | 0.4073 |
| region Sjælland | 0.8304 | 0.6644 | 1.0378 | 0.1137 | -1.6337 | 0.1023 |
| region Syddanmark | 0.6986 | 0.5690 | 0.8576 | 0.1046 | -3.4268 | 0.0006 |
| region Midtjylland | 0.5772 | 0.4634 | 0.7190 | 0.1120 | -4.9042 | <0.0001 |
| region Nordjylland | 0.7104 | 0.5463 | 0.9238 | 0.1340 | -2.5508 | 0.0107 |
| mother with disorder | 1.9286 | 1.3686 | 2.7177 | 0.1749 | 3.7534 | 0.0001 |
| father with disorder | 1.6667 | 1.1832 | 2.3478 | 0.1748 | 2.9226 | 0.0034 |

Table S6
| 17. Digestive - all | exp(coef) | LCL | UCL | SE | Z | Pr(Z) |
| --- | --- | --- | --- | --- | --- | --- |
| surgery | 1.1319 | 1.0747 | 1.1921 | 0.0264 | 4.6864 | <0.0001 |
| paternal age | 0.9811 | 0.9705 | 0.9918 | 0.0055 | -3.4424 | 0.0005 |
| maternal age | 0.9174 | 0.9060 | 0.9289 | 0.0063 | -13.5176 | <0.0001 |
| gestation length | 0.9826 | 0.9741 | 0.9912 | 0.0044 | -3.9438 | <0.0001 |
| maternal bleeding | 1.0647 | 1.0329 | 1.0974 | 0.0154 | 4.0634 | <0.0001 |
| fetal oxygen deprivation | 1.0383 | 0.8749 | 1.2323 | 0.0873 | 0.4308 | 0.6665 |
| pregnancy oedema | 1.1367 | 1.0668 | 1.2112 | 0.0323 | 3.9597 | <0.0001 |
| apgar5 score | 1.0578 | 1.0234 | 1.0934 | 0.0168 | 3.3320 | 0.0008 |
| birth weight | 0.9809 | 0.9729 | 0.9889 | 0.0041 | -4.6309 | <0.0001 |
| preexisting hypertension | 1.0011 | 0.8090 | 1.2388 | 0.1086 | 0.0107 | 0.9913 |
| preexisting diabetes | 1.1478 | 1.0010 | 1.3161 | 0.0698 | 1.9752 | 0.0482 |
| previous induced abortion | 1.0374 | 1.0163 | 1.0590 | 0.0104 | 3.5102 | 0.0004 |
| previous spontaneous abortion | 1.0460 | 1.0231 | 1.0694 | 0.0112 | 3.9893 | <0.0001 |
| education level | 0.9591 | 0.9501 | 0.9682 | 0.0048 | -8.6570 | <0.0001 |
| parental income | 0.9855 | 0.9756 | 0.9955 | 0.0051 | -2.8342 | 0.0045 |
| country | 0.8454 | 0.8096 | 0.8828 | 0.0220 | -7.6009 | <0.0001 |
| region Sjælland | 0.9590 | 0.9346 | 0.9840 | 0.0131 | -3.1806 | 0.0014 |
| region Syddanmark | 1.0309 | 1.0082 | 1.0542 | 0.0113 | 2.6825 | 0.0073 |
| region Midtjylland | 0.8759 | 0.8560 | 0.8964 | 0.0117 | -11.2440 | <0.0001 |
| region Nordjylland | 0.9177 | 0.8918 | 0.9444 | 0.0146 | -5.8751 | <0.0001 |
| mother with disorder | 1.3052 | 1.2825 | 1.3284 | 0.0089 | 29.7029 | <0.0001 |
| father with disorder | 1.1841 | 1.1634 | 1.2051 | 0.0089 | 18.8522 | <0.0001 |

| 18. Endocrine - all | exp(coef) | LCL | UCL | SE | Z | Pr(Z) |
| --- | --- | --- | --- | --- | --- | --- |
| surgery | 1.2498 | 1.1615 | 1.3448 | 0.0373 | 5.9680 | <0.0001 |
| paternal age | 0.9493 | 0.9343 | 0.9646 | 0.0081 | -6.3778 | <0.0001 |
| maternal age | 0.8715 | 0.8556 | 0.8878 | 0.0094 | -14.5499 | <0.0001 |
| gestation length | 0.9888 | 0.9758 | 1.0020 | 0.0067 | -1.6617 | 0.0965 |
| maternal bleeding | 1.0630 | 1.0151 | 1.1131 | 0.0235 | 2.5993 | 0.0093 |
| fetal oxygen deprivation | 1.1572 | 0.9064 | 1.4773 | 0.1246 | 1.1720 | 0.2411 |
| pregnancy oedema | 1.2313 | 1.1321 | 1.3393 | 0.0428 | 4.8531 | <0.0001 |
| apgar5 score | 1.1654 | 1.1098 | 1.2237 | 0.0249 | 6.1377 | <0.0001 |
| birth weight | 1.0353 | 1.0227 | 1.0482 | 0.0062 | 5.5293 | <0.0001 |
| preexisting hypertension | 1.0635 | 0.7852 | 1.4404 | 0.1547 | 0.3979 | 0.6906 |
| preexisting diabetes | 1.5223 | 1.2944 | 1.7905 | 0.0827 | 5.0777 | <0.0001 |
| previous induced abortion | 1.0197 | 0.9881 | 1.0524 | 0.0160 | 1.2183 | 0.2230 |
| previous spontaneous abortion | 1.0518 | 1.0169 | 1.0878 | 0.0171 | 2.9394 | 0.0032 |
| education level | 0.8468 | 0.8349 | 0.8589 | 0.0072 | -23.0261 | <0.0001 |
| parental income | 0.8842 | 0.8709 | 0.8978 | 0.0077 | -15.8490 | <0.0001 |
| country | 0.8548 | 0.8056 | 0.9070 | 0.0302 | -5.1861 | <0.0001 |
| region Sjælland | 1.2519 | 1.2060 | 1.2995 | 0.0190 | 11.7993 | <0.0001 |
| region Syddanmark | 1.1694 | 1.1305 | 1.2095 | 0.0172 | 9.0799 | <0.0001 |
| region Midtjylland | 0.8271 | 0.7972 | 0.8580 | 0.0187 | -10.1180 | <0.0001 |
| region Nordjylland | 0.8678 | 0.8292 | 0.9083 | 0.0232 | -6.0970 | <0.0001 |
| mother with disorder | 1.6514 | 1.6044 | 1.6997 | 0.0147 | 34.1040 | <0.0001 |
| father with disorder | 1.5128 | 1.4625 | 1.5649 | 0.0172 | 23.9887 | <0.0001 |

Table S6
| 19. Endocrine - obesity |  |  |  |  |  |  |
| --- | --- | --- | --- | --- | --- | --- |
|  | exp(coef) | LCL | UCL | SE | Z | Pr(Z) |
| surgery | 1.1916 | 1.0885 | 1.3046 | 0.0462 | 3.7952 | 0.0001 |
| paternal age | 0.9285 | 0.9103 | 0.9471 | 0.0101 | -7.3198 | <0.0001 |
| maternal age | 0.8198 | 0.8009 | 0.8391 | 0.0118 | -16.6960 | <0.0001 |
| gestation length | 1.0050 | 0.9885 | 1.0218 | 0.0084 | 0.5936 | 0.5527 |
| maternal bleeding | 1.0765 | 1.0160 | 1.1406 | 0.0295 | 2.4979 | 0.0124 |
| fetal oxygen deprivation | 1.1821 | 0.8724 | 1.6019 | 0.1550 | 1.0795 | 0.2803 |
| pregnancy oedema | 1.2645 | 1.1456 | 1.3959 | 0.0504 | 4.6559 | <0.0001 |
| apgar5 score | 1.1945 | 1.1242 | 1.2693 | 0.0309 | 5.7399 | <0.0001 |
| birth weight | 1.0755 | 1.0590 | 1.0923 | 0.0079 | 9.2091 | <0.0001 |
| preexisting hypertension | 1.1374 | 0.7892 | 1.6392 | 0.1864 | 0.6905 | 0.4898 |
| preexisting diabetes | 1.6679 | 1.3602 | 2.0452 | 0.1040 | 4.9169 | <0.0001 |
| previous induced abortion | 1.0187 | 0.9790 | 1.0600 | 0.0202 | 0.9145 | 0.3604 |
| previous spontaneous abortion | 1.0662 | 1.0216 | 1.1128 | 0.0218 | 2.9422 | 0.0032 |
| education level | 0.7639 | 0.7503 | 0.7778 | 0.0092 | -29.2593 | <0.0001 |
| parental income | 0.8557 | 0.8394 | 0.8723 | 0.0098 | -15.8812 | <0.0001 |
| country | 0.6193 | 0.5712 | 0.6716 | 0.0413 | -11.5984 | <0.0001 |
| region Sjælland | 1.4857 | 1.4184 | 1.5563 | 0.0236 | 16.7269 | <0.0001 |
| region Syddanmark | 1.3026 | 1.2478 | 1.3599 | 0.0219 | 12.0420 | <0.0001 |
| region Midtjylland | 0.7980 | 0.7602 | 0.8377 | 0.0247 | -9.1105 | <0.0001 |
| region Nordjylland | 0.8534 | 0.8042 | 0.9055 | 0.0302 | -5.2406 | <0.0001 |
| mother with disorder | 2.3340 | 2.2258 | 2.4475 | 0.0242 | 34.9937 | <0.0001 |
| father with disorder | 1.9770 | 1.8367 | 2.1281 | 0.0375 | 18.1434 | <0.0001 |

| 20. Genitourinary - all |  |  |  |  |  |  |
| --- | --- | --- | --- | --- | --- | --- |
|  | exp(coef) | LCL | UCL | SE | Z | Pr(Z) |
| surgery | 1.1776 | 1.0853 | 1.2777 | 0.0416 | 3.9267 | <0.0001 |
| paternal age | 0.9672 | 0.9510 | 0.9836 | 0.0086 | -3.8722 | 0.0001 |
| maternal age | 0.8900 | 0.8726 | 0.9076 | 0.0100 | -11.6140 | <0.0001 |
| gestation length | 0.9816 | 0.9681 | 0.9953 | 0.0070 | -2.6129 | 0.0089 |
| maternal bleeding | 1.1251 | 1.0738 | 1.1789 | 0.0238 | 4.9503 | <0.0001 |
| fetal oxygen deprivation | 0.8135 | 0.5889 | 1.1239 | 0.1648 | -1.2512 | 0.2108 |
| pregnancy oedema | 1.1380 | 1.0275 | 1.2605 | 0.0521 | 2.4806 | 0.0131 |
| apgar5 score | 1.0480 | 0.9934 | 1.1056 | 0.0272 | 1.7202 | 0.0853 |
| birth weight | 0.9581 | 0.9456 | 0.9707 | 0.0066 | -6.4206 | <0.0001 |
| preexisting hypertension | 1.2001 | 0.8862 | 1.6252 | 0.1547 | 1.1793 | 0.2382 |
| preexisting diabetes | 1.1710 | 0.9452 | 1.4507 | 0.1092 | 1.4447 | 0.1485 |
| previous induced abortion | 1.1267 | 1.0917 | 1.1629 | 0.0161 | 7.4076 | <0.0001 |
| previous spontaneous abortion | 1.0440 | 1.0078 | 1.0815 | 0.0180 | 2.3915 | 0.0167 |
| education level | 0.9250 | 0.9113 | 0.9389 | 0.0076 | -10.2393 | <0.0001 |
| parental income | 0.9283 | 0.9138 | 0.9430 | 0.0080 | -9.2393 | <0.0001 |
| country | 0.8874 | 0.8347 | 0.9433 | 0.0312 | -3.8270 | 0.0001 |
| region Sjælland | 0.7068 | 0.6791 | 0.7357 | 0.0203 | -17.0072 | <0.0001 |
| region Syddanmark | 0.6676 | 0.6445 | 0.6915 | 0.0179 | -22.4944 | <0.0001 |
| region Midtjylland | 0.6189 | 0.5970 | 0.6415 | 0.0183 | -26.1738 | <0.0001 |
| region Nordjylland | 0.6194 | 0.5913 | 0.6488 | 0.0236 | -20.2316 | <0.0001 |
| mother with disorder | 1.3177 | 1.2758 | 1.3610 | 0.0164 | 16.7316 | <0.0001 |
| father with disorder | 1.2786 | 1.2218 | 1.3379 | 0.0231 | 10.6155 | <0.0001 |

Table S6
| 21. Genitourinary - kidney infection |  |  |  |  |  |  |
| --- | --- | --- | --- | --- | --- | --- |
|  | exp(coef) | LCL | UCL | SE | Z | Pr(Z) |
| surgery | 1.4194 | 1.1611 | 1.7350 | 0.1024 | 3.4185 | 0.0006 |
| paternal age | 0.9844 | 0.9431 | 1.0277 | 0.0219 | -0.7128 | 0.4759 |
| maternal age | 0.8572 | 0.8151 | 0.9014 | 0.0256 | -6.0013 | <0.0001 |
| gestation length | 1.0140 | 0.9792 | 1.0502 | 0.0178 | 0.7834 | 0.4333 |
| maternal bleeding | 1.1370 | 1.0100 | 1.2799 | 0.0604 | 2.1254 | 0.0335 |
| fetal oxygen deprivation | 0.7711 | 0.3202 | 1.8569 | 0.4483 | -0.5795 | 0.5622 |
| pregnancy oedema | 0.8451 | 0.6180 | 1.1555 | 0.1596 | -1.0540 | 0.2918 |
| apgar5 score | 1.1387 | 0.9981 | 1.2991 | 0.0672 | 1.9321 | 0.0533 |
| birth weight | 0.9874 | 0.9553 | 1.0206 | 0.0168 | -0.7468 | 0.4551 |
| preexisting hypertension | 0.5216 | 0.1679 | 1.6195 | 0.5780 | -1.1259 | 0.2602 |
| preexisting diabetes | 1.4261 | 0.8824 | 2.3047 | 0.2449 | 1.4493 | 0.1472 |
| previous induced abortion | 1.1702 | 1.0817 | 1.2659 | 0.0401 | 3.9185 | <0.0001 |
| previous spontaneous abortion | 1.1164 | 1.0232 | 1.2181 | 0.0444 | 2.4758 | 0.0132 |
| education level | 0.9354 | 0.9003 | 0.9719 | 0.0194 | -3.4217 | 0.0006 |
| parental income | 0.9560 | 0.9183 | 0.9953 | 0.0205 | -2.1859 | 0.0288 |
| country | 0.6327 | 0.5305 | 0.7546 | 0.0898 | -5.0920 | <0.0001 |
| region Sjælland | 0.6796 | 0.6139 | 0.7524 | 0.0519 | -7.4392 | <0.0001 |
| region Syddanmark | 0.6308 | 0.5764 | 0.6904 | 0.0460 | -10.0116 | <0.0001 |
| region Midtjylland | 0.6346 | 0.5801 | 0.6942 | 0.0457 | -9.9351 | <0.0001 |
| region Nordjylland | 0.6390 | 0.5696 | 0.7169 | 0.0586 | -7.6325 | <0.0001 |
| mother with disorder | 1.5065 | 1.2137 | 1.8699 | 0.1102 | 3.7173 | 0.0002 |
| father with disorder | 1.7034 | 1.1196 | 2.5916 | 0.2141 | 2.4877 | 0.0128 |

| 22. Musculoskeletal - all |  |  |  |  |  |  |
| --- | --- | --- | --- | --- | --- | --- |
|  | exp(coef) | LCL | UCL | SE | Z | Pr(Z) |
| surgery | 1.1507 | 1.1060 | 1.1972 | 0.0201 | 6.9515 | <0.0001 |
| paternal age | 0.9734 | 0.9655 | 0.9814 | 0.0041 | -6.4612 | <0.0001 |
| maternal age | 0.9366 | 0.9278 | 0.9454 | 0.0047 | -13.6665 | <0.0001 |
| gestation length | 1.0002 | 0.9937 | 1.0067 | 0.0033 | 0.0669 | 0.9465 |
| maternal bleeding | 1.0876 | 1.0634 | 1.1122 | 0.0114 | 7.3392 | <0.0001 |
| fetal oxygen deprivation | 1.1704 | 1.0404 | 1.3165 | 0.0600 | 2.6209 | 0.0087 |
| pregnancy oedema | 1.1067 | 1.0522 | 1.1641 | 0.0257 | 3.9332 | <0.0001 |
| apgar5 score | 1.0178 | 0.9925 | 1.0437 | 0.0128 | 1.3744 | 0.1693 |
| birth weight | 0.9898 | 0.9838 | 0.9959 | 0.0031 | -3.2653 | 0.0010 |
| preexisting hypertension | 1.0574 | 0.9087 | 1.2304 | 0.0773 | 0.7221 | 0.4701 |
| preexisting diabetes | 1.0131 | 0.9114 | 1.1261 | 0.0539 | 0.2420 | 0.8087 |
| previous induced abortion | 1.0479 | 1.0320 | 1.0640 | 0.0078 | 5.9985 | <0.0001 |
| previous spontaneous abortion | 1.0304 | 1.0135 | 1.0477 | 0.0084 | 3.5428 | 0.0003 |
| education level | 0.9552 | 0.9484 | 0.9620 | 0.0036 | -12.6205 | <0.0001 |
| parental income | 0.9932 | 0.9856 | 1.0008 | 0.0038 | -1.7512 | 0.0799 |
| country | 0.8558 | 0.8291 | 0.8834 | 0.0161 | -9.6201 | <0.0001 |
| region Sjælland | 0.8460 | 0.8293 | 0.8630 | 0.0101 | -16.4093 | <0.0001 |
| region Syddanmark | 0.8842 | 0.8692 | 0.8995 | 0.0087 | -14.0782 | <0.0001 |
| region Midtjylland | 1.0389 | 1.0220 | 1.0561 | 0.0083 | 4.5636 | <0.0001 |
| region Nordjylland | 0.8327 | 0.8144 | 0.8513 | 0.0112 | -16.2125 | <0.0001 |
| mother with disorder | 1.3412 | 1.3250 | 1.3576 | 0.0061 | 47.3784 | <0.0001 |
| father with disorder | 1.2297 | 1.2147 | 1.2449 | 0.0062 | 33.0616 | <0.0001 |

Table S6
| 23. Neoplasms - all |  |  |  |  |  |  |
| --- | --- | --- | --- | --- | --- | --- |
|  | exp(coef) | LCL | UCL | SE | Z | Pr(Z) |
| surgery | 1.1965 | 1.1090 | 1.2910 | 0.0387 | 4.6301 | <0.0001 |
| paternal age | 0.9826 | 0.9669 | 0.9986 | 0.0082 | -2.1247 | 0.0336 |
| maternal age | 0.9713 | 0.9537 | 0.9893 | 0.0093 | -3.0998 | 0.0019 |
| gestation length | 0.9992 | 0.9866 | 1.0120 | 0.0064 | -0.1093 | 0.9129 |
| maternal bleeding | 1.0699 | 1.0235 | 1.1183 | 0.0225 | 2.9919 | 0.0027 |
| fetal oxygen deprivation | 1.1125 | 0.8807 | 1.4053 | 0.1191 | 0.8948 | 0.3708 |
| pregnancy oedema | 1.0002 | 0.9063 | 1.1039 | 0.0503 | 0.0050 | 0.9960 |
| apgar5 score | 0.9849 | 0.9368 | 1.0355 | 0.0255 | -0.5920 | 0.5538 |
| birth weight | 1.0192 | 1.0071 | 1.0314 | 0.0060 | 3.1394 | 0.0016 |
| preexisting hypertension | 0.9166 | 0.6637 | 1.2658 | 0.1646 | -0.5285 | 0.5970 |
| preexisting diabetes | 1.0152 | 0.8217 | 1.2542 | 0.1078 | 0.1399 | 0.8887 |
| previous induced abortion | 0.9909 | 0.9612 | 1.0216 | 0.0155 | -0.5825 | 0.5601 |
| previous spontaneous abortion | 1.0214 | 0.9889 | 1.0550 | 0.0165 | 1.2880 | 0.1977 |
| education level | 1.0070 | 0.9932 | 1.0210 | 0.0070 | 1.0015 | 0.3165 |
| parental income | 1.0274 | 1.0124 | 1.0426 | 0.0074 | 3.6129 | 0.0003 |
| country | 0.9162 | 0.8565 | 0.9801 | 0.0343 | -2.5445 | 0.0109 |
| region Sjælland | 1.1096 | 1.0683 | 1.1525 | 0.0193 | 5.3800 | <0.0001 |
| region Syddanmark | 1.1874 | 1.1488 | 1.2273 | 0.0168 | 10.1777 | <0.0001 |
| region Midtjylland | 1.1417 | 1.1044 | 1.1802 | 0.0169 | 7.8273 | <0.0001 |
| region Nordjylland | 0.9370 | 0.8967 | 0.9790 | 0.0224 | -2.9026 | 0.0037 |
| mother with disorder | 1.1846 | 1.1558 | 1.2140 | 0.0125 | 13.5036 | <0.0001 |
| father with disorder | 1.1538 | 1.1164 | 1.1924 | 0.0167 | 8.5200 | <0.0001 |

| 24. Neoplasms - benign |  |  |  |  |  |  |
| --- | --- | --- | --- | --- | --- | --- |
|  | exp(coef) | LCL | UCL | SE | Z | Pr(Z) |
| surgery | 1.1925 | 1.0976 | 1.2956 | 0.0423 | 4.1633 | <0.0001 |
| paternal age | 0.9773 | 0.9605 | 0.9944 | 0.0088 | -2.5897 | 0.0096 |
| maternal age | 0.9775 | 0.9584 | 0.9970 | 0.0100 | -2.2489 | 0.0245 |
| gestation length | 0.9976 | 0.9841 | 1.0113 | 0.0069 | -0.3386 | 0.7348 |
| maternal bleeding | 1.0580 | 1.0086 | 1.1098 | 0.0243 | 2.3147 | 0.0206 |
| fetal oxygen deprivation | 1.1542 | 0.8989 | 1.4820 | 0.1275 | 1.1243 | 0.2608 |
| pregnancy oedema | 1.0003 | 0.8981 | 1.1141 | 0.0549 | 0.0062 | 0.9950 |
| apgar5 score | 0.9801 | 0.9286 | 1.0343 | 0.0274 | -0.7304 | 0.4651 |
| birth weight | 1.0163 | 1.0034 | 1.0295 | 0.0065 | 2.4868 | 0.0128 |
| preexisting hypertension | 0.8470 | 0.5918 | 1.2121 | 0.1828 | -0.9080 | 0.3638 |
| preexisting diabetes | 1.0284 | 0.8214 | 1.2876 | 0.1146 | 0.2447 | 0.8066 |
| previous induced abortion | 0.9972 | 0.9651 | 1.0303 | 0.0166 | -0.1656 | 0.8683 |
| previous spontaneous abortion | 1.0420 | 1.0066 | 1.0787 | 0.0176 | 2.3338 | 0.0196 |
| education level | 1.0121 | 0.9971 | 1.0273 | 0.0075 | 1.5853 | 0.1128 |
| parental income | 1.0249 | 1.0088 | 1.0413 | 0.0080 | 3.0484 | 0.0023 |
| country | 0.9226 | 0.8588 | 0.9912 | 0.0365 | -2.2005 | 0.0277 |
| region Sjælland | 1.0989 | 1.0548 | 1.1449 | 0.0208 | 4.5164 | <0.0001 |
| region Syddanmark | 1.2012 | 1.1592 | 1.2447 | 0.0181 | 10.1049 | <0.0001 |
| region Midtjylland | 1.1515 | 1.1112 | 1.1933 | 0.0181 | 7.7536 | <0.0001 |
| region Nordjylland | 0.9003 | 0.8581 | 0.9445 | 0.0244 | -4.2881 | <0.0001 |
| mother with disorder | 1.2137 | 1.1791 | 1.2494 | 0.0147 | 13.1172 | <0.0001 |
| father with disorder | 1.1925 | 1.1420 | 1.2453 | 0.0220 | 7.9773 | <0.0001 |

Table S6
| 25. Circulatory - all | exp(coef) | LCL | UCL | SE | Z | Pr(Z) |
| --- | --- | --- | --- | --- | --- | --- |
| surgery | 1.1121 | 0.9969 | 1.2407 | 0.0558 | 1.9046 | 0.0568 |
| paternal age | 0.9728 | 0.9501 | 0.9961 | 0.0120 | -2.2772 | 0.0227 |
| maternal age | 0.9462 | 0.9211 | 0.9721 | 0.0137 | -4.0185 | <0.0001 |
| gestation length | 0.9690 | 0.9509 | 0.9874 | 0.0095 | -3.2755 | 0.0010 |
| maternal bleeding | 1.1394 | 1.0697 | 1.2136 | 0.0322 | 4.0526 | <0.0001 |
| fetal oxygen deprivation | 0.8315 | 0.5654 | 1.2229 | 0.1968 | -0.9371 | 0.3486 |
| pregnancy oedema | 1.0120 | 0.8785 | 1.1658 | 0.0721 | 0.1661 | 0.8680 |
| apgar5 score | 1.2349 | 1.1544 | 1.3209 | 0.0343 | 6.1392 | <0.0001 |
| birth weight | 0.9760 | 0.9589 | 0.9934 | 0.0090 | -2.6886 | 0.0071 |
| preexisting hypertension | 1.1761 | 0.8152 | 1.6970 | 0.1870 | 0.8676 | 0.3855 |
| preexisting diabetes | 0.9493 | 0.6974 | 1.2921 | 0.1573 | -0.3307 | 0.7408 |
| previous induced abortion | 0.9829 | 0.9393 | 1.0285 | 0.0231 | -0.7436 | 0.4571 |
| previous spontaneous abortion | 1.0404 | 0.9917 | 1.0916 | 0.0244 | 1.6201 | 0.1052 |
| education level | 0.9741 | 0.9544 | 0.9942 | 0.0104 | -2.5081 | 0.0121 |
| parental income | 0.9792 | 0.9582 | 1.0007 | 0.0110 | -1.8948 | 0.0581 |
| country | 0.8924 | 0.8121 | 0.9806 | 0.0481 | -2.3651 | 0.0180 |
| region Sjælland | 0.9897 | 0.9370 | 1.0454 | 0.0279 | -0.3682 | 0.7126 |
| region Syddanmark | 0.9163 | 0.8723 | 0.9625 | 0.0251 | -3.4798 | 0.0005 |
| region Midtjylland | 0.9554 | 0.9101 | 1.0029 | 0.0247 | -1.8413 | 0.0655 |
| region Nordjylland | 0.7704 | 0.7214 | 0.8227 | 0.0334 | -7.7861 | <0.0001 |
| mother with disorder | 1.4909 | 1.4270 | 1.5576 | 0.0223 | 17.8648 | <0.0001 |
| father with disorder | 1.3222 | 1.2689 | 1.3777 | 0.0209 | 13.3130 | <0.0001 |

| 26. Nervous System - all | exp(coef) | LCL | UCL | SE | Z | Pr(Z) |
| --- | --- | --- | --- | --- | --- | --- |
| surgery | 1.3524 | 1.1959 | 1.5293 | 0.0627 | 4.8112 | <0.0001 |
| paternal age | 0.9804 | 0.9521 | 1.0096 | 0.0149 | -1.3191 | 0.1871 |
| maternal age | 0.8900 | 0.8603 | 0.9208 | 0.0173 | -6.7132 | <0.0001 |
| gestation length | 0.9920 | 0.9687 | 1.0160 | 0.0121 | -0.6543 | 0.5129 |
| maternal bleeding | 1.1314 | 1.0441 | 1.2260 | 0.0409 | 3.0149 | 0.0025 |
| fetal oxygen deprivation | 1.5102 | 1.0471 | 2.1782 | 0.1868 | 2.2065 | 0.0273 |
| pregnancy oedema | 1.1832 | 1.0098 | 1.3865 | 0.0808 | 2.0807 | 0.0374 |
| apgar5 score | 1.0732 | 0.9799 | 1.1755 | 0.0464 | 1.5231 | 0.1277 |
| birth weight | 0.9768 | 0.9553 | 0.9988 | 0.0113 | -2.0651 | 0.0389 |
| preexisting hypertension | 1.0825 | 0.6139 | 1.9085 | 0.2893 | 0.2740 | 0.7840 |
| preexisting diabetes | 1.4650 | 1.0485 | 2.0469 | 0.1706 | 2.2380 | 0.0252 |
| previous induced abortion | 1.0664 | 1.0082 | 1.1279 | 0.0286 | 2.2463 | 0.0246 |
| previous spontaneous abortion | 1.0933 | 1.0300 | 1.1606 | 0.0304 | 2.9303 | 0.0033 |
| education level | 0.9016 | 0.8787 | 0.9251 | 0.0131 | -7.8894 | <0.0001 |
| parental income | 0.9399 | 0.9145 | 0.9661 | 0.0139 | -4.4255 | <0.0001 |
| country | 0.6862 | 0.6027 | 0.7813 | 0.0661 | -5.6882 | <0.0001 |
| region Sjælland | 1.3932 | 1.3035 | 1.4890 | 0.0339 | 9.7673 | <0.0001 |
| region Syddanmark | 1.1524 | 1.0828 | 1.2265 | 0.0317 | 4.4642 | <0.0001 |
| region Midtjylland | 0.9780 | 0.9166 | 1.0434 | 0.0330 | -0.6726 | 0.5011 |
| region Nordjylland | 0.7574 | 0.6935 | 0.8272 | 0.0449 | -6.1784 | <0.0001 |
| mother with disorder | 1.5697 | 1.4651 | 1.6818 | 0.0351 | 12.8151 | <0.0001 |
| father with disorder | 1.4063 | 1.2917 | 1.5311 | 0.0433 | 7.8650 | <0.0001 |

Table S6
| 27. Eye/adnexa - all | exp(coef) | LCL | UCL | SE | Z | Pr(Z) |
| --- | --- | --- | --- | --- | --- | --- |
| surgery | 1.3439 | 1.2057 | 1.4980 | 0.0553 | 5.3374 | <0.0001 |
| paternal age | 1.0036 | 0.9883 | 1.0191 | 0.0078 | 0.4623 | 0.6438 |
| maternal age | 0.9761 | 0.9591 | 0.9935 | 0.0089 | -2.6837 | 0.0072 |
| gestation length | 0.9557 | 0.9439 | 0.9677 | 0.0063 | -7.1251 | <0.0001 |
| maternal bleeding | 1.0895 | 1.0440 | 1.1370 | 0.0217 | 3.9370 | <0.0001 |
| fetal oxygen deprivation | 1.1580 | 0.9055 | 1.4810 | 0.1255 | 1.1691 | 0.2423 |
| pregnancy oedema | 1.0570 | 0.9579 | 1.1664 | 0.0502 | 1.1039 | 0.2696 |
| apgar5 score | 1.0668 | 1.0179 | 1.1179 | 0.0239 | 2.7062 | 0.0068 |
| birth weight | 0.9580 | 0.9469 | 0.9693 | 0.0059 | -7.1712 | <0.0001 |
| preexisting hypertension | 1.0174 | 0.7591 | 1.3635 | 0.1493 | 0.1157 | 0.9078 |
| preexisting diabetes | 1.0961 | 0.9015 | 1.3329 | 0.0997 | 0.9206 | 0.3572 |
| previous induced abortion | 1.0355 | 1.0057 | 1.0662 | 0.0149 | 2.3417 | 0.0191 |
| previous spontaneous abortion | 1.0649 | 1.0320 | 1.0988 | 0.0159 | 3.9337 | <0.0001 |
| education level | 0.9582 | 0.9454 | 0.9711 | 0.0068 | -6.2411 | <0.0001 |
| parental income | 0.9567 | 0.9431 | 0.9704 | 0.0072 | -6.0702 | <0.0001 |
| country | 1.0333 | 0.9785 | 1.0913 | 0.0278 | 1.1797 | 0.2381 |
| region Sjælland | 0.7686 | 0.7414 | 0.7968 | 0.0183 | -14.3026 | <0.0001 |
| region Syddanmark | 0.7535 | 0.7302 | 0.7775 | 0.0160 | -17.6485 | <0.0001 |
| region Midtjylland | 0.6397 | 0.6192 | 0.6610 | 0.0166 | -26.8004 | <0.0001 |
| region Nordjylland | 0.6694 | 0.6419 | 0.6980 | 0.0213 | -18.7850 | <0.0001 |
| mother with disorder | 1.3313 | 1.2831 | 1.3814 | 0.0188 | 15.1965 | <0.0001 |
| father with disorder | 1.2361 | 1.1901 | 1.2838 | 0.0193 | 10.9628 | <0.0001 |

| 28. Mental - all | exp(coef) | LCL | UCL | SE | Z | Pr(Z) |
| --- | --- | --- | --- | --- | --- | --- |
| surgery | 1.1956 | 1.1306 | 1.2644 | 0.0285 | 6.2604 | <0.0001 |
| paternal age | 1.0122 | 1.0007 | 1.0239 | 0.0058 | 2.0859 | 0.0369 |
| maternal age | 0.9184 | 0.9061 | 0.9308 | 0.0068 | -12.3935 | <0.0001 |
| gestation length | 0.9811 | 0.9717 | 0.9906 | 0.0049 | -3.8751 | 0.0001 |
| maternal bleeding | 1.1328 | 1.0971 | 1.1696 | 0.0163 | 7.6405 | <0.0001 |
| fetal oxygen deprivation | 1.1972 | 0.9948 | 1.4407 | 0.0944 | 1.9052 | 0.0567 |
| pregnancy oedema | 1.0729 | 0.9953 | 1.1566 | 0.0383 | 1.8385 | 0.0659 |
| apgar5 score | 1.0600 | 1.0219 | 1.0995 | 0.0186 | 3.1251 | 0.0017 |
| birth weight | 0.9718 | 0.9630 | 0.9807 | 0.0046 | -6.1335 | <0.0001 |
| preexisting hypertension | 1.1690 | 0.9421 | 1.4505 | 0.1100 | 1.4188 | 0.1559 |
| preexisting diabetes | 1.0875 | 0.9378 | 1.2611 | 0.0755 | 1.1106 | 0.2667 |
| previous induced abortion | 1.1292 | 1.1049 | 1.1541 | 0.0111 | 10.9336 | <0.0001 |
| previous spontaneous abortion | 1.0701 | 1.0443 | 1.0965 | 0.0124 | 5.4413 | <0.0001 |
| education level | 0.9607 | 0.9505 | 0.9710 | 0.0054 | -7.3597 | <0.0001 |
| parental income | 0.7863 | 0.7774 | 0.7953 | 0.0058 | -41.1905 | <0.0001 |
| country | 0.4847 | 0.4597 | 0.5111 | 0.0270 | -26.7723 | <0.0001 |
| region Sjælland | 0.8470 | 0.8234 | 0.8713 | 0.0144 | -11.5155 | <0.0001 |
| region Syddanmark | 0.9173 | 0.8954 | 0.9396 | 0.0122 | -7.0261 | <0.0001 |
| region Midtjylland | 0.7687 | 0.7493 | 0.7885 | 0.0129 | -20.2398 | <0.0001 |
| region Nordjylland | 0.5602 | 0.5402 | 0.5811 | 0.0186 | -31.1175 | <0.0001 |
| mother with disorder | 2.0378 | 1.9897 | 2.0871 | 0.0121 | 58.4125 | <0.0001 |
| father with disorder | 1.6307 | 1.5853 | 1.6775 | 0.0144 | 33.9188 | <0.0001 |

**Table S7.**
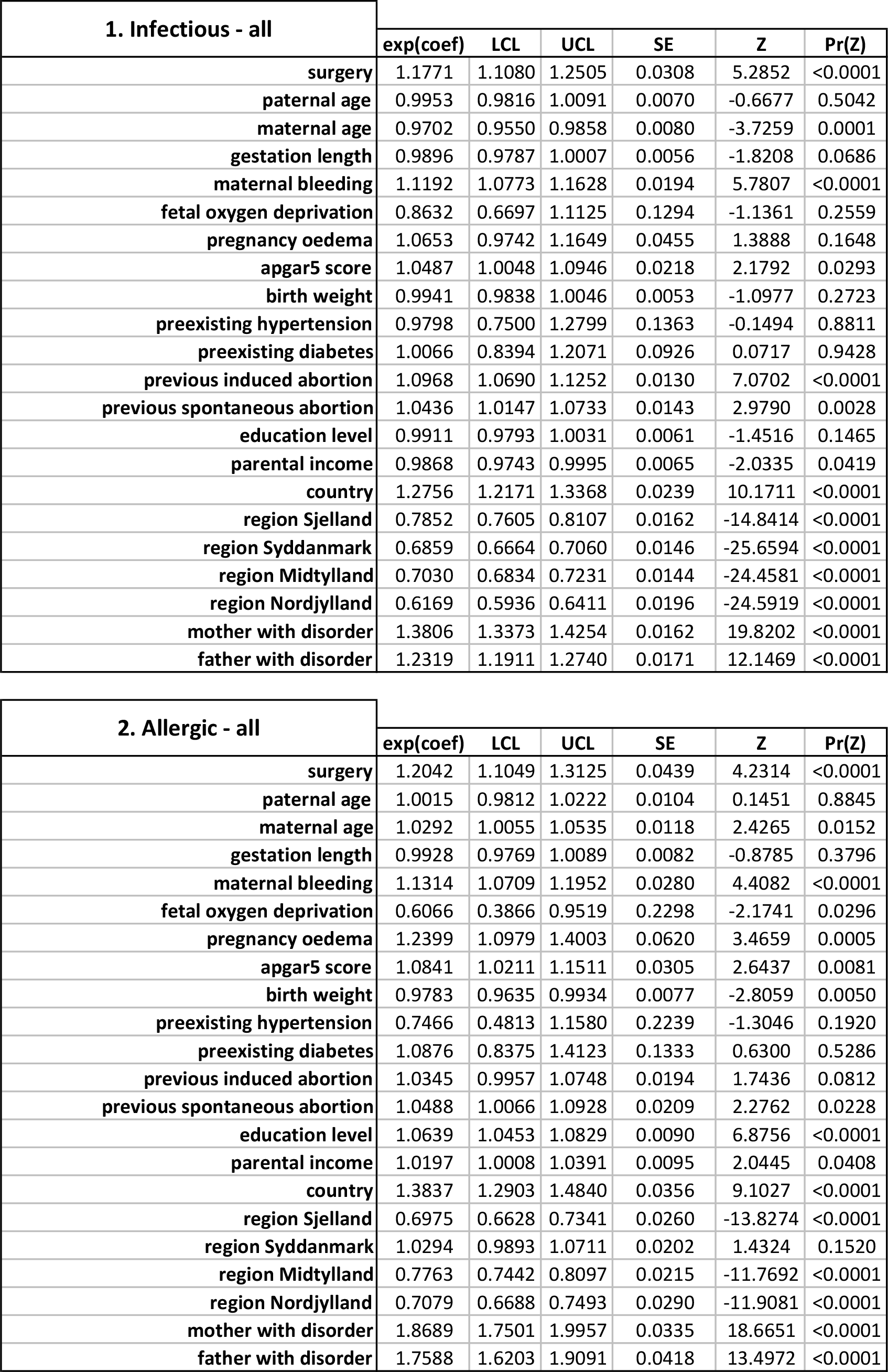
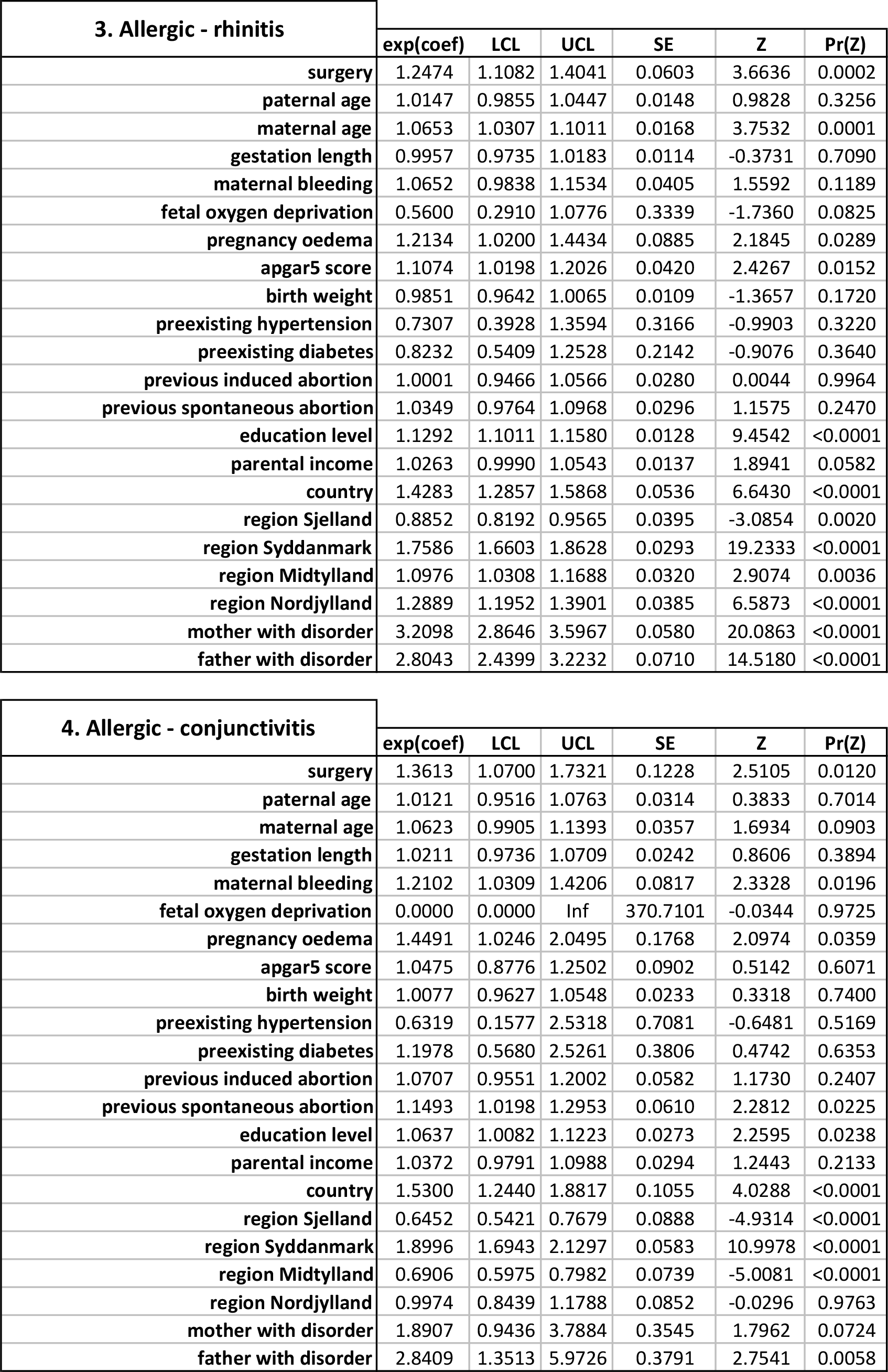
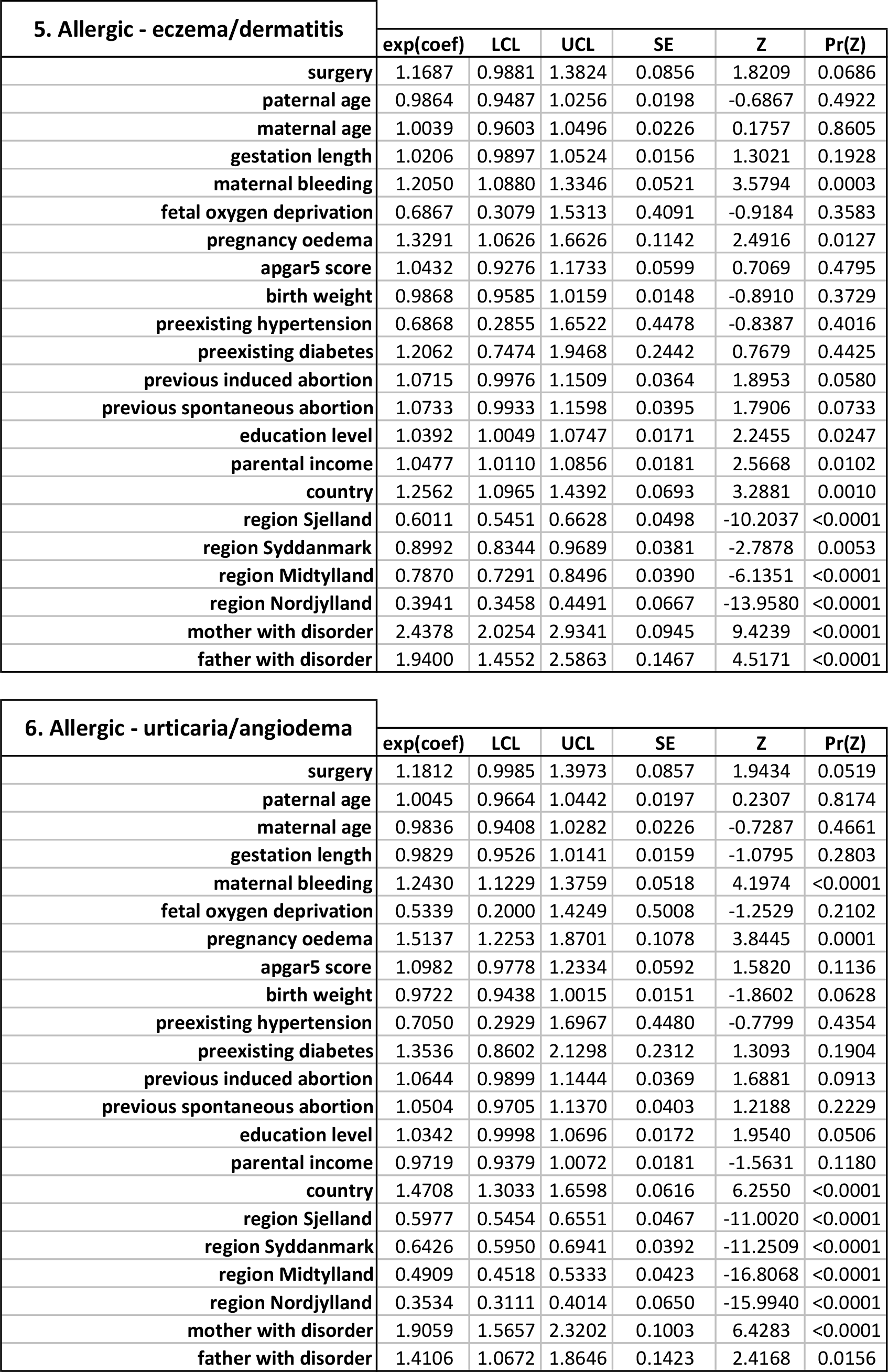
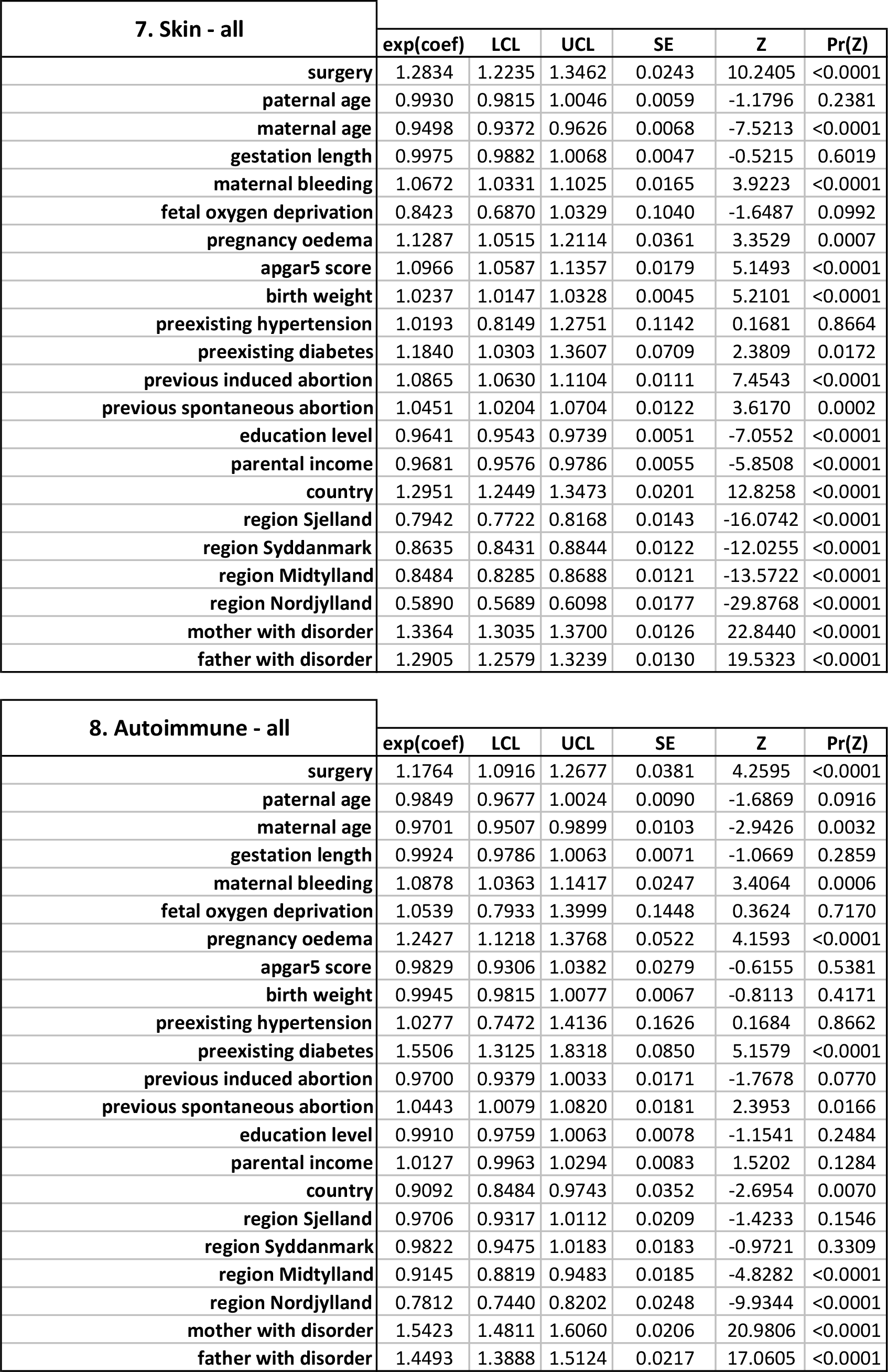
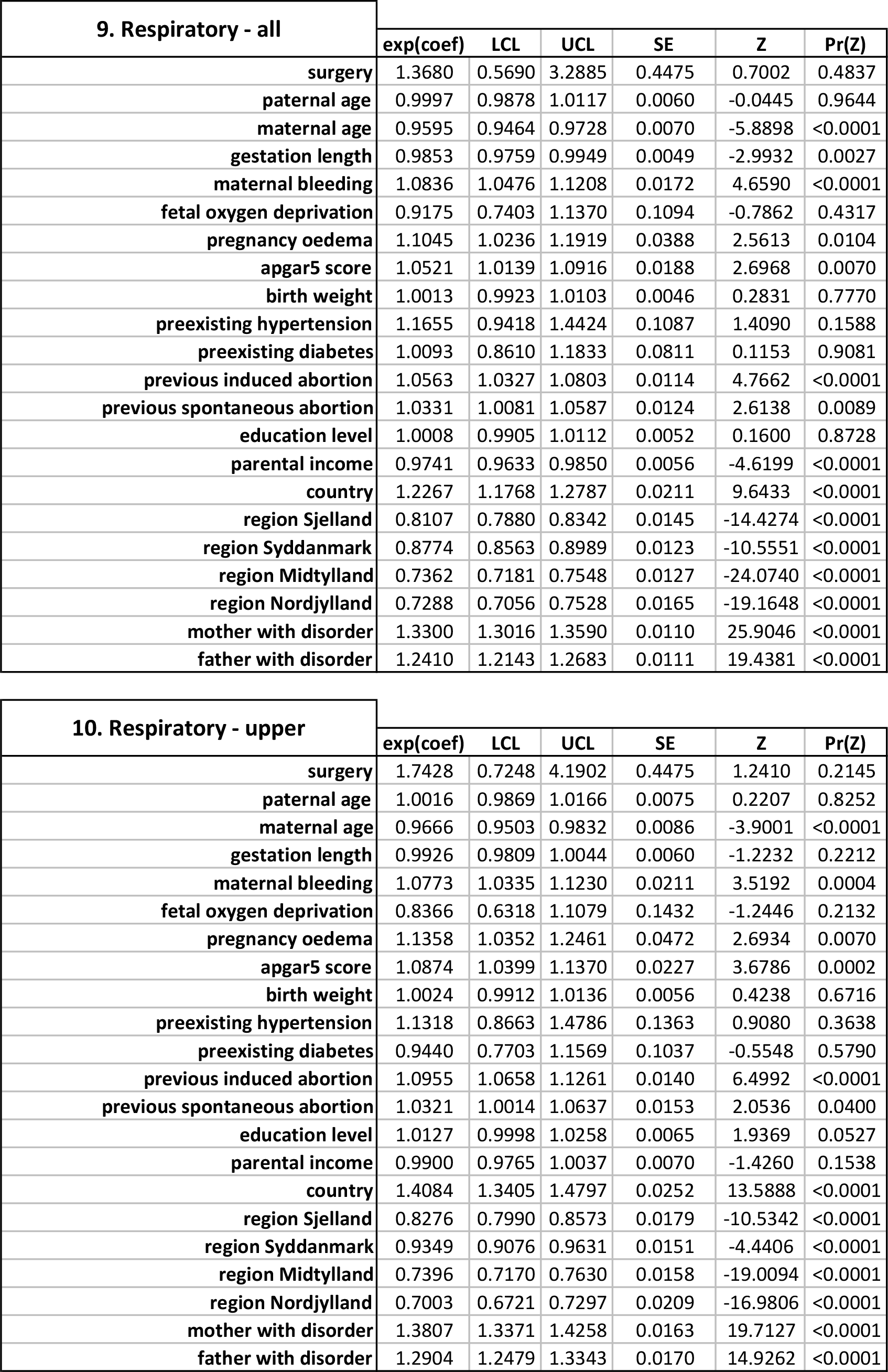
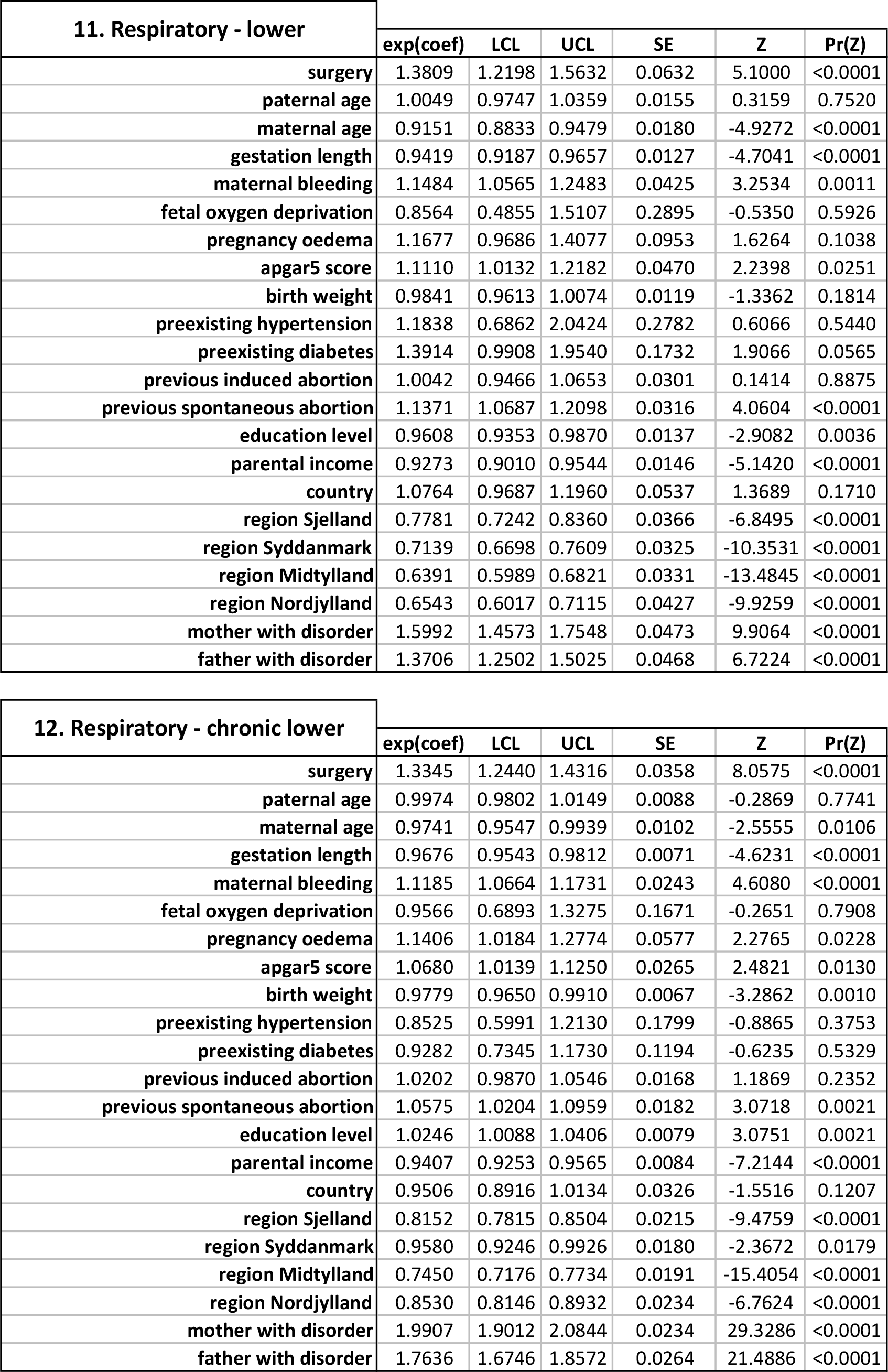
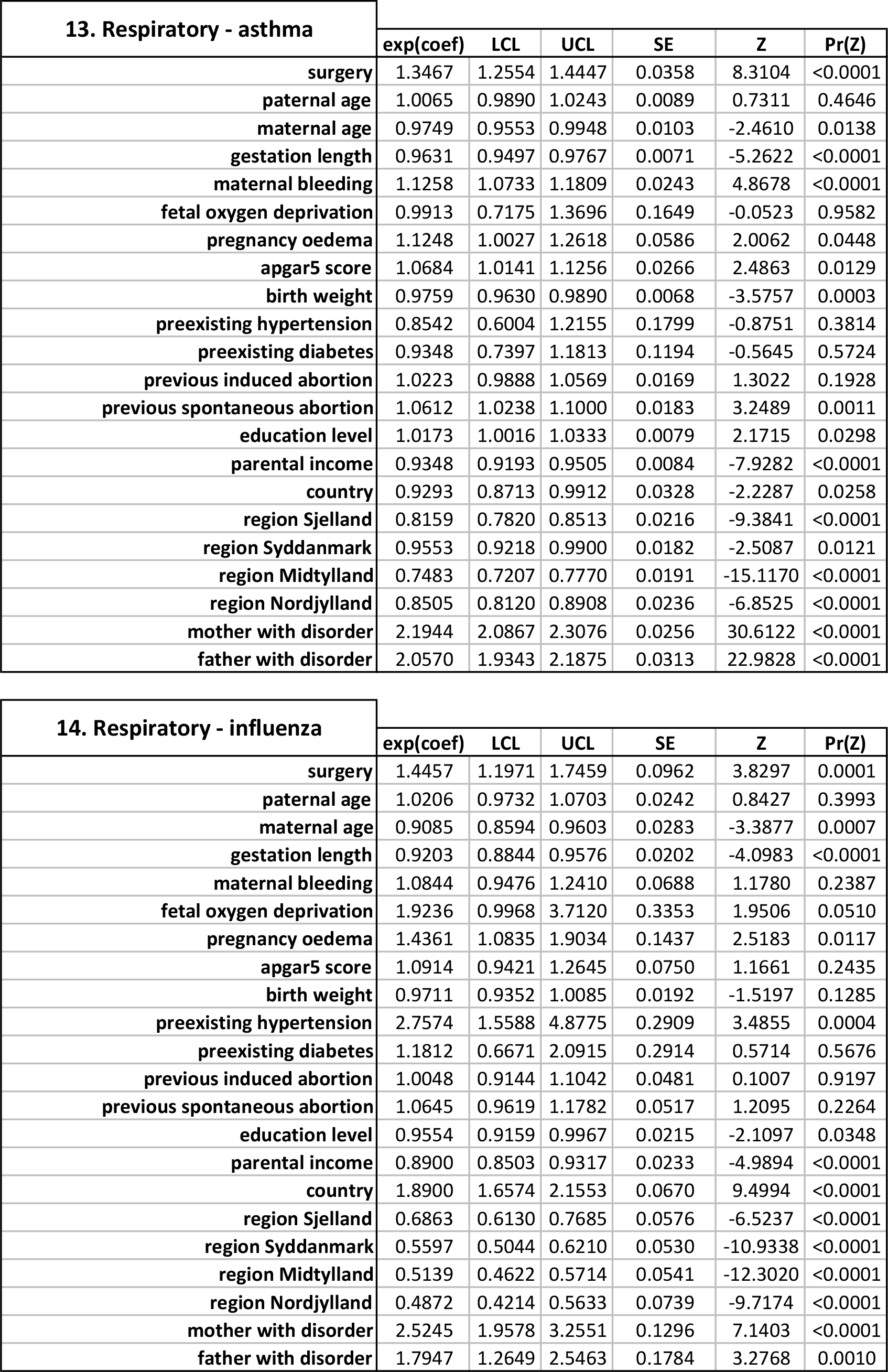
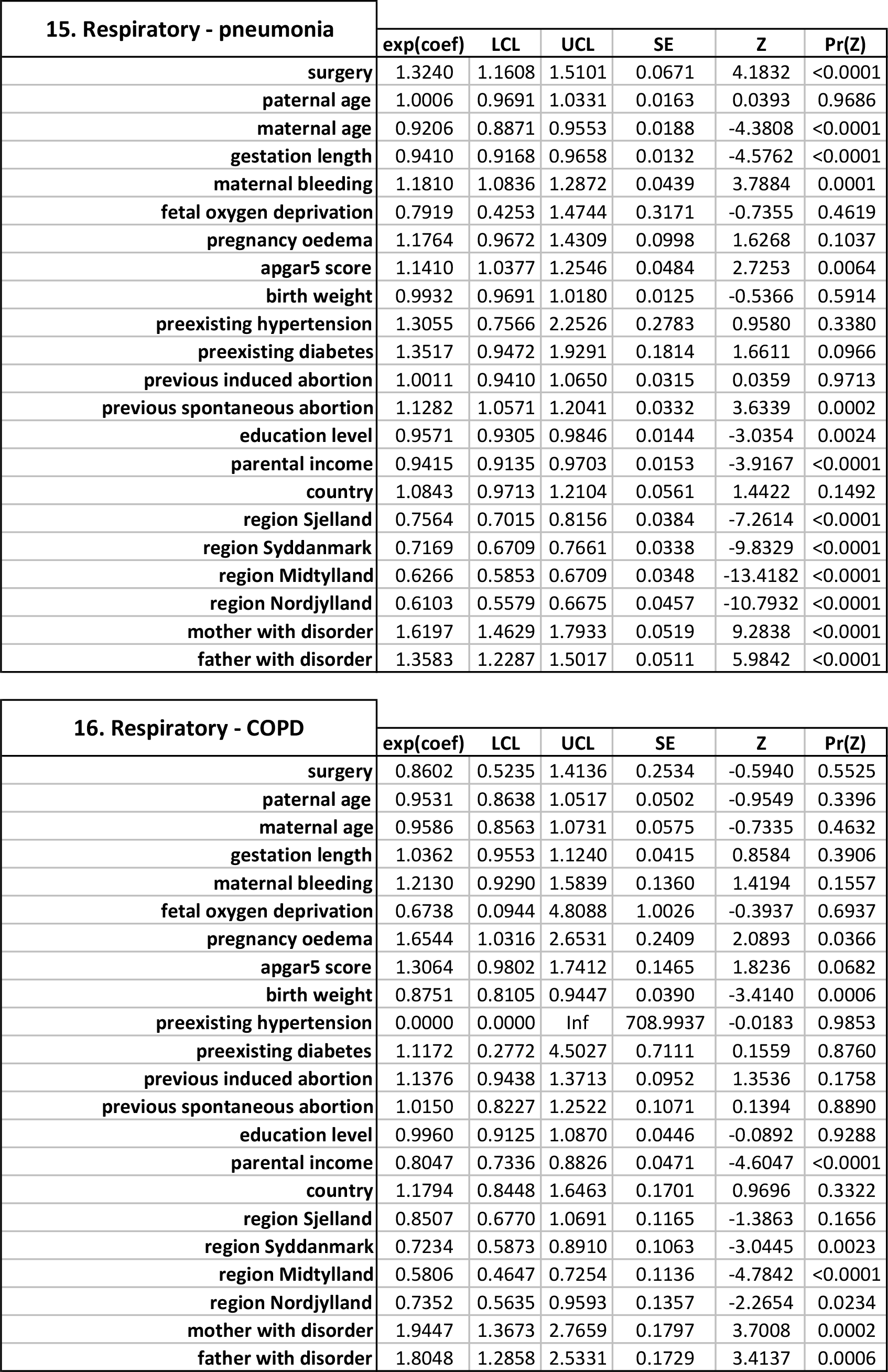
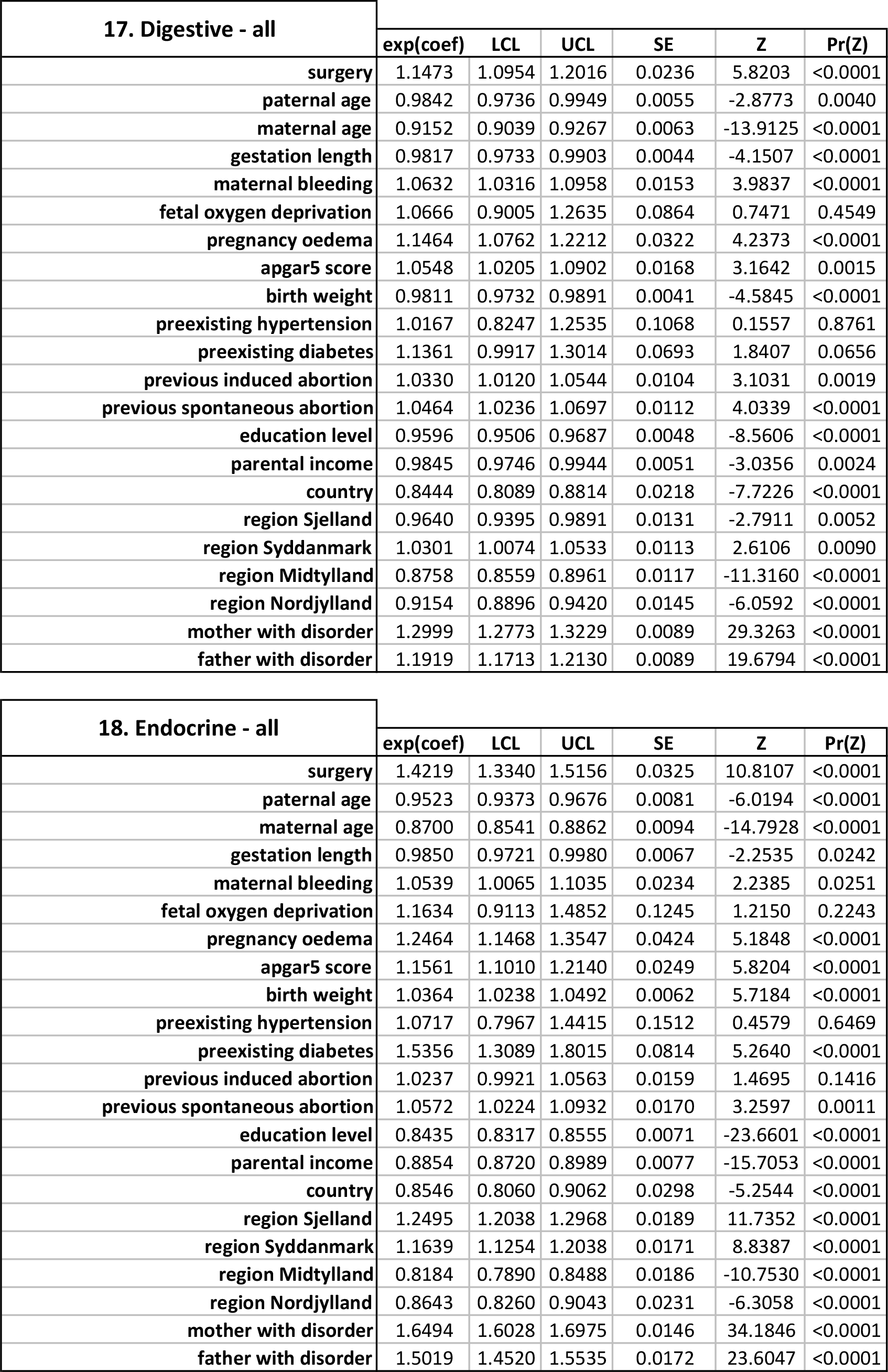
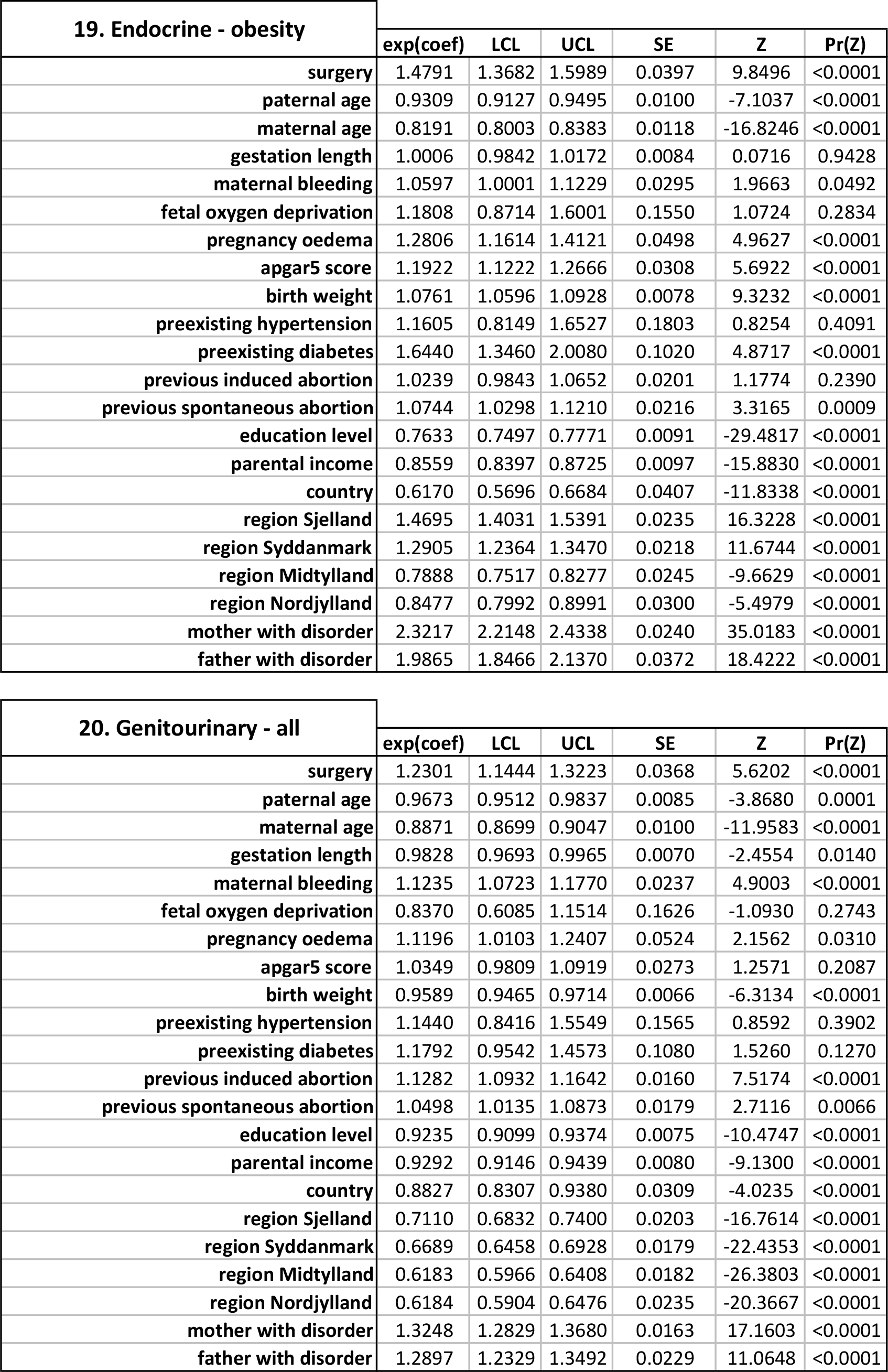
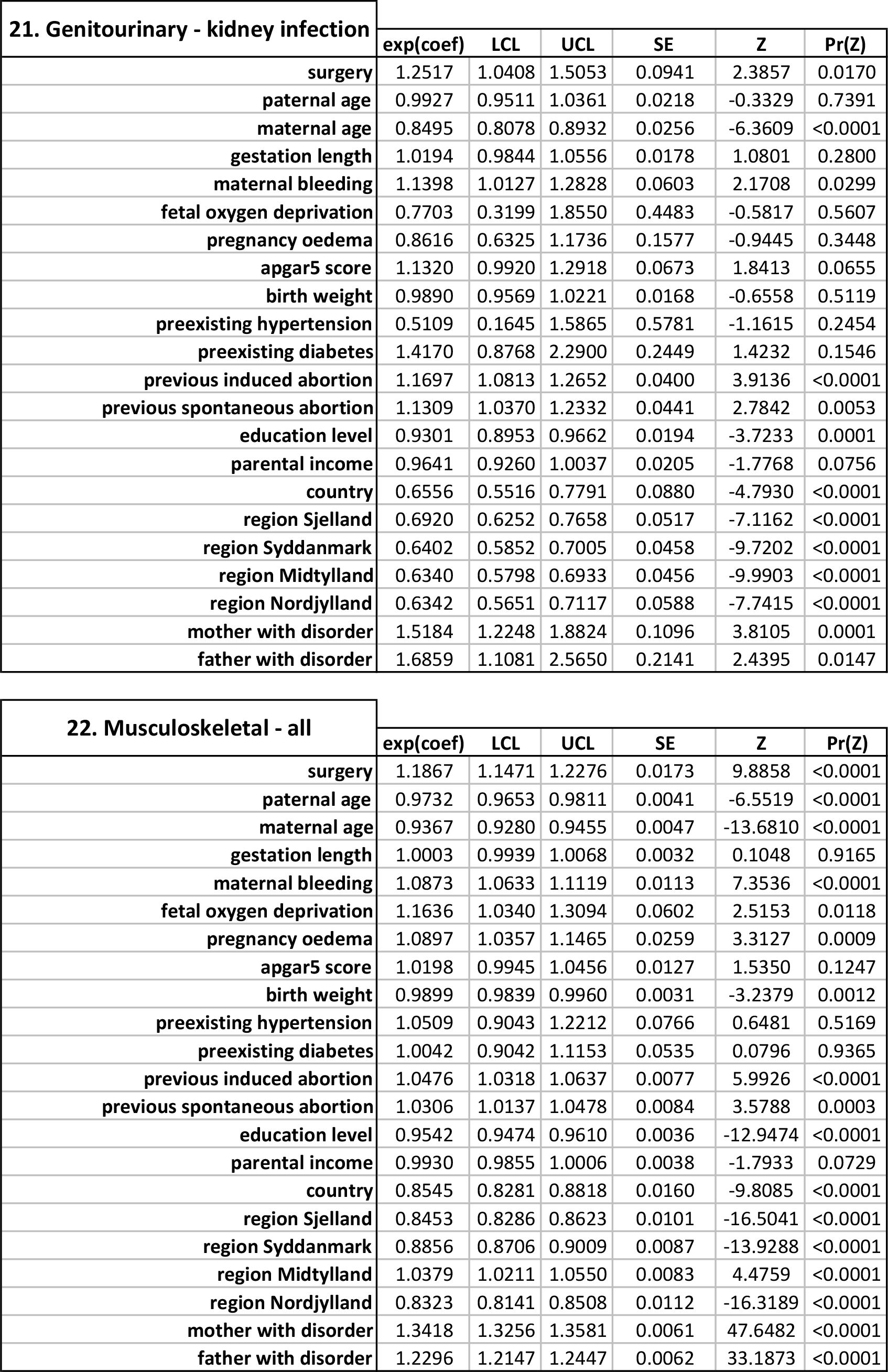
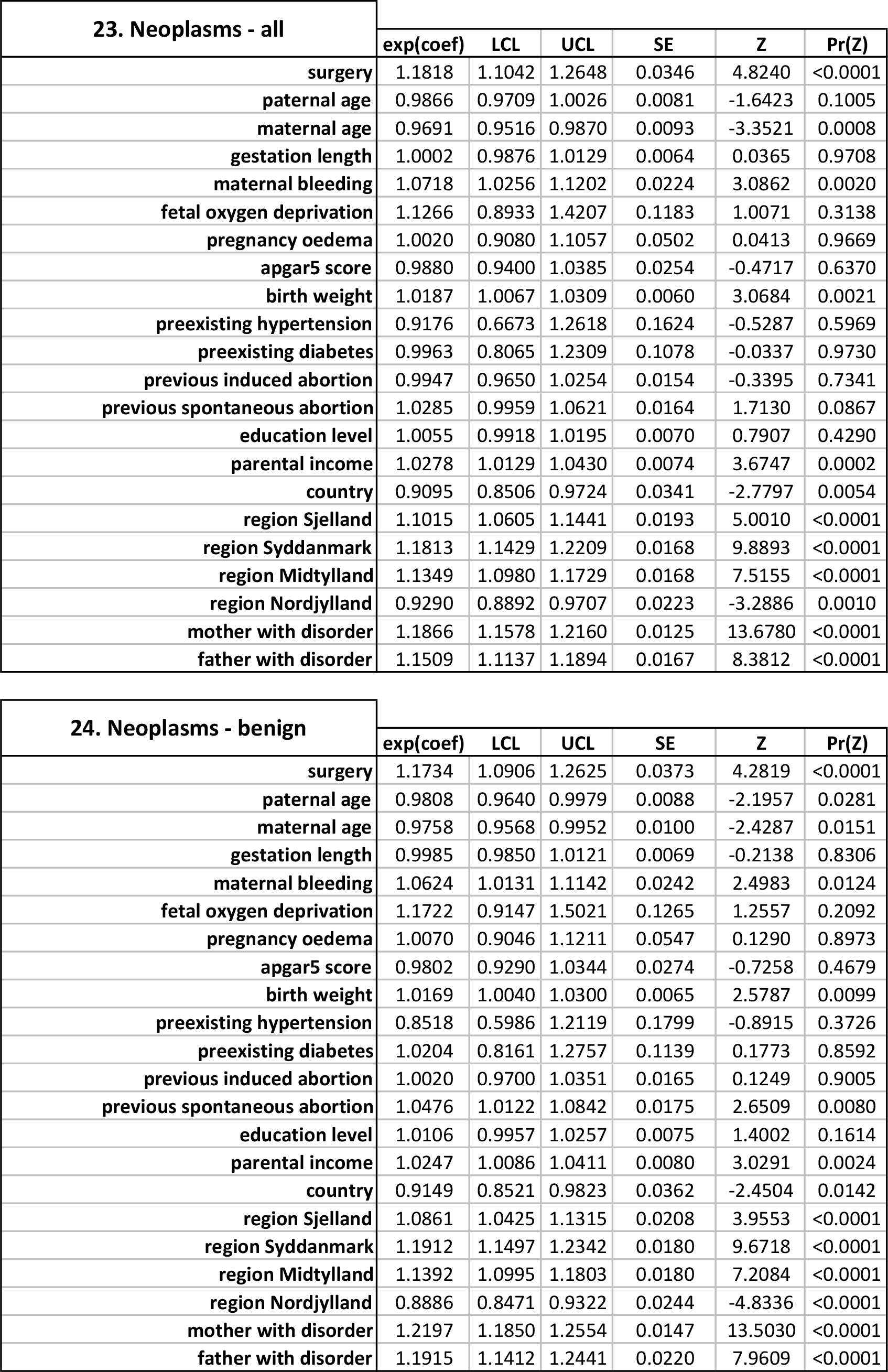
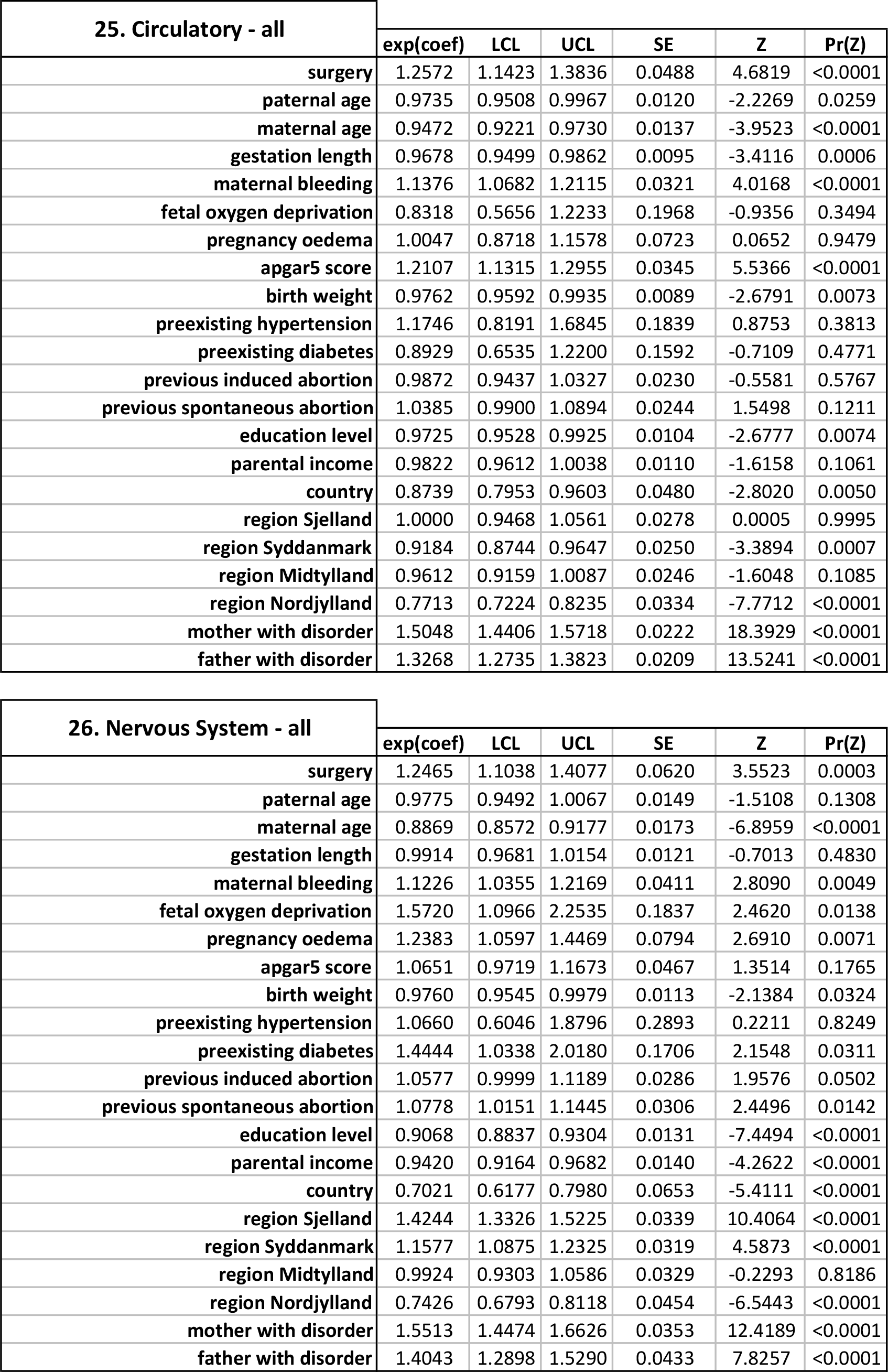
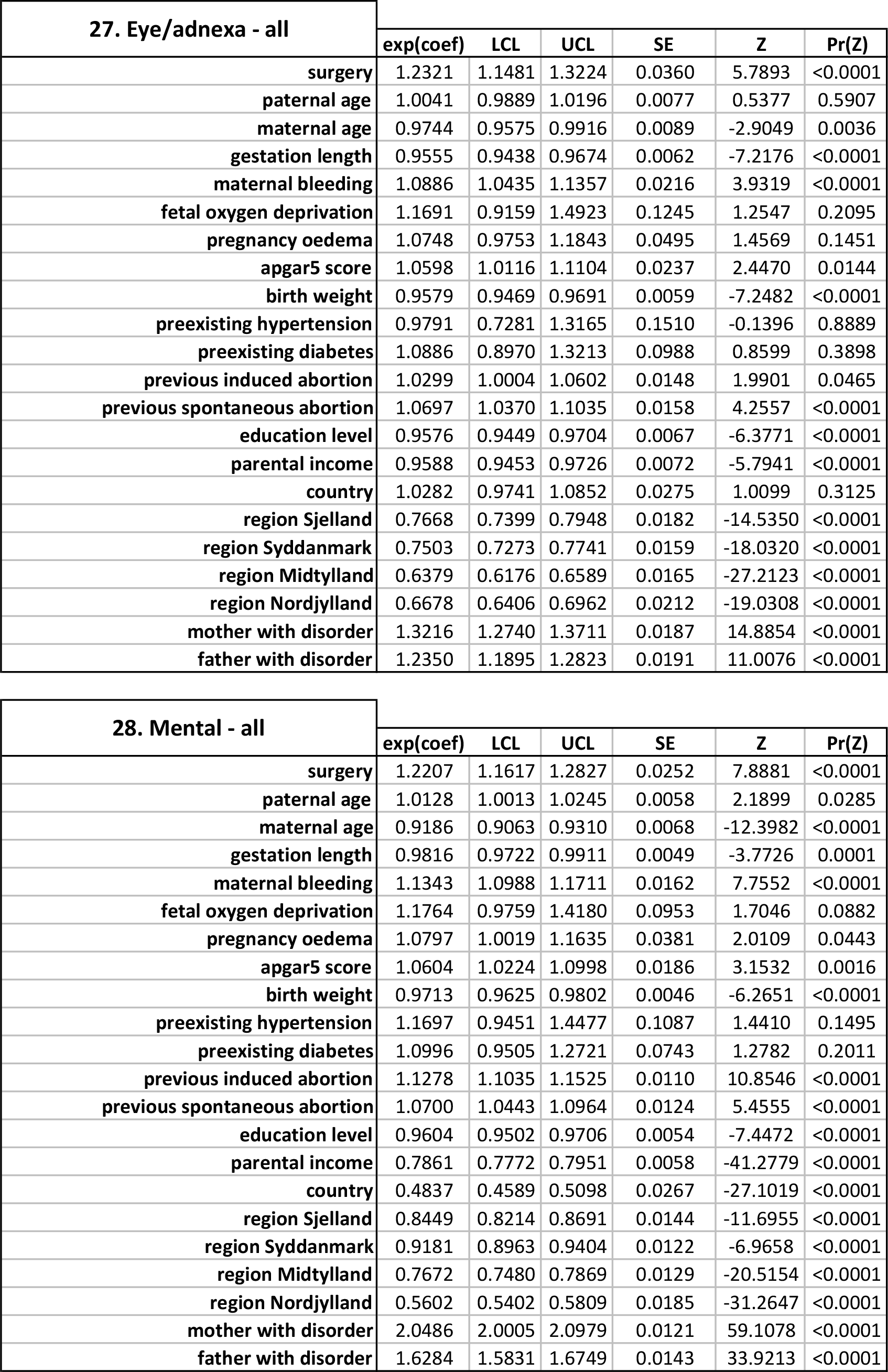
**Full Cox regression outputs - impact of adenotonsillectomy on risk of later disease.**

**Table S8.** **Relative risks (RR), absolute risk difference (ARD), number needed to treat (NNT(B/H) or benefit/harm) for conditions surgeries aim to treat**

| Conditions | adenoidectomy |  |  |  | tonsillectomy |  |  |  | adenotonsillectomy |  |  |  | CR |
| --- | --- | --- | --- | --- | --- | --- | --- | --- | --- | --- | --- | --- | --- |
|  | RR | ER | ARD | NNT(B/H) | RR | ER | ARD | NNT(B/H) | RR | ER | ARD | NNT(B/H) |  |
| Sleep disorders | 0.30(0.15-0.60)* | 1.05 | -0.083 % | -1209 (B) | 0.24(0.07-0.75) | - | - | - | 0.46(0.28-0.77) | - | - | - | 0.119 |
| Abnormal breathing | 1.04(0.61-1.78) | - | - | - | 0.70(0.26-1.89) | - | - | - | 0.59(0.30-1.14) | - | - | - | 0.045 |
| Sinusitis | 1.54(1.14-2.08) | - | - | - | 1.03(0.63-1.69) | - | - | - | 1.68(1.32-2.14)*** | 0.479 | +0.116% | 865 (H) | 0.169 |
| Chronic sinusitis | 2.25(1.69-3.00)*** | 1.47 | +0.11% | 905 (H) | 1.25(0.75-2.08) | - | - | - | 1.17(0.84-1.63) | - | - | - | 0.088 |
| Labyrinthitis | 4.76(1.82-12.40)* | 10.21 | +0.171% | 586 (H) | - | - | - | - | 3.27(0.99-10.74) | - | - | - | 0.045 |
| Otitis media | 4.84(4.48-5.23)*** | 57.75 | +19.495% | 5 (H) | 2.22(1.94-2.54)*** | 10.263 | +6.217% | 16 (H) | 2.06(1.88-2.25)*** | 27.513 | +5.385% | 19 (H) | 5.073 |
| Tonsillitis | 0.21(0.16-0.29)*** | 5.86 | -1.815 % | -55 (B) | 0.09(0.05-0.17)*** | 16.487 | -2.102 % | -48 (B) | 0.09(0.06-0.14)*** | 14.685 | -2.092 % | -48 (B) | 2.321 |
| Chronic tonsillitis | 0.54(0.39-0.74)** | 92.36 | -0.299 % | -334 (B) | 0.20(0.11-0.38)*** | 80.486 | -0.52 % | -192 (B) | 0.11(0.06-0.19)*** | 98.315 | -0.582 % | -172 (B) | 0.657 |

- CR - Control Risk (i.e. event rate (%) in the control group) - ER - Experimental Risk (i.e. event rate (%) in the case (surgery) group) - ARD - Absolute Risk Difference (a.k.a. absolute risk reduction/increase) [ARD=100 X CR X (1-RR)] - NNT(B/H) - Number Needed to Treat-Benefit or Harm [NNT=100/ARD] - RR exclusion: Relative risks (RR) are presented only for analyses with sufficient power for hypothesis testing (see methods) - > i.e. there was insufficient power for testing labyrinthitis for the tonsillectomy analysis and insufficient power to test obstructive sleep apnea in any analysis (hence, there are no results here) - ER, ARD and NNT exclusion: these values are presented only when Bonferroni corrected p-value (23 tests; $\alpha=0.05/23=0.00217$ ) for relative risks (RR) were significant

## Supplementary Methods

### Study sample

While we had access to health data on individuals older than 30 years, we focused on individuals born and living between 1978o 2009 for two important reasons. Firstly, we needed to code whether surgery was performed for adenoidectomy, tonsillectomy or adenotonsillectomy between birth and 9 years of age. For any individual born before 1978 (who had no record of these surgeries after 1978), we could not determine that. Secondly, for individuals born before 1978, we had no prior disease history or information on most of their covariates. Therefore, including those individuals would have introduced both bias in censoring in the Cox regression model between those born before or after 1978 as well as introducing large amounts of missing information for those born before 1978 (i.e. they would have had to be removed before analysis anyway). Electronic health data were not available before 1977-78 in Denmark, and data after 2009 were not available at the time of request.

### Power analyses

To reduce the possibility of encountering false negative results, we carried out power analyses with the powerSurvEpi package in R v 3.2.1, which checks whether sample sizes are large enough to test the null hypothesis of no association between predictor and subsequent disease. The powerSurvEpi algorithm calculates the minimum sample size needed to detect an effect of a given size with a given degree of confidence based on the risk ratio, which we derived from the actual relative risks for focal diseases, the variance of the main covariate of interest (surgery), the proportion of subjects who were diagnosed with the disease, the postulated power, and the type I error rate. Most diseases within each surgery group met these thresholds. Of the 84 disease x surgery combinations examined (28 disease groups x 3 surgery groups = 84), 6 (̴7%) did not and were excluded from the results reported.

### Converting RR to AR and NNT

To interpret risks based on their actual effect size in the population [1] and provide statistics that are more clinically useful [2], we used risks derived from Cox regressions and prevalence of disease in control samples (control risk, CR) to estimate absolute risk difference (ARD, also known as absolute risk reduction or increase) and number needed to treat (NNT). ARD=100 x CR x (1-RR) and NNT=100/ARD. Conversion between RR and ARD is mainly modified by the control risk, i.e.if RR for a disease is large but its prevalence in the population is low, it will generally have a small ARD and thus low impact. Conversion between ARD and NNT implies they are inversely linked, i.e. if ARD is large, very few patients need to be treated to observe benefit or harm in one of them. If ARD is small, many patients need to be treated to observe benefit or harm in one of them. Positive and negative NNT values are used to indicate whether treatment results in harm (NNT-harm) or benefit (NNT-benefit) respectively, with numbers closer to zero indicating harm or benefit more often than larger values. For example, an NNT-benefit value of 5 suggests that treatment in 5 patients would need to be performed for one of them to benefit. An NNT-harm value of 100 suggests that treatment in 100 patients would need to be performed for one of them to experience an adverse effect.

### Testing for biases in early general health between cases and controls

In order to test whether there were significant differences between (surgery) cases and controls regarding diseases diagnosed in the first 9 years of life, i.e. testing whether individuals who underwent surgery in the first 9 years of life represented a generally ‘more sickly’ sample of individuals that could bias estimation of incident disease risk beyond 9 years of age, we tested two null hypotheses. *Null hypothesis 1:* There is no difference between case and control age distributions for diseases diagnosed in the first 9 years of life. This sought to compare age at diagnosis distributions for any disease (potentially multiple diagnoses for each individual) to see whether individuals who underwent the three focal surgeries also tended to have more disease diagnoses overall in the first 9 years of life. *Null hypothesis 2:* There is no difference in age at first disease diagnosed in the first 9 years of life between cases and controls. This sought to compare age at diagnosis distributions for the first disease diagnosed to see whether individuals who underwent the three focal surgeries received their first disease diagnosis earlier in life within the first 9 years. General health: In order to test for differences in general health, we pooled ICD diagnoses that capture almost all of the main disease groups described by ICD10, i.e. the same broad set defined in Hollegaard et al [3]. These capture a much wider range of diseases and should be more representative of general health in a population.

*For null hypotheis 1,* we randomly sampled from the control individuals (those who did not undergo surgery during 1978-2009) the same number of individuals who had undergone each specific surgery. For example, there were 20,238 (out of 1,668,632) individuals who underwent adenoidectomy between birth and 9 years of age in our sample. We randomly sampled the same number (n=20,238) from the control sample (n=1,648,394 without surgery) and tested whether distributions for general health diseases diagnosed at any age within the first 9 years between surgery individuals and controls were significantly different with a Kolmogorov-Smirnov test. We repeated this 1000 times, each time randomly sampling 20,238 individuals (with replacement) from the control sample and comparing this to the surgery group’s distribution for age at any general health disease diagnosis within the first 9 years. The permuted p value was based on the number of times (out of 1000) that the two distributions were the same. Permuted p was >0.05 in all cases supporting the null hypothesis no difference between case and control age distributions for general health diseases diagnosed in the first 9 years of life. *For null hypothesis 2,* the same permutation procedure was carried out as for null hypothesis 1, except only the age at first diagnosis was considered. For all three surgery groups, permuted p was > 0.05, supporting the null hypothesis. These results show that general health in the first 9 years of life was not significantly different between the surgery groups versus control individuals. In other words, individuals who underwent these surgeries in the first 9 years of life were not sicklier prior to surgery, compared to the general population.

There are two further pieces of information that help us be more confident here. Firstly, medical records for Danish residents are (unusually compared to other countries) very complete given that 1) the Danish health system is free for all residents so there is no need for residents to worry about going to the doctor whenever they need to, meaning there is also no difference or bias in health care access based on socioeconomic status and 2) clinicians are paid per diagnosis, so it is in their interest to rigorously document all diagnosis and health problems seen in their patients, which means that the health information this study is based on is likely to reflect the complete medical history of individuals included.

## Supplementary Results

### Consistent impact of covariates on risk of many diseases

Some covariates showed very consistent effects on risk for many diseases. Accounting for these effects is therefore likely to be important in any longitudinal population;based study examining these types of disease risks.

The most consistent effect in Fig. 2 in terms of disease relative risk magnitude (e.g. RR ranged from 1.10 to 3.71) and direction of effect (i.e. risk was increased for every disease) was whether either parent had previously been diagnosed within the same disease group (i.e. had received an ICD code for the same disease within the hospital system). The largest relative risks were found for asthma (RR ranged from 2.05 to 2.19 across all analyses), allergies (RR ranged from 1.10 to 3.26), obesity (RR ranged from 1.97 to 2.34), and mental disorders (RR ranged from 1.60 to 2.04): a family history of these diseases greatly increased risk of these in their children. As would be expected for polygenic diseases, these effects were largely consistent regardless of whether that disease was diagnosed on the maternal or paternal side. The exceptions were urticaria/angiodema, influenza, and kidney infections, where risk was significantly increased only if the mother (and not father) had a history of these diseases.

Maternal age at birth of individuals in our sample was consistently significantly associated with decreased risk for 16 of the 28 disease groups in Fig. 2. This effect was largest in endocrine (RR 0.79 to 0.87 across all analyses), genitourinary (RR 0.84 to 0.89), and nervous system (RR 0.87 to 0.89) diseases. The exception was allergic rhinitis, where older mothers conferred increased risk (RR 1.05 to 1.07). Maternal bleeding during pregnancy had many widespread and consistent effects, with 16 out of 28 diseases examined showing significantly elevated risks if this complication occurred (RR 1.02 to 1.27).

Other covariates such as parental education, income and country of origin also showed many significant effects, but risk direction varied considerably depending on the disease considered. For example, mental disorders were less frequent in Danish nationals than in immigrants (RR 0.48 to 0.49), yet risk of influenza was much higher in Danes (RR 1.89 to 2.06). Higher parental education and income mostly decreased the risk of many diseases (RR 0.76 to 0.99), but increased the risk of a few, including some allergic diseases (RR 1.04;1.06). Both effects could result from differences in the microbiota children are exposed to. Many complex patterns highlighted regional differences in diseases. For example, obesity risk was higher in Sjelland (RR 1.46 to 1.48) and Syddanmark (RR 1.29 to 1.39) but lower in Midtylland (RR 0.78 to 0.84) and Nordjylland (RR 0.84 to 0.92) relative to Hovedstaden.

### Other important and interesting associations of covariates with disease risk

That risk for several diseases was sensitive to many covariates suggests that these diseases may have particularly complex causation. For example, risk of skin (group: all), digestive (group: all) endocrine (group: all, obesity) and mental diseases (group: all) were influenced by many (up to 15 out of 21) covariates. There were also very specific disease risk; covariate interactions. For example, the relative risk of influenza (RR 2.75 to 3.12) (but no other disease) was significantly and massively increased by maternal hypertension prior to pregnancy; oedema during pregnancy greatly increased relative risk forurticaria/angiodema (RR 1.51 to 1.56); and diabetes prior to pregnancy greatly increased risk for all endocrine (RR 1.44 to 1.66) and autoimmune diseases (RR 1.47 to 1.55).

